# Combining case-control status and family history of disease increases association power

**DOI:** 10.1101/722645

**Authors:** Margaux L.A. Hujoel, Steven Gazal, Po-Ru Loh, Nick Patterson, Alkes L. Price

**Affiliations:** Department of Biostatistics, Harvard T.H. Chan School of Public Health, Boston, MA; Department of Epidemiology, Harvard T.H. Chan School of Public Health, Boston, MA; Broad Institute of MIT and Harvard, Cambridge, MA; Brigham and Women’s Hospital/Harvard Medical School, Boston, MA

## Abstract

Family history of disease can provide valuable information about an individual’s genetic liability for disease in case-control association studies, but it is currently unclear how to best combine case-control status and family history of disease. We developed a new association method based on posterior mean genetic liabilities under a liability threshold model, conditional on both case-control status and family history (LT-FH); association statistics are computed via linear regression of genotypes and posterior mean genetic liabilities, equivalent to a score test. We applied LT-FH to 12 diseases from the UK Biobank (average N=350K). We compared LT-FH to genome-wide association without using family history (GWAS) and a previous proxy-based method for incorporating family history (GWAX). LT-FH was +63% (s.e. 6%) more powerful than GWAS and +36% (s.e. 4%) more powerful than the trait-specific maximum of GWAS and GWAX, based on the number of independent genome-wide significant loci detected across all diseases (e.g. 690 independent loci for LT-FH vs. 423 for GWAS); the second best method was GWAX for lower-prevalence diseases and GWAS for higher-prevalence diseases, consistent with simulations. We also confirmed that LT-FH was well-calibrated (assessed via stratified LD score regression attenuation ratio), consistent with simulations. When using BOLT-LMM (instead of linear regression) to compute association statistics for all three methods (increasing the power of each method), LT-FH was +67% (s.e. 6%) more powerful than GWAS and +39% (s.e. 4%) more powerful than the trait-specific maximum of GWAS and GWAX. In summary, LT-FH greatly increases association power in case-control association studies when family history of disease is available.

## Introduction

Family history of disease can provide valuable information about an individual’s genetic liability for disease, potentially increasing the power of case-control association studies^1^, the focus of this study. More broadly, leveraging data from ungenotyped but phenotyped relatives has a rich history in genetic risk prediction^2,3^ and analyses of quantitative traits in humans^4^ and livestock^5–9^. Standard case-control genome-wide association studies (GWAS) ignore family history information (Fig 1a). A recent method, genome-wide association by proxy^1^ (GWAX), compares disease cases and “proxy cases” (controls with family history of disease) to controls without family history of disease (Fig 1a). This approach has proven successful in studies of Alzheimer’s disease^10–12^, and has provided a valuable contribution in highlighting the value of family history information, but more accurate modeling of family history — for example, distinguishing disease cases from proxy cases — may be beneficial^1^.

**Figure 1:**
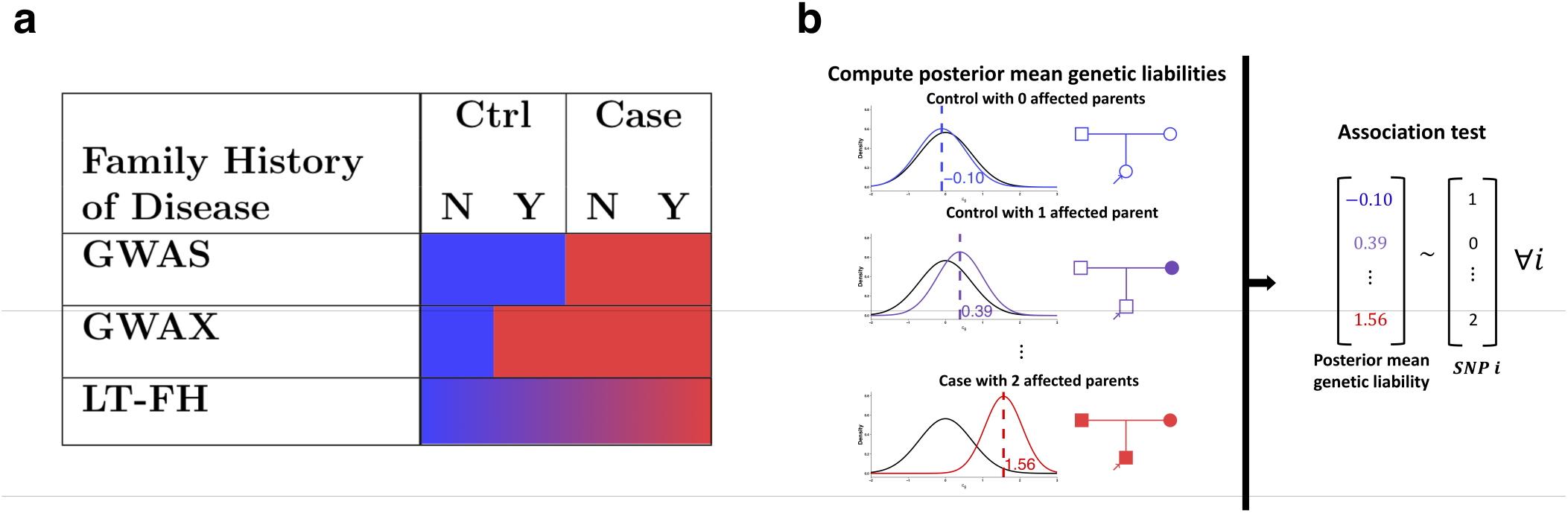
Overview of LT-FH and other methods. **(a)** GWAS uses binary case-control status, ignoring family history; GWAX uses binary proxy-case-control status, merging controls with family history of disease with disease cases; LT-FH uses continuous-valued posterior mean genetic liability, appropriately differentiating all case-control and family history configurations. **(b)** LT-FH computes posterior mean genetic liabilities (left panel) and then tests for association between genotype and posterior mean genetic liability (right panel).

We propose a new association method based on posterior mean genetic liabilities under a liability threshold model, conditional on both case-control status and family history (LT-FH); the liability threshold model, in which an individual is a disease case if and only if an underlying continuous-valued liability lies above a threshold, has proven valuable in a wide range of settings^2,3,13–19^. LT-FH computes association statistics via linear regression of genotypes and posterior mean genetic liabilities; association statistics can also be computed using efficient mixed-model methods^20,21^. LT-FH captures the fact that controls with family history have higher genetic liability than controls without family history (and likewise for disease cases) (Fig 1a). In contrast to GWAX, which assigns the same binary phenotype to disease cases and proxy cases, LT-FH accurately models a broad range of case-control status and family history configurations, greatly increasing power for diseases of both low and high prevalence in analyses of 12 diseases from the UK Biobank.

## Results

### Overview of methods

The LT-FH method relies on the liability threshold model^13^, and consists of two main steps (Fig 1b): (1) compute posterior mean genetic liabilities for each genotyped individual, conditional on available case-control status and/or family history information; (2) compute association statistics via linear regression of genotypes and posterior mean genetic liabilities. In step 1, we first compute posterior mean genetic liabilities for every possible configuration of case-control status and family history (377 configurations, accounting for parental history, sibling history, missing data, etc.) and then perform a lookup for each genotyped individual, avoiding redundant computation. The posterior mean genetic liability for each configuration is computed via Monte Carlo integration, incorporating estimates of disease prevalence and parental disease prevalence from the target samples and estimates of narrow-sense heritability^22^ (which differs from SNP-heritability^23^) from the literature; we show that Monte Carlo integration outperforms an analytical approach (the Pearson-Aitken(PA) formula^2,24,25^) in analyses of sibling history data.

Step 2 generalizes the Armitage trend test^26^, and we show that it is equivalent to a score test; an alternative is to apply BOLT-LMM^20,21^ (see URLs) to posterior mean genetic liabilities, increasing power in large samples. We emphasize that we utilize posterior mean *genetic* liabilities. We note that raw LT-FH effect sizes are not on the liability scale, but can nonetheless be transformed to the observed scale. Details of the LT-FH method are provided in the Methods section; we have publicly released open-source software implementing the method (see URLs). We consider a simple example of a disease with a prevalence of 5% and narrow-sense heritability of 50%, utilizing parental history only. The posterior mean genetic liability is −0.10 for a control with 0 affected parents vs. 0.39 for a control with 1 affected parent, but GWAS treats these individuals identically. In addition, the posterior mean genetic liability is 1.56 for a case with 2 affected parents vs. 0.39 for a control with 1 affected parent, but GWAX treats these individuals identically. On the other hand, LT-FH appropriately differentiates all case-control and family history configurations.

### Simulations

We performed simulations by simulating genotypes at 100,000 unlinked SNPs and case-control status plus family history (parental history for both parents) for 100,000 unrelated target samples; we did not include sibling history in these simulations. We simulated genotypes for both parents, used these to simulate genotypes for target samples (offspring), and simulated case-control status for both parents and target samples using a liability threshold model; target samples were not ascertained for case-control status. Our default parameter settings involved 500 causal SNPs explaining *h*^2^ = 50% of variance in liability, disease prevalence *K* = 5% (implying liability threshold *T* = 1.64 and observed-scale *h*^2^ = 11%), and accurate specification of *h*^2^ and *K* to the LT-FH method; other parameter settings were also explored. We compared three methods: GWAS, GWAX, and LT-FH. Further details of the simulation framework are provided in the Methods section. We note that simulations using real LD patterns are essential for methods impacted by LD between SNPs; however, the LT-FH method (like GWAS and GWAX) is not impacted by LD between SNPs, because no genotype data is used to compute posterior mean genetic liabilities, and thus LD between the focal SNPs and other SNPs cannot impact the power of the LT-FH method at a focal SNP (aside from the fixed loss of information due to incomplete tagging of causal SNPs). We further note that simulations with LD using a subset of individuals from UK Biobank would not be feasible, as simulations of family history require genotypes of both target samples and relatives (in order to simulate the case-control status of both target samples and relatives), but genotypes of relatives are not available for (nearly all) UK Biobank samples.

We assessed the calibration of each method using average *χ*^2^ statistics at null SNPs (Figure 2a and Table S1). All methods were correctly calibrated, with an average *χ*^2^ of 1.00 for GWAS, GWAX and LT-FH. Accordingly, false-positive rates at various *α* levels (5 * 10^−2^ to 5 * 10^−6^) matched the corresponding *α* level (Table S2), as indicated by QQ plots for null SNPs (Figure S1).

**Figure 2:**
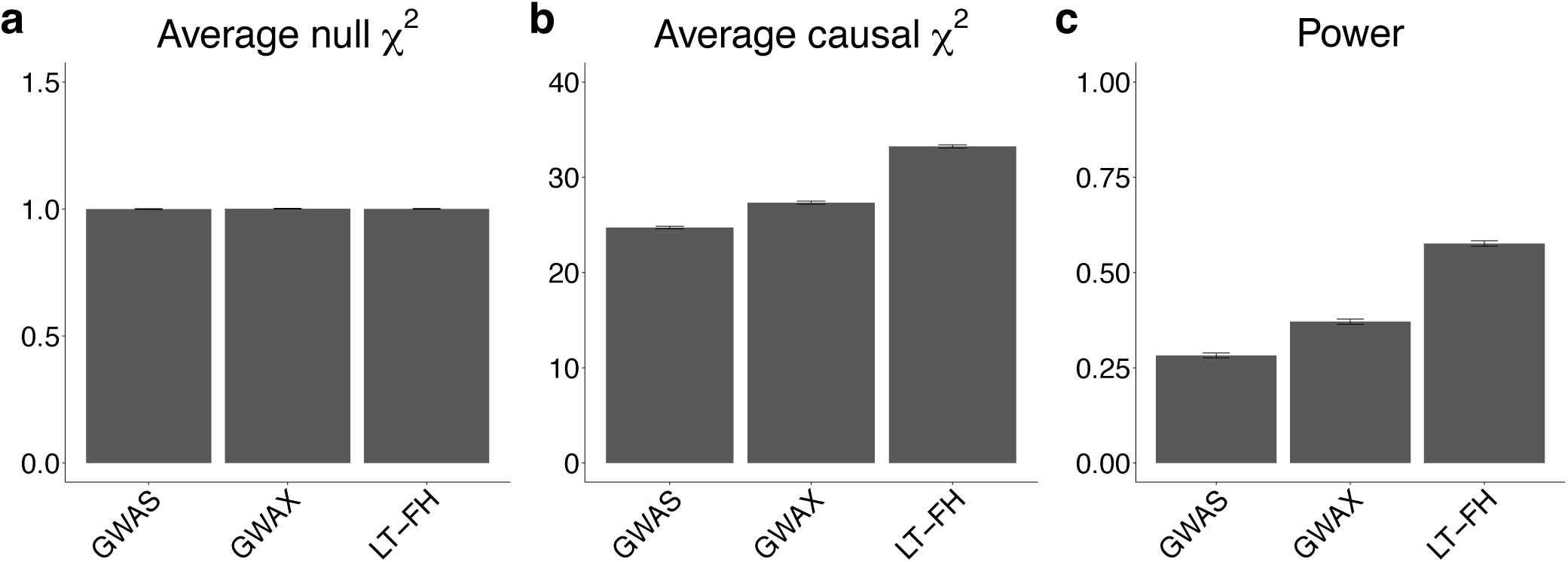
LT-FH is well-calibrated and increases association power in simulations. **(a)** Average *χ*^2^ for null causal SNPs. **(b)** Average *χ*^2^ for causal SNPs. **(c)** Power, defined as the proportion of causal SNPs with *p* < 5 * 10^−8^. Results are averaged across 10 simulations with 100,000 SNPs (500 causal SNPs) per simulation. Error bars denote standard errors. Numerical results are reported in Table S1.

We assessed the association power of each method using both average *χ*^2^ statistics at causal SNPs and formal power calculations (proportion of causal SNPs with *p* < 5 * 10^−8^) (Figure 2b,c and Table S1). LT-FH was the most powerful method, with a +22% increase in average *χ*^2^ and a +55% increase in power compared to GWAX, which outperformed GWAS at default parameter settings.

We also considered a 2 degree-of-freedom (df) extension of GWAX that treats cases, proxy cases, and controls without family history of disease as 3 distinct groups (GWAX-2df; see Methods). In contrast to previously proposed 2df extensions of GWAX^1^, our GWAX-2df test allows for incorporation of covariates, enabling its application to real traits (see below). In our simulations, GWAX-2df was well-calibrated (Table S1, Table S2 and Figure S1) and more powerful than GWAX, but LT-FH still attained a +25% increase in power compared to GWAX-2df (Table S1).

We investigated how the improvement in power attained by LT-FH varied as a function of disease prevalence *K* by performing additional simulations at lower prevalence (*K* = 1%) and higher prevalence (*K* = 25%). We varied the number of causal SNPs to approximately match the average *χ*^2^ and power (for the GWAS method) in our main simulations, and used default settings for other parameters. Results are reported in Table S3. At lower prevalence, LT-FH attained a +28% increase in power as compared to GWAX, which far outperformed GWAS. At higher prevalence, LT-FH attained a +82% increase in power as compared GWAS, which far outperformed GWAX. These results are consistent with previous findings that the GWAX method is most useful for diseases of low prevalence^1^.

We performed 12 secondary analyses to assess the robustness of LT-FH. First, we varied the number of causal SNPs by performing additional simulations with 250 or 750 causal SNPs (Table S4). As expected, decreasing (resp. increasing) the number of causal SNPs — which increases (resp. decreases) causal effect sizes — increased (resp. decreased) the power of all methods; average *χ*^2^ statistics scaled inversely with the number of causal SNPs for each method, such that the relative ordering of the methods was unchanged. Second, we varied the liability-scale heritability (*h*^2^) by performing additional simulations with *h*^2^ equal to 0.25 or 0.75 (Table S5). Decreasing (resp. increasing) the heritability led to larger (resp. smaller) improvements for GWAX and LT-FH compared to GWAS but a smaller (resp. larger) improvement for LT-FH vs. GWAX; LT-FH still attained a ≥45% increase in power compared to GWAX at each value of *h*^2^. Third, we performed simulations in which the LT-FH method utilized a misspecified value of disease prevalence (*K* = 2.5% or *K* = 7.5%, vs. true *K* = 5%) or liability-scale heritability (*h*^2^ = 0.25 or *h*^2^ = 0.75, vs. true *h*^2^ = 0.50) (Table S6). The impact on association power was negligible in each case. Fourth, we performed simulations with shared environment, which introduces a covariance in the non-genetic component of liability between parents and target samples (offspring); this covariance was set to 0.5 times the parent-offspring covariance in the genetic component of liability, i.e. 0.5 * 0.5*h*^2^ = 0.125 (Table S7). This led to smaller improvements for GWAX and LT-FH compared to GWAS but a larger (+72%) increase in power for LT-FH vs. GWAX, which still slightly outperformed GWAS. Fifth, we investigated two forms of family history reporting bias: controls failing to report family history (data not missing at random) and all controls reporting both parents as unaffected (recall bias). In each case, we observed a much smaller improvement in power of LT-FH compared to GWAS (although LT-FH was still the most powerful method tested), but we confirmed that LT-FH did not suffer from false positives (Table S8). Sixth, we evaluated an analytical approach (the Pearson-Aitken (PA) formula^2,24,25^; see Methods) for estimating posterior mean genetic liabilities. As expected, results were identical to LT-FH in simulations with no sibling history (Table S9) (but see below). Seventh, we performed simulations that include sibling history in addition to parental history; we assumed that sibling history was provided as a binary response (i.e. at least one affected sibling), as in UK Biobank data. We determined LT-FH was the most powerful method in these simulations, with a +28% increase in average *χ*^2^ and a +56% increase in power compared to GWAX, which outperformed GWAS at default parameter settings (Table S10). Eighth, we compared the (analytical) PA formula to our (Monte Carlo) LT-FH method in simulations with sibling history; in order to apply the PA formula, we assumed that exactly one sibling (rather than at least one sibling) is affected in the case of positive sibling history. We determined that our LT-FH method attained higher power than the PA formula, as a function of number of siblings and disease prevalence (Table S11). Ninth, we explored an extension of GWAX (denoted GWAX+) that assigns controls with no family history of disease a value of 0, controls with family history of disease a value of 0.5, and cases a value of 1, and uses simple linear regression to compute *χ*^2^(1 dof) statistics and p-values. We determined that GWAX+ attained lower power than LT-FH in simulations, particularly at higher disease prevalence (Table S12). Tenth, we assessed the performance of operating on the observed binary scale. We determined that operating on the binary scale performed similarly to operating on the liability scale (Table S9). Eleventh, we investigated whether modeling variance heterogeneity, defined as differences in the genetic predictor error variance across individuals, could increase power. We ran a simulation in which 10% of the individuals have case-control status data only and 90% of the individuals have case-control status plus parental history information; this represents a realistic scenario relative to the UK Biobank data set (see Supplementary Note). We considered a method that incorporates weights equal to the inverse of the genetic predictor error variance. In this scenario, the increase in power of this method ranged from −0.2% to 0.5% as a function of disease prevalence (see Supplementary Note and Table S13). This suggests that it is unlikely that modeling variance heterogeneity would substantially increase power in realistic scenarios. Finally, we assessed the concordance between LT-FH effect sizes (transformed to the observed scale) and GWAS effect sizes, and observed high concordance (slope of 1.054 (s.e. 0.005) between transformed LT-FH effect sizes and GWAS effect sizes; see Methods). In summary, LT-FH was well-calibrated and more powerful than all other methods at all parameter settings tested. However, we caution that the generative model in all of our simulations was the same as the liability threshold model that we used for inference, and thus these simulations should be viewed as a best-case scenario for LT-FH.

### Analysis of 12 diseases in the UK Biobank

We analyzed 12 diseases in the UK Biobank^27^ with genotype and case-control and/or family history data available for up to 381,493 unrelated individuals of European ancestry (20 million imputed SNPs with MAF >0.1%) and family history of disease available for most individuals (Methods, Table 1, Table S14,Table S15). Disease prevalence ranged from 0.001 for Alzheimer’s disease to 0.32 for hypertension; for lower-prevalence diseases, we applied stricter MAF thresholds to avoid type I error in unbalanced case-control settings^28^ (Table S16, Table S17). Family history information was available for each disease, including parental history (presence or absence of disease in each respective parent) and sibling history (number of brothers and sisters + presence or absence of disease in the set of all siblings). Averaged across the 12 diseases, 88% of target samples had complete parental history information and 93% of target samples had complete sibling history information (Table S18); the accuracy of family history information is assessed below. We defined GWAS, GWAX and LT-FH phenotypes accordingly (Methods, Table S19; Figure S2) and compared these three methods. We included 20 principal components (PCs), assessment center, genotype array, sex, age and age squared as covariates in association analyses for each method. In this data set, the computational cost of computing LT-FH phenotypes (posterior mean genetic liabilities) was 1-2 orders of magnitude lower than the cost of computing association statistics (Table S20).

**Table 1:** Overview of 12 diseases in the UK Biobank. We report the prevalence, parental prevalence, liability-scale SNP-heritability, and number of samples analyzed by GWAS, GWAX and LT-FH. GWAX sample sizes are lower than GWAS and LT-FH sample sizes due to incomplete family history data. * denotes sex-specific diseases (for which GWAX and LT-FH incorporate family history data from samples of both sexes). Estimates of (liability-scale) narrow-sense heritability (*h*^2^) from the literature, which are used by the LT-FH method, are reported in Table S15. Diseases are listed in order of disease prevalence. AD: Alzheimer’s disease/dementia. PD: Parkinson’s disease. COPD: Chronic bronchitis/emphysema. T2D: Type 2 diabetes. CAD: Coronary artery disease. HTN: Hypertension.

| Trait | UKBB<br>Prevalence | UKBB<br>Parental<br>Prevalence | $\hat{h}_g^2(s.e.)$ | $N_{GWAS}$ | $N_{GWAX}$ | $N_{LT-FH}$ |
| --- | --- | --- | --- | --- | --- | --- |
| AD | 0.001 | 0.065 | 0.12 (0.11) | 381493 | 324512 | 381493 |
| PD | 0.003 | 0.020 | 0.15 (0.05) | 381493 | 318792 | 381493 |
| Lung cancer | 0.006 | 0.064 | 0.10 (0.03) | 381493 | 323838 | 381493 |
| Bowel cancer | 0.013 | 0.054 | 0.14 (0.018) | 381493 | 322887 | 381493 |
| Stroke | 0.023 | 0.14 | 0.08 (0.011) | 381493 | 332468 | 381493 |
| COPD | 0.035 | 0.082 | 0.17 (0.0092) | 381493 | 329495 | 381493 |
| Prostate cancer * | 0.037 | 0.075 | 0.30 (0.019) | 175450 | 331458 | 368940 |
| T2D | 0.042 | 0.092 | 0.36 (0.0093) | 380180 | 330267 | 381390 |
| Breast cancer* | 0.061 | 0.082 | 0.20 (0.012) | 206043 | 348170 | 371064 |
| Depression | 0.073 | 0.052 | 0.11 (0.0058) | 381493 | 328186 | 381493 |
| CAD | 0.083 | 0.25 | 0.20 (0.0058) | 381493 | 341810 | 381493 |
| HTN | 0.32 | 0.26 | 0.31 (0.0037) | 381493 | 354208 | 381493 |

We assessed the calibration of GWAS, GWAX and LT-FH using stratified LD score regression (S-LDSC) attenuation ratio^21,29–31^, defined as (S-LDSC intercept -1)/(mean *χ*^2^ - 1) (Methods and Table S21a). Attenuation ratios were similar for each method, with averages across 12 diseases of 0.18, 0.16 and 0.15, respectively (and inverse variance-weighted averages close to 0.1, consistent with ref.^21^); we note that attenuation ratios slightly above 0 may be caused by attenuation bias^21^. Although there can be no guarantee that analyses of UK Biobank data using PC covariates are completely devoid of confounding^32^, these results confirm that that LT-FH is well-calibrated and there is no confounding that is specific to the LT-FH method.

We assessed the association power of each method by computing the number of independent genome-wide significant loci identified for each disease, as defined similar to ref.^21^ (Methods, Figure 3a and Table S21b); we computed standard errors (s.e.) on relative improvements via block-jackknife (Methods). Across all diseases, LT-FH was +63% (s.e. 6%) more powerful than GWAS and +36% (s.e. 4%) more powerful than the trait-specific maximum of GWAS and GWAX (e.g. 690 independent loci for LT-FH vs. 423 for GWAS). For the 8 lower-prevalence diseases (*K* < 5%), LT-FH was +39% (s.e. 7%) more powerful than the trait-specific maximum of GWAS and GWAX, and GWAX was more powerful than GWAS, consistent with simulations at lower prevalence. For the 4 higher-prevalence diseases (*K >* 5%), LT-FH was +35% (s.e. 4%) more powerful than the trait-specific maximum of GWAS and GWAX, and GWAS was more powerful than GWAX, consistent with simulations at higher prevalence; GWAS vs. GWAX results were dominated by hypertension, the disease with highest prevalence. We also evaluated the association power of each method using relative effective sample size^21^; the average relative effective sample size for was 1.31 for LT-FH vs. GWAS and 1.27 for LT-FH vs. the trait-specific maximum of GWAS and GWAX (Table S22). Finally, we assessed the association power of each method across all 12 diseases by computing the average *χ*^2^ across all genome-wide significant SNPs identified by any method (Table S21c) and average *χ*^2^ across all SNPs (Table S21d); we determined that these quantities were much larger for LT-FH than for the other methods.

**Figure 3:**
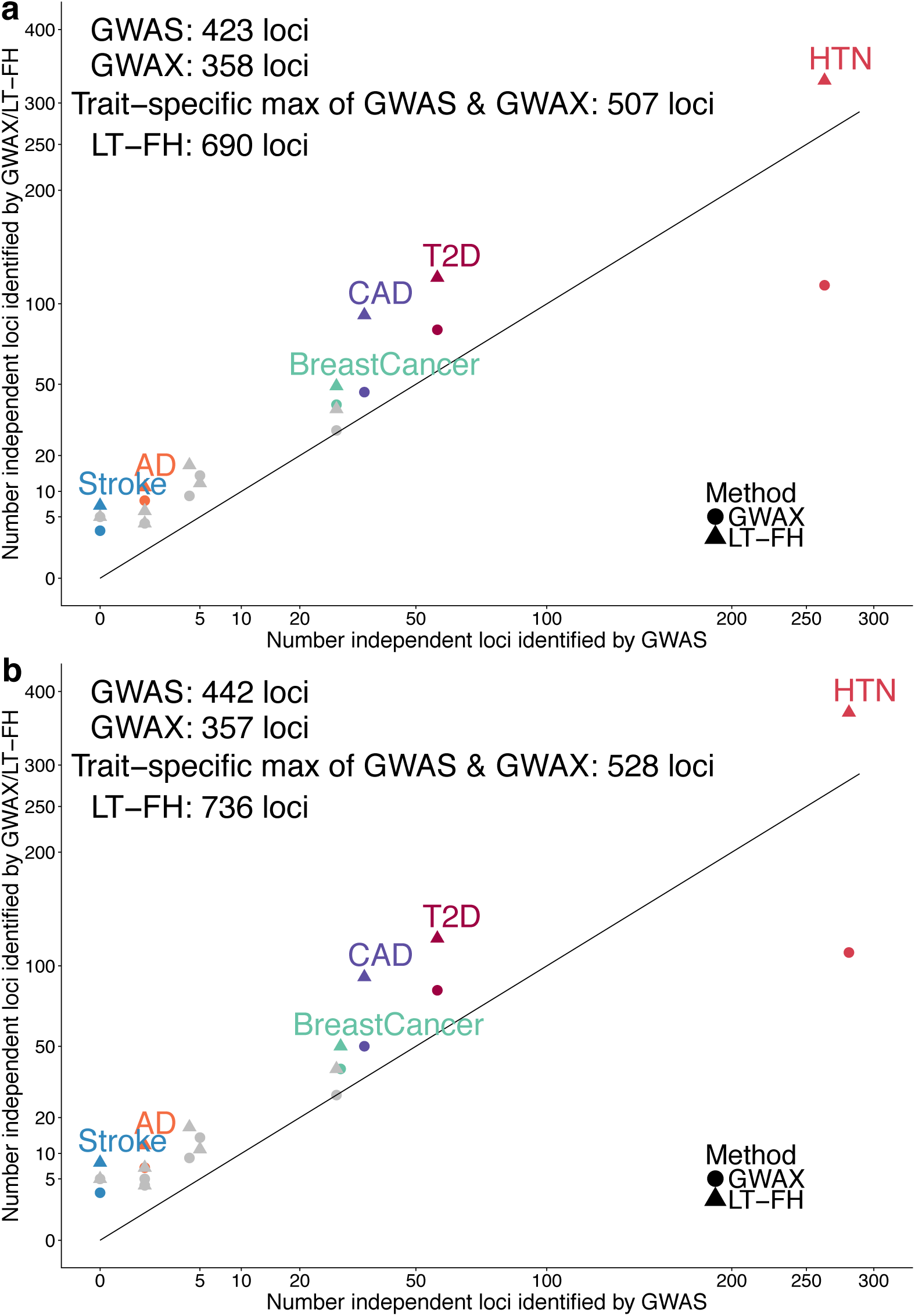
LT-FH increases association power across 12 diseases from the UK Biobank. We report results of GWAS, GWAX and LT-FH using either **(a)** linear regression or **(b)** BOLT-LMM on unrelated European individuals. Numerical results are reported in Table S21 and Table S35.

We assessed whether GWAS, GWAX and LT-FH phenotypes reflect the same underlying genetic architectures by estimating pairwise genetic correlations between these phenotypes using BOLT-REML^33^ (Table 2). Genetic correlations were high, much higher than the corresponding phenotypic correlations (e.g. 0.97 and 0.70 respectively for correlations between LT-FH and GWAS, averaged across 12 diseases), indicating that incorporating family history preserves the same underlying genetic architecture but provides substantial independent information. Phenotypic correlations for LT-FH (or GWAX) vs. GWAS were particularly low for low-prevalence diseases, for which family history is more informative than case-control status. We also computed correlations between −log_10_ association *p*-values, which mirrored the phenotypic correlations (Table S23). Notably, LT-FH phenotypes had higher values of sample size times observed-scale SNP-heritability (a measure of total genetic signal^30^) than GWAS phenotypes for all 12 diseases (59% higher on average; Table S24). These findings support the use of the LT-FH method to increase association power.

**Table 2:**
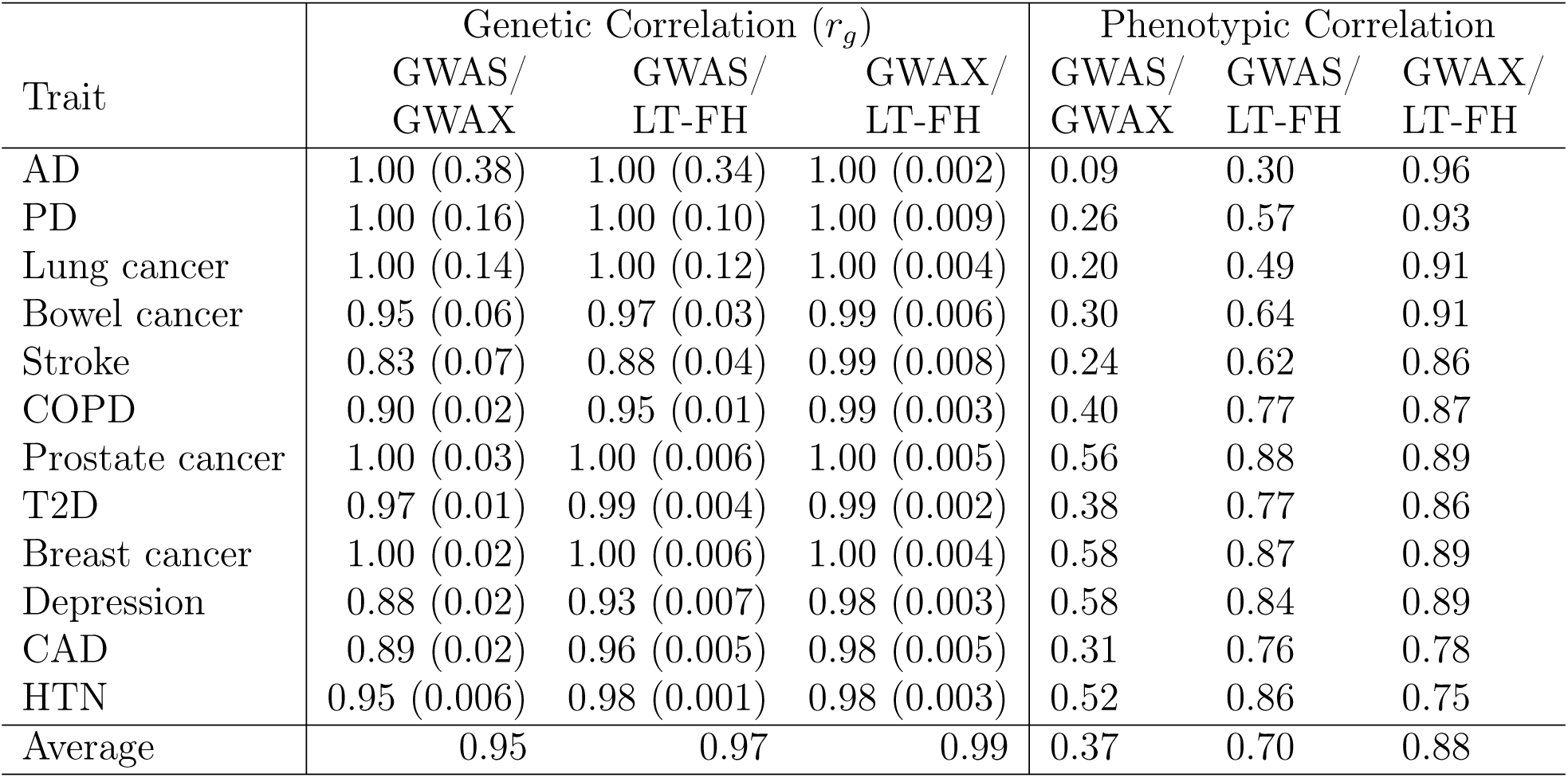
Genetic and phenotypic correlations between GWAS, GWAX and LT-FH phenotypes. We report the genetic correlation (estimated using BOLT-REML) and phenotypic correlation between each pair of GWAS, GWAX and LT-FH phenotypes. BOLT-REML estimates are constrained to be ≤1.00. Standard errors of genetic correlation estimates are reported in parentheses. Standard errors of phenotypic correlation estimates are ≤0.003 in each case.

We performed a replication analysis using independent data to assess whether the novel associations identified by LT-FH are genuine. We analyzed 4 diseases (CAD, T2D, breast cancer and prostate cancer) with publicly available summary statistics for case-control data independent from UK Biobank (see Methods). For both genome-wide significant loci identified by GWAS and genome-wide significant loci identified by LT-FH, we computed the replication slope (the slope of a regression of standardized effect sizes in case-control replication data vs. UK Biobank discovery data^34,35^); association statistics for case-control replication data were always computed using GWAS. The replication slopes were similar for GWAS (slope=0.81, se=0.02, 124 loci) and LT-FH (slope=0.79, se=0.01, 243 loci) (Figure 4 and Table S25). In addition, the replication slopes were similar for loci detected only by GWAS (slope=0.67, se=0.06, 7 loci) and loci detected only by LT-FH (slope=0.69, se=0.03, 126 loci); as expected, these slopes were lower than 0.81 and 0.79 since loci detected by only one method represent weaker effects that are more susceptible to winner’s curse. These results indicate that the novel loci identified by LT-FH are genuine associations, as assessed by the GWAS method in replication data.

**Figure 4:**
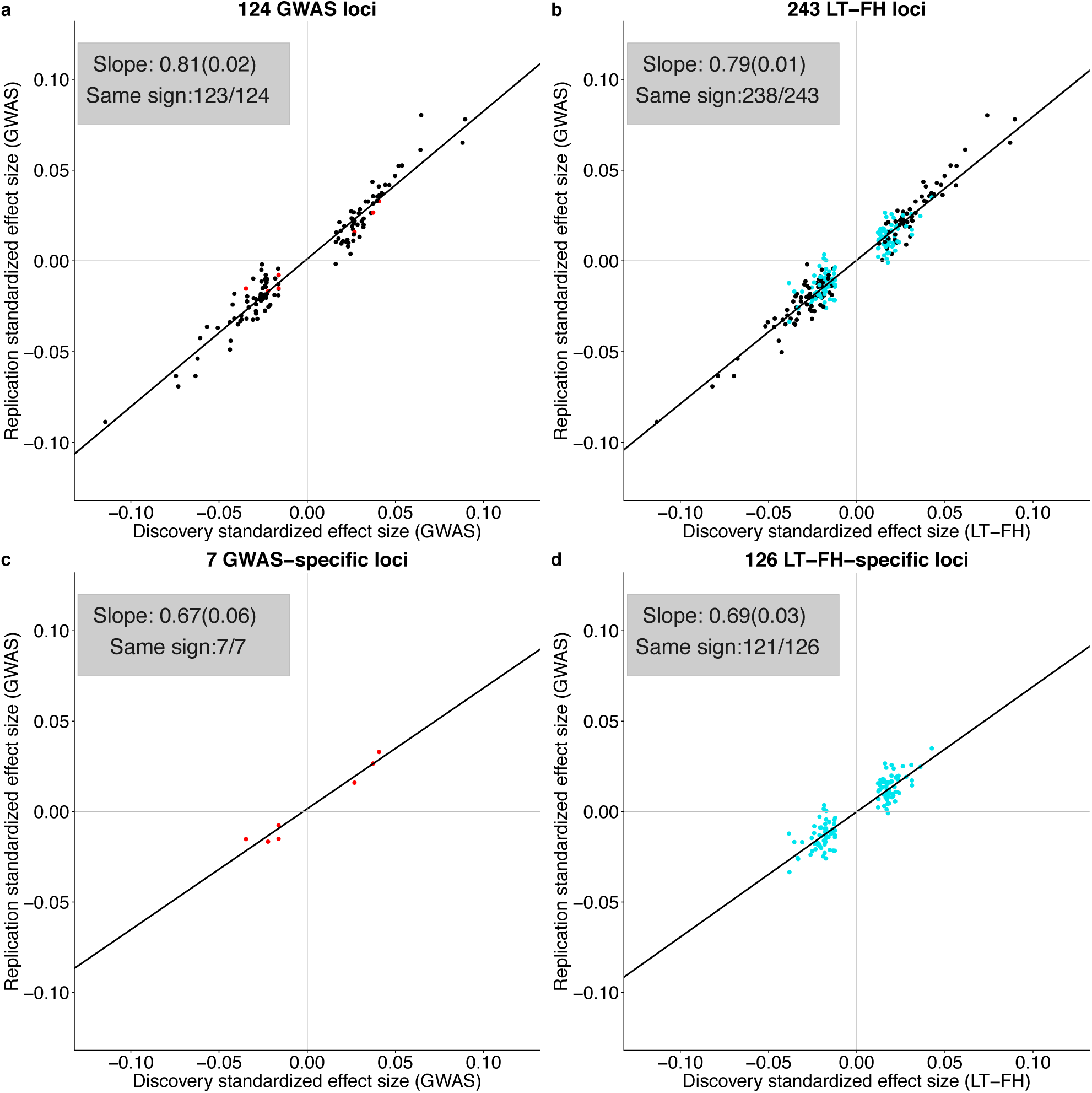
Loci identified by LT-FH replicate in independent data sets. We plot standardized effect sizes 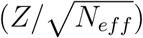 in the non-UK Biobank replication data (average *N*_*eff*_ = 99K for GWAS) vs. the UK Biobank discovery data (average *N*_*eff*_ = 62K for GWAS, 102K for LT-FH), aggregated across 4 diseases (CAD, T2D, breast cancer and prostate cancer), for (a) the 124 loci identified by GWAS, (b) the 243 loci identified by LT-FH, (c) the 7 loci identified by GWAS but not LT-FH (shown in red), and (d) the 126 loci identified by LT-FH but not GWAS (shown in turquoise). Numerical results are reported in Table S25.

We performed six secondary analyses. First, we computed association statistics for GWAX-2df, restricting this analysis to 672,292 genotyped SNPs to limit computational cost (Methods). GWAX-2df was substantially more powerful than GWAX (assessed using the number of independent loci), but this had little impact on our conclusions about LT-FH, which was +23% (s.e. 3%) more powerful than the trait-specific maximum of GWAS and GWAX-2df (vs. +35% (s.e. 4%) more powerful than the trait-specific maximum of GWAS and GWAX) in analyses of genotyped SNPs (Table S26). Second, we computed association statistics for analogues of GWAX and LT-FH that incorporate only parental history and not sibling history (GWAX_*no*−*sib*_ and LT-FH_*no*−*sib*_). We determined that using only parental history was slightly less powerful, e.g. LT-FH_*no*−*sib*_ was +54% (s.e. 5%) more powerful than GWAS (whereas LT-FH was +63% (s.e. 6%) more powerful than GWAS) (Table S27). Third, we assessed the accuracy of family history information by computing the correlation of self-reported family history between siblings (Methods and Table S28). Averaged across traits, the correlation between siblings was 0.685 for number of affected parents and 0.583 for presence or absence of disease in the set of all siblings; it follows that the correlation between true and self-reported family history is equal to the square root of these numbers (0.827 and 0.764), if errors are uncorrelated between siblings. To investigate the importance of accounting for inaccurate family history information, we modified the LT-FH method to downweight family history information based on its accuracy for each disease (Methods), but we determined that this had little impact on association results (0% increase in power for modified LT-FH vs. LT-FH, average phenotypic correlation = 0.996 across 12 diseases; Table S29). Fourth, we investigated whether LT-FH could be improved by explicitly accounting for age when computing posterior mean genetic liabilities^16^ (Table S30, Table S31). We determined that this had little impact on association results (< 2% increase in power; Table S32, Figure S3), consistent with previous findings that including age as a simple covariate is sufficient in case-control studies with random ascertainment^16^. Fifth, for the 8 lower-prevalence diseases for which we applied stricter MAF thresholds to avoid type I error in unbalanced case-control settings^28^, we recomputed association statistics for GWAX and LT-FH using MAF thresholds chosen specifically for each method, based on the kurtosis of the corresponding phenotypes (Table S16, Table S17; the number of diseases requiring stricter MAF thresholds reduces to 0 for GWAX and 2 for LT-FH, due to lower kurtosis). The number of independent loci identified by GWAX and LT-FH increased for these diseases, but overall results were little changed, e.g. LT-FH was +37% (s.e. 4%) more powerful than the trait-specific maximum of GWAS and GWAX (vs. +36% (s.e. 4%) more powerful than the trait-specific maximum of GWAS and GWAX in our primary analysis) (Table S33). Finally, we compared the (analytical) PA formula^2,24,25^ to our (Monte Carlo) LT-FH method in analyses of UK Biobank diseases. As in simulations, in order to apply the PA formula, we assumed that exactly one sibling (rather than at least one sibling) is affected in the case of positive sibling history. We determined that our LT-FH method attained higher power than the PA formula, identifying 12 more genome-wide significant loci in aggregate, a 2% relative improvement for LT-FH versus the PA formula (jackknife P= 0.01 for difference; Table S34).

In summary, LT-FH was well-calibrated and more powerful than all other methods in all primary and secondary analyses.

### BOLT-LMM increases GWAS, GWAX and LT-FH association power

BOLT-LMM, a method for efficient Bayesian mixed-model analysis, has been shown to increase association power (for GWAS phenotypes) as compared to linear regression^20,21^. We thus investigated the application of BOLT-LMM to GWAS, GWAX and LT-FH phenotypes, analyzing the same data for 12 UK Biobank diseases as above. We first confirmed that GWAS, GWAX and LT-FH remained well-calibrated (Table S35a). We then assessed association power (Figure 3b and Table S35b). Across all diseases, LT-FH was +67% (s.e. 6%) more powerful than GWAS and +39% (s.e. 4%) more powerful than the trait-specific maximum of GWAS and GWAX (e.g. 736 independent loci for LT-FH vs. 442 for GWAS), implying similar relative power among the three methods when using BOLT-LMM; the absolute increases in power attained by using BOLT-LMM were modest (Figure 3b vs. Figure 3a) and smaller than in analyses of highly heritable UK Biobank traits^21^, because disease traits have lower observed-scale SNP-heritability (particularly for lower-prevalence diseases) and the increase in power attained by BOLT-LMM scales with SNP-heritability^21^. Analyses of average *χ*^2^ across all genome-wide significant SNPs identified by any method (Table S35c) and average *χ*^2^ across all SNPs (Table S35d) yielded similar conclusions.

Our above analyses using both linear regression and BOLT-LMM were restricted to unrelated target samples (up to 381,493 unrelated individuals of European ancestry), consistent with previous studies leveraging family history using GWAX^1,10–12^. We investigated the consequences of including related individuals in BOLT-LMM analyses. We first applied BOLT-LMM to GWAS, GWAX and LT-FH phenotypes of target samples defined without applying any filter for relatedness (up to 459,256 related individuals of European ancestry). We determined that GWAX and LT-FH suffered poor calibration (average S-LDSC attenuation ratios equal to 0.26 for GWAX and 0.22 for LT-FH, vs. 0.14 for GWAS) (Table S36a), rendering moot the increases in power due to larger sample size (Table S36b). A likely explanation for the poor calibration is the extreme concordance between sibling pair phenotypes (close to 100% for GWAX) arising from nearly identical family histories. Our results indicate that BOLT-LMM is unable to correct for inflation in test statistics in such extreme cases, as it does not have the flexibility to model excess phenotypic concordance for sibling pairs specifically. Thus, BOLT-LMM analysis of GWAX and LT-FH phenotypes should not be applied to target samples without any filter for relatedness.

We thus applied BOLT-LMM to LT-FH phenotypes for all target samples (up to 459,256 related individuals of European ancestry) that were modified to incorporate only case-control status (ignoring family history information) for each individual from a sibling pair or parent-offspring pair within the set of target samples (also see secondary analysis below). We determined that this approach was well-calibrated, with an average S-LDSC attenuation ratio of 0.14 for LT-FH, vs. 0.14 for GWAS (and inverse variance-weighted average close to 0.1, consistent with ref.^21^) (Table S37a). We compared the power of this approach to applying BOLT-LMM to GWAS for all related Europeans (see above) and GWAX for all unrelated Europeans (as previously recommended^1,10–12^; see above). Across all diseases, LT-FH was +54% (s.e. 5%) more powerful than GWAS and +44% (s.e. 4%) more powerful than the trait-specific maximum of GWAS and GWAX (e.g. 908 independent loci for LT-FH vs. 590 for GWAS) (Figure S4 and Table S37b), similar to our previous analyses (Figure 3). Thus, BOLT-LMM analysis of LT-FH phenotypes defined using only case-control status for all sibling pairs and parent-offspring pairs within the set of target samples is the recommended approach; we have publicly released LT-FH association statistics computed using this approach (see URLs).

We performed a secondary analysis in which we instead applied BOLT-LMM to LT-FH phenotypes for all target samples (up to 459,256 related individuals of European ancestry) that were modified to incorporate family history information for exactly one sibling for each set of siblings within the set of target samples (with no filter on family history information for parent-offspring pairs). We note that this approach may suffer from residual concordance between sibling pair phenotypes (and parent-offspring phenotypes) arising from redundancy between the family history of one target sample and the case-control status of the other target sample. Indeed, we determined that LT-FH phenotypes defined in this way were slightly but significantly miscalibrated (average S-LDSC attenuation ratio equal to 0.15 for LT-FH, vs. 0.14 for the recommended LT-FH approach; *p* < 10^−6^ for difference in inverse-variance weighted means) (Table S38a), rendering moot the slightly higher power of this approach (Table S38b).

In summary, BOLT-LMM analysis increases the power of LT-FH (and GWAS and GWAX), but care is required to avoid poor calibration of LT-FH (and GWAX) due to extreme concordance in sibling pair phenotypes. Our recommendation is for BOLT-LMM analysis of LT-FH phenotypes defined using only case-control status for all sibling pairs and parent-offspring pairs within the set of target samples.

## Discussion

We have introduced a new method, LT-FH, that accurately models a broad range of case-control status and family history configurations to increase association power compared to prevailing methods^1^; in particular, LT-FH differs from other methods, including methods developed in the livestock genetics community^5–9^, by allowing for complex modes of reporting family history (e.g. binary response for sibling history, i.e. at least one sibling has the disease), instead of requiring phenotypes for each relative. Across 12 diseases from the UK Biobank, LT-FH was +63% (s.e. 6%) more powerful than GWAS and +36% (s.e. 4%) more powerful than the trait-specific maximum of GWAS and GWAX, while maintaining correct calibration. LT-FH attained similar increases in power when using BOLT-LMM to compute association statistics for all three methods, and when incorporating related individuals. These findings provide a strong motivation to apply LT-FH to genetic data sets for which family history information is available, and to collect family history information in future genetic studies.

Although LT-FH greatly increases association power, it has several limitations. First, self-reported family history information may be inaccurate. Indeed, we determined that the sibling concordance of self-reported family history was substantially lower than 100% (Table S28); however, our efforts to account for this had little impact on association power (Table S29). Second, family history may reflect a different underlying genetic architecture than case-control status, e.g. due to differences in the etiology of early-onset vs. late-onset disease or differences in diagnostic criteria over time; however, we observed high genetic correlations between GWAS and LT-FH phenotypes (average of 0.97 across diseases; Table 2). Third, real diseases may not adhere to the liability threshold model used by LT-FH, which may explain why increases in power in analyses of real diseases (Figure 3) were not as large as in simulations (Figure 2c). Moreover, posterior mean genetic liabilities inferred from family history information may be biased by unmodeled effects. This might reduce the power of the LT-FH method, but would not cause its test for additive association with genetic variants to produce false positives (Table S7 and Table S8) Fourth, just like standard GWAS analysis, LT-FH may require MAF thresholds stricter than our default value of 0.001 to avoid type I error in unbalanced case-control settings^28^. Although the number of diseases requiring stricter MAF thresholds was reduced from 8 for standard GWAS analysis to 2 for LT-FH (due to lower kurtosis; Table S16), more sophisticated approaches (generalizing ref.^28^) could be explored. Fifth, just like GWAX, LT-FH should not be applied to target samples without any filter for relatedness, because extreme concordance in sibling pair phenotypes leads to miscalibration — even when applying BOLT-LMM, an efficient mixed-model method that is intended to account for phenotypic concordance among related individuals. Specifically, GWAS analysis of related target samples can induce miscalibration because target samples have correlated genotypes and correlated phenotypes. Mixed model methods, such as BOLT-LMM, are designed to correct this miscalibration. However, if related target samples have *extremely* concordant phenotypes due to inclusion of family history information, BOLT-LMM may fail to correct this miscalibration (Table S36). Thus, scenarios of *extreme* concordance between phenotypes of related target samples should be avoided. We instead recommend applying BOLT-LMM to LT-FH phenotypes defined using only case-control status for all sibling pairs and parent-offspring pairs within the set of target samples; this approach is straightforward to implement and still greatly increases association power while maintaining correct calibration. Sixth, LT-FH does not currently incorporate family history information for genetically correlated diseases, which could be highly informative; however, LT-FH association results for two or more genetically correlated diseases could be jointly analyzed using existing methods^36^. Seventh, the novel genome-wide significant associations detected by LT-FH explain less variance on average, such that the increase in variance explained by LT-FH compared to other methods is lower than the increase in number of genome-wide significant loci. However, small genetic effects can still produce important biological insights^37,38^. Eighth, posterior mean genetic liabilities can be computed using an analytical approach, the PA formula^2,24,25^, instead of LT-FH. However, the PA formula is not well-suited to data that includes sibling history as a binary response (i.e. at least one sibling has the disease). Indeed, we confirmed that LT-FH attains higher power than the PA formula in simulations with sibling history (Table S11) and analyses of UK Biobank diseases (Table S34). Finally, we have not explored the use of family history to improve the accuracy of genetic risk prediction^2,3^, which remains as a direction for future research. Despite all these limitations, we anticipate that LT-FH will attain large increases in power in future association studies.

## URLs

UK Biobank: http://www.ukbiobank.ac.uk/. BOLT-LMM v2.3 software: https://data.broadinstitute.org/alkesgroup/BOLT-LMM. LT-FH software (v1 and v2): https://data.broadinstitute.org/alkesgroup/UKBB/LTFH/. LT-FH summary association statistics for 12 diseases: https://data.broadinstitute.org/alkesgroup/UKBB/LTFH/sumstats/. LTSOFT software: https://data.broadinstitute.org/alkesgroup/LTSOFT/. PLINK software: https://www.cog-genomics.org/plink2.

## Methods

### LT-FH method

The liability threshold model can be written as *ϵ* = ***Xβ*** + *ϵ*_*e*_, where *ϵ* is the liability, ***X*** is the genotypes at candidate SNPs (normalized to mean 0, variance 1), *β* are the effect sizes of SNPs on the liability scale, and an individual is a case (*z* = 1) if and only if *ϵ* ≥ *T* and is a control otherwise (*z* = 0). *T* determines the disease prevalence (Φ(*T*) = *P* (*x* ≥ *T*) where *x* ∼ *N* (0, 1)). We further assume that *ϵ*_*g*_ = ***Xβ*** ∼ *N* (0, *h*^2^), and assume that when testing a SNP *g* for association, the effect size is small enough such that the liability is *βg* + *ϵ* = *βg* + *ϵ*_*g*_ + *ϵ*_*e*_ where *ϵ* ∼ *N* (0, 1), *ϵ*_*g*_ ∼ *N* (0, *h*^2^), *ϵ*_*e*_ ∼ *N* (0, 1 − *h*^2^).

We assume a multivariate normal distribution for ***ϵ***: for two individuals *Cov*(*ϵ*_1_, *ϵ*_2_) = *K*_12_*h*^2^, where *K*_12_ is the coefficient of relationship for the pair of individuals (e.g. 1*/*2 for parent-offspring). We assume that we have the genotype and case-status of an individual, and potentially the disease status of family members. For example, to incorporate parental history and sibling history we assume

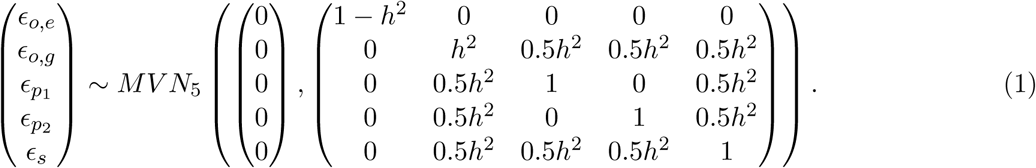

where *ϵ*_*o,e*_, *ϵ*_*o,g*_ are the environmental and genetic components of the liability of the target sample (offspring), *ϵ*_*p*1_ and *ϵ*_*p*2_ are the liabilities of the parents, and *ϵ*_*s*_ is the liability of the sibling(s) (for simplicity we include only one sibling here, but this can be extended to an arbitrary number of siblings). We estimate the posterior mean genetic liability 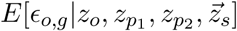 for each individual given the case-control status of the genotyped individual (*z*_*o*_), both parents (*z*_*p*1_, *z*_*p*2_), and sibling(s) 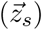. When disease status is unknown for subsets of these individuals, we condition only on known information. We are interested in genetic signal and therefore we estimate the posterior mean genetic liability, rather than the posterior mean liability; genetic association power is increased by integrating out the noise contributed by the environmental component of liability.

We estimate *E*[*ϵ*_*o,g*_ | *z*_*o*_, *z*_*p*1_, *z*_*p*2_, ***z***_*s*_] using Monte Carlo integration. Briefly, (1) we sample from a multivariate normal with the given covariance structure and compute the posterior mean genetic liability of the offspring in all possible configurations of case-control status and family history (using liabilities that fall above or below a given threshold to determine case-control status and family history). If the posterior mean genetic liabilities in any configuration has a standard error of the mean above 0.01, we resample from a conditional multivariate normal to ensure that we sample in a manner that represents the tails of the distribution appropriately. (2) We sample relevant liabilities conditional on the offspring’s case-control status (*z*_*o*_) or we sample conditional on the parents (i.e.*z*_*p*1_ + *z*_*p*2_ = {0, 1, 2}). (3) We then combine all samples of *E*_*o,g*_ from given configurations. (4) We compute posterior mean genetic liabilities and assess the standard error of each posterior mean. We then repeat (2)-(4) until the standard error of the mean is less than 0.01 for all configurations. We chose 0.01 as convergence criterion as the differences between posterior mean genetic liabilities of individuals with different case-control and family history configurations are generally much larger than 0.01. In simulations, we determined the results were virtually identical regardless of the choice of convergence criterion (Table S39). We note that each round of our Monte Carlo integration procedure produces an independent sample of the target individual’s genetic liability from the posterior distribution (conditional on the target individual’s case-control status and family history, and independent of previous rounds), i.e. we are not performing Gibbs sampling or Markov Chain Monte Carlo. We also note that family history reporting bias may reduce the power of LT-FH, but would not lead to false positives (Table S8). In addition, distinct from family history, non-random missingness of target sample phenotypes would equally impact GWAS and LT-FH. Specifically, LT-FH is identical to GWAS in the absence of family history information, because in this setting both methods exclude individuals with missing case-control status.

In UK Biobank data, sibling history is provided as a binary response, i.e., at least one sibling has the disease; this “at least one” condition is straightforward to incorporate using Monte Carlo integration. For a given number of siblings (*S*), we sample *S* liabilities (*ϵ*_*s*1_, *ϵ*_*s*2_,…, *ϵ*_*sS*_) and can easily assess the presence or absence of disease in the set of all siblings by comparing each sibling’s liability to the relevant threshold; the absence of disease corresponds to *ϵ*_*si*_ < *T* for all *i* ∈ {1, 2,…, S} whereas the presence of disease corresponds to at least one *i* with *ϵ*_*si*_ ≥ *T*. Thus, we estimate posterior mean genetic liabilities using Monte Carlo integration in our main analyses and open-source software.

The LT-FH association statistic is a measure of association between genotype *g* and posterior mean genetic liability across samples. We can compute this statistic either using linear regression or using other methods such as BOLT-LMM^20,21^, while incorporating additional covariates such as principal components. If we treat the posterior mean genetic liability as a continuous variable and compute the number of samples times the squared correlation between *g* and posterior mean genetic liability (generalizing the Armitage trend test^26^), this is equivalent to the Score test (see Supplementary Note).

### LT-FH effect sizes

Although LT-FH estimates the posterior mean genetic liability, the raw effect sizes computed when using LT-FH are not on the liability scale. However, the raw effect sizes computed when using linear regression can be transformed to the observed scale by making use of the effective sample size^21^. In detail, to obtain per-allele observed-scale effect sizes for a non-standardized phenotype (as is computed by BOLT-LMM software) we compute:

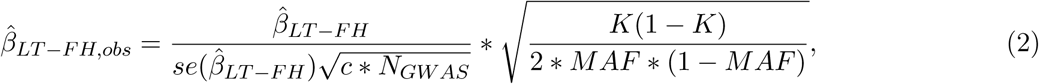

where 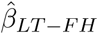 is raw per-allele effect size, *c* is the relative effective sample size for LT-FH vs GWAS, *K* is the disease prevalence, and *MAF* is the minor allele frequency. We note that it is straightforward to use minor allele frequency (MAF) to convert between per-allele and per-standardized-genotype effect sizes, distinct from converting between liability and observed scales. For our default simulation scenario we computed linear regression effect sizes using `lm()` in `R`. LT-FH effect sizes were transformed as described in (2) with *c* computed for each simulation replicate as the median ratio of LT-FH *χ*^2^ statistics to GWAS *χ*^2^ statistics across SNPs with *χ*^2^ ≥30 in GWAX^21^. We observed a strong concordance (slope of 1.055, standard error of 0.005) between 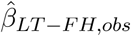 and 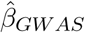 for genome-wide significant effect sizes (defined as P ≤ 5 * 10^−8^ for both GWAS and LT-FH). This slope may differ slightly from 1 due to the noise in estimating *c*.

BOLT-LMM software only produces effect size estimates from the BOLT-LMM approximation to infinitesimal mixed model (*β*_*BOLT*−*LMM*−*inf*_). When applying BOLT-LMM to real traits, we observed a strong concordance between GWAS and raw LT-FH BOLT-LMM-inf effect sizes for genome-wide significant effect sizes (Table S40, average weighted correlation of 0.996). LT-FH BOLT-LMM-inf effect sizes can be transformed to the observed scale as described in (2) with *c* equal to the relative effective sample size for LT-FH BOLT-LMM-inf vs. GWAS (computed as the median ratio of LT-FH BOLT-LMM-inf *χ*^2^ statistics to GWAS linear regression *χ*^2^ statistics across genotyped SNPs with *χ*^2^ ≥30 in GWAS applied via BOLT-LMM to all related Europeans^21^; see Table S41). We used the relative effective sample size for LT-FH BOLT-LMM-inf vs. GWAS using linear regression to reflect the power gained by using both LT-FH (vs. GWAS) and BOLT-LMM-inf (vs. linear regression). We observed a strong concordance between GWAS BOLT-LMM-inf effect sizes and transformed LT-FH BOLT-LMM-inf effect sizes (slope of 0.94; Figure S5).

### PA formula

The Pearson-Aitken (PA) formula is an analytical approach that can be used to estimate mean vectors and covariance matrices after selecting on subsets of variables^2,24,25^. Consider sets of variables that can be partitioned into ***x*** and ***y*** with mean vector and covariance matrix:

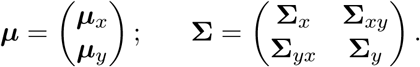

If we select on subset ***x***, and the mean of ***x*** becomes 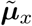 and variance becomes 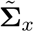, then the mean vector and covariance matrix becomes:

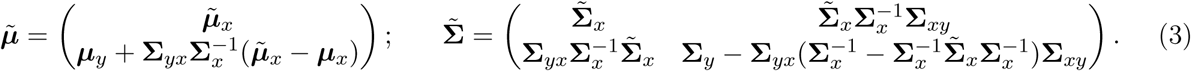

The PA formula can be used to obtain posterior mean genetic liability values for individuals conditional on their own case-control status as well as the disease status of their first degree relatives (see (1)).

We considered the Pearson-Aitken (PA) formula as an approximate analytical approach and implemented this approximation for LT-FH_*no*−*sib*_^2,24,25^ (Table S32). However, in UK Biobank data, sibling history is provided as a binary response, i.e., at least one sibling has the disease. This “at least one” condition is complex to incorporate using analytical approaches but straightforward to incorporate using Monte Carlo integration. We applied the PA formula in the case of sibling history by assuming that exactly one sibling (rather than at least one sibling) is affected in the case of positive sibling history (Table S11 and Table S34). We have included an implementation of the PA formula in our LT-FH software for use in data sets that do not include sibling history as provided by UK Biobank.

### GWAS and GWAX methods

GWAS aims to discover associations between variants and disease by comparing cases to controls of a disease. GWAX compares cases and proxy cases (controls with a family history of the disease) to controls with no family history of disease. GWAX can be particularly valuable when the case-control status of the genotyped individual is unknown, however when the case-control status of the genotyped individual is known incorporating this information increases power^1^, and as such we incorporate the case-control status of the genotyped individual (if known) when computing GWAX association statistics. For both GWAS and GWAX, association statistics can be computed using methods such as linear regression, logistic regression, BOLT-LMM^21^, or SAIGE^28^.

For GWAX-2df test we perform an F-test, regressing genotype on case status and control with family history status (two binary variables) and testing whether the coefficients associated with these two indicator variables are both 0 (see Supplementary Note). When no covariates are included, results are virtually identical to a Pearson-*χ*^2^ test on a 2×3 table.

#### Simulations

We simulate genotypes at 100,000 unlinked SNPs and case-control status plus family history (parental history for both parents) for 100,000 unrelated target samples; we do not include sibling history in these simulations. We simulate genotypes for both parents and use these to simulate genotypes for target samples (offspring); the underlying MAF is ∼ *U*(0.01, 0.5). Using these genotypes we simulate case-control status for both parents and target samples using a liability threshold model in which we normalize genotype to a mean of 0 and variance of 1 according to the true MAF, set the effect sizes for the *C* causal SNPs on the liability scale to be 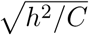, and add environmental noise with a variance of 1 − *h*^2^. Target samples are not ascertained for case-control status. Our default parameter settings involve 500 causal SNPs explaining *h*^2^ = 50% of variance in liability, disease prevalence *K* = 5% (implying liability threshold *T* = 1.64 and observed-scale *h*^2^ = 11%), and accurate specification of *h*^2^ and *K* to the LT-FH method. For each parameter setting we compute 10 simulation replicates in which there is a different underlying MAF distribution for each simulation replicate.

We apply linear regression (implemented in BOLT-LMM software) to obtain *χ*^2^ statistics for GWAS (case vs. control), GWAX (case+proxy-case vs. control), and LT-FH (*E*[*ϵ*_*o,g*_ | *z*_*o*_, *z*_*p*1_, *z*_*p*2_] for each individual). We compute the posterior mean genetic liability for each of the 6 configurations of case-control status and family history using 1,000,000 values of *ϵ*_*g*_ sampled conditional on the 3 parent history configurations (i.e.*z*_*p*1_ + *z*_*p*2_ = {0, 1, 2}).

We investigate 12 simulation scenarios total in which we vary *C* (number of causal SNPs), *K* (prevalence), *h*^2^, the assumed *h*^2^ and prevalence when calculating *E*[*ϵ*_*o,g*_|*z*_*o*_, *z*_*p*1_, *z*_*p*2_] for each individual, and the environmental correlation between parents and offspring (Table S1-Table S7). For simulations in which we vary parameters other than *C* we manipulate *C* such that the average *χ*^2^ for causal SNPs for GWAS is approximately the same as in the default parameter setting for comparative reasons. Using the same underlying genotypes and phenotypes as in the default parameter setting, we investigate the effect of mis-specifying 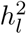 and *K* on the performance of LT-FH. When introducing environmental covariance we assume the following covariance structure between the non-genetic (environmental) components of the offspring (*o*) and the two parents (*p*_1_, *p*_2_):

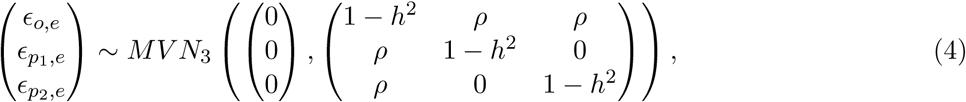

where in simulations we let *ρ* = 0.5 * (0.5*h*^2^) (half of the genetic covariance between offspring and parents). Note that in this scenario we are assuming the parents still share no environmental covariance.

### UK Biobank data set

We analyzed genetic data from the UK Biobank^27^ consisting of 381,493 unrelated individuals or 459,256 related individuals of European ancestry (based on self-reported white ethnicity; sample sizes depended on the trait analyzed; Table 1; Table S14) and ∼20 million imputed variants with MAF >0.1%^21^. We identified pairs of siblings (full siblings and MZ twins) as outlined previously^27,39^. We note that it is easy to distinguish sibling pairs from parent-child pairs using identity-by-descent (IBD). We identified parent-offspring pairs as individuals having a kinship in (0.177, 0.354) and having proportion of SNPs with identity-by-state equal to 0 (IBS0) < 0.0012. We computed association statistics applying either linear regression or default BOLT-LMM analysis (both implemented in BOLT-LMM software). We note that for this default analysis, BOLT-LMM association statistics reduce to infinitesimal mixed model association statistics (BOLT-LMM-inf) when the non-infinitesimal BOLT-LMM association statistics are not expected to increase power^20,21^. We controlled for assessment center, genotype array, sex, age, age squared, and the first 20 principal components^21^.

We considered 12 diseases in UK Biobank for which family history of disease was available (Table 1); these 12 diseases were considered when GWAX was proposed^1^. These 12 diseases had a wide range of narrow-sense heritabilities (Table S15). We assigned case-control status for all genotyped individuals (Table S14). We assigned individuals with no ICD9/10 codes as controls. An alternative would be to assign missing phenotypes to individuals with no ICD9/10 codes, which reduces sample size while increasing disease prevalence (Table S42). We decided against this possibility, because we determined that it slightly but consistently reduces sample size times observed-scale SNP-heritability (a measure of total genetic signal^30^) and has very little impact on GWAS power (Table S43)

Parental history and sibling history information was available for all 12 diseases. Parental history information consisted of presence or absence of disease in each respective parent; we note that the parental prevalence was often greater than the disease prevalence (Table 1). Sibling history information consisted of number of brothers and sisters + presence or absence of disease in the set of all siblings. Family history information is self-reported and thus there is a risk of misreporting, recall bias, or phenotype misclassification. For example, participants were asked about their family history of “diabetes” but we assume that this family history corresponds to that specifically of type II diabetes, consistent with previous studies^1^. To assess the accuracy of family history information we compute the correlation of self-reported family history between siblings. When computing the correlation of self-reported sibling history we restricted to concordant sibling pairs (e.g. both cases or both controls) and for sex-specific diseases (breast and prostate cancer) we restricted to concordant sibling pairs of the relevant sex, sibling pairs of the non-relevant sex, and sibling pairs of discordant sex where the relevant sex is a control. We observed a moderately high concordance of self-reported family history among siblings (Table S28).

### LT-FH in UK Biobank

There are 377 possible configurations of case-control status and family history of disease. For example, there are 2 configurations when case-control status is available but no family history is available (case or control) and 4 configurations when case-control status and one parent’s disease status is available (case with affected parent, case with unaffected parent, control with affected parent, control with unaffected parent). In detail, there are 252 configurations when case-control status is available: 2 configurations of case-control status with no family history (see above); 4 of case-control status and one parent’s disease status (see above); 6 of case-control status and both parents’ disease status; 40 of case-control status and sibling’s disease status (1-10 siblings; at least one or none affected); 80 of case-control status, one parent’s disease status, and sibling’s disease status; 120 of case-control status, both parents’ disease status, and sibling’s disease status. There are 125 configurations when case-control status is unavailable: 2 of one parent’s disease status; 20 of sibling’s disease status; 3 of both parents’ disease status; 40 of one parent’s disease status and sibling’s disease status; 60 of both parents’ disease status and sibling’s disease status. A comprehensive list of configurations is provided in Table S45).

We condition on known information to compute the posterior mean genetic liability. We computed *E*[*ϵ*_*o,g*_|·] (where · is all known disease status information) through Monte Carlo integration as described above. We allow for genotyped individuals and their siblings to have a different disease prevalence than their parents. We select the prevalence for genotyped individuals and their siblings using the prevalence of the disease in genotyped (unrelated European) individuals, and the prevalence for parents using the parental disease prevalence (for parents of unrelated European individuals). *h*^2^ values were obtained from the published literature (Table S15). We used *h*^2^ estimates from twin studies, but we note the existence of *h*^2^ estimates based on family history^40^.

For individuals who report having 0 siblings, more than 10 siblings, or who do not know or prefer not to answer how many brothers and/or sisters they have, we ignore sibling history and use parental history only for LT-FH; for all other individuals we incorporate both the number of siblings as well as whether or not at least one is affected in the computation of *E*[*ϵ*_*g*_|·]. For sex-specific diseases our exclusion criteria remains consistent however we now incorporate the number of siblings of the relevant sex as well as whether or not at least one is affected in the computation of *E*[*ϵ*_*g*_|·].

In a secondary analysis, we modified the LT-FH method to downweight family history information based on its accuracy for each disease such that 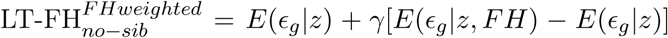 where *z* was case-control status, FH represented parental history information (*z*_*p*1_, *z*_*p*2_), and *γ* was the accuracy of parental history estimated through sibling disease concordance rates (Table S28). Thus, if reported history was 0% accurate we assign the LT-FH phenotype to be *E*(*ϵ*_*g*_ |*z*_*o*_) (*z*_*o*_ is case-control status of genotyped individual), if reported history was 100% accurate we assign the LT-FH phenotype as before, *E*(*ϵ*_*g*_|*z*_*o*_, *z*_*p*1_, *z*_*p*2_), we use linear interpolation for intermediate values of accuracy.

### GWAS and GWAX in UK Biobank

We implemented GWAS considering the case-control status of the genotyped individual only (Table S19). The prevalence ranged from 0.1% to 32% (Table 1). Due to the increased risk of a type I error in unbalanced case-control settings, for some diseases we selected a stringent MAF threshold when computing the number of independent loci (see below)^21,28^. However, one could also use a method such as SAIGE that controls type I error in unbalanced case-control settings^28^.

We considered the case-control status of the genotyped individual and the disease history of parents and siblings when implementing GWAX (Table S19). The GWAX prevalence (i.e. prevalence of proxy cases) was generally more than double the parental prevalence and many times larger than the disease prevalence (see Table S44). For individuals with at least 1 sibling or who do not know or prefer not to answer how many brothers and/or sisters they have we assign using sibling and parental disease history; for individuals with 0 reported siblings we assign using parental disease history only (Table S19). For sex-specific diseases (breast and prostate cancer) the conditions are replaced with siblings of the relevant sex.

We considered the case-control status of the genotyped individual and the disease history of parents and siblings when implementing GWAX-2df (Table S19). We controlled for assessment center, genotype array, sex, age, age squared, and the first 20 principal components^21^. We restricted the GWAX-2df analysis to 672,292 genotyped SNPs to limit computational cost.

### Assessing calibration in UK Biobank

We assessed the calibration of each method using stratified LD score regression (S-LDSC) attenuation ratio^21,29–31^. We used stratified LD score regression with the baselineLD (v1.1) model to compute the attenuation ratio, defined as (S-LDSC intercept -1)/(mean *χ*^2^ - 1), for each set of association statistics^21,30,31^; the standard error of attenuation ratios is computed as s.e.(S-LDSC intercept)/(mean *χ*^2^ - 1). Regression SNPs for S-LDSC are HapMap Project Phase 3 (HapMap3) SNPs with an INFO score > 0.9, MAF > 0.01 and 0 < *P* ≤ 1. If the intercept is less than 1 S-LDSC reports that the attenuation ratio is less than 0; in these scenarios we set the attenuation ratio equal to 0 but still compute the standard error of the attenuation ratio as s.e.(S-LDSC intercept)/(mean *χ*^2^ - 1). We also compute the difference in attenuation ratios between various methods; the standard error of the difference is computed via block-jackknife.

We report two estimates for the mean attenuation ratio (and mean attenuation ratio difference) across all traits. Let *x*_*i*_ denote the attenuation ratio (or attenuation ratio difference) for trait *i* and *s*_*i*_ the reported jackknife standard error (for the ratio or the difference of ratios). We compute a simple average across traits, with standard error computed as 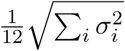. We also compute an inverse-variance weighted mean where the weights are determined by the variance of the GWAS attenuation ratio. The inverse-variance weighted estimate is 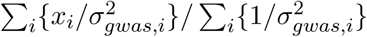, with standard error computed as 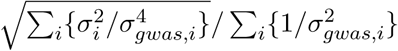.

### Assessing power in UK Biobank

We computed the number of independent loci as defined similar to ref.^21^. In detail, we applied PLINK’s LD clumping algorithm^41^ (see URLs) using LD computed in N=113,851 unrelated British individuals^42^ at million imputed SNPs with MAF>0.1% and INFO>0.6, employing a genome-wide significance threshold of p<5 * 10^−8^. We used a stringent 5Mb window and *R*^2^ threshold of 0.01 for LD clumping. We then collapsed independent signals that were within 100 kb of one another (as assessed using the top SNP from a LD-clump) into a single locus.

When computing the number of independent loci (as well as average *χ*^2^) we restrict to a MAF above a given threshold determined by prevalence in order to avoid an increased type I error in unbalanced case-control settings^21,28^. In the case of triallelic SNPs in which one variant allele has MAF > 0.1% and one variant allele has MAF < 0.1%, BOLT-LMM software retains association statistics for both variant alleles; we post-processed association statistics by manually removing all association statistics for variant alleles with MAF below a given threshold, thus removing the association statistic for the variant allele below the MAF threshold in these triallelic instances. The MAF threshold selected for rare diseases is a conservative choice for both GWAX and LT-FH as both of these methods have reduced risk of increased false-positives at lower MAF, due to higher case prevalence (or lower kurtosis for LT-FH) (Table S16).

The relative improvement for a method *M* versus a method *m* is defined as 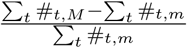 where #_*t,M*_ is the number of independent loci for trait *t* and method *M*; the standard error is computed via block jackknife of the genome (200 blocks).

### Assessing heritability and correlation in UK Biobank

We estimate observed-scale SNP-heritability 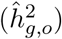 for each method by applying BOLT-REML^33^ to unrelated European individuals. Observed-scale SNP-heritability is converted to liability-scale SNP-heritability using the in-sample disease prevalence^43^. Genetic correlation is estimated using BOLT-REML^33^. The phenotypic correlation and correlation of −*log*_10_(*p*) (across all variants with a minimum MAF determined by case prevalence; Table S16) is estimated in `R`. For both genetic and phenotypic correlation we restrict attention to individuals with non-NA values for both methods being compared.

### Replication analysis

We conducted a replication analysis of loci identified by GWAS and/or LT-FH in independent non-UK Biobank data sets (4 diseases: coronary artery disease, type 2 diabetes, breast cancer, and prostate cancer) with publicly available summary statistics^44–47^(Table S25). The replication summary statistics are from studies consisting of predominantly non-UK Europeans and were always computed using GWAS (not LT-FH). For both genome-wide significant loci identified in UK Biobank by GWAS and genome-wide significant loci identified in UK Biobank by LT-FH, we computed the replication slope (the slope of a regression of standardized effect sizes in case-control replication data vs. UK Biobank discovery data). In detail, we restricted association results from linear regression on unrelated Europeans for GWAS and LT-FH to SNPs which (1) appear in the replication studies SNP set, (2) meet the MAF threshold for that trait from our main analysis, and (3) are significant in GWAS or LT-FH. We then LD-clumped this restricted set of association statistics (stringent 5Mb window and *R*^2^ threshold of 0.01 for LD clumping). For GWAS (and LT-FH, respectively) for each LD clump with at least one significant SNP, we selected the most significant SNP as the lead SNP. When an LD clump contained only SNPs that were significant for LT-FH (resp. GWAS), this was considered a LT-FH only locus (resp. GWAS only). The replication slope^34,35^ was then computed as the slope of a regression of standardized effect sizes of lead SNPs in case-control replication data vs. GWAS or LT-FH UK Biobank discovery data. Standardized effect sizes were computed as 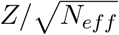, where *Z* denotes z-score, for GWAS (and GWAS replication data) *N*_*eff*_ was computed as 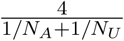, and for LT-FH *N*_*eff*_ was computed as *N*_*eff,GW AS*_ * *c* (where *c* is the relative effective sample size of LT-FH vs GWAS; see Table S22).

## Data availability

This study analyzed data from the UK Biobank, which is publicly available by application (see URLs). We have publicly released summary association statistics computed by applying our LT-FH method to UK Biobank data (see URLs).

## Code availability

We have publicly released open-source software implementing our LT-FH method (see URLs).

## Acknowledgements

We are grateful to L. O’Connor, O. Weissbrod, N. Zaitlen, G. Kichaev, and A. Gusev for helpful discussions, and E.M. Pedersen for computational suggestions. This research was funded by NIH grants R01 HG006399, R01 MH101244, R01 MH107649 and 5T32CA009337-32. This research was conducted using the UK Biobank Resource under Application #10438.

