## SupplementaryNote for "Combining case-control status and family history of disease increases association power"

**Supplementary Note**

**Equivalence to score test**

We consider an alternative, yet equivalent, parameterization of the liability threshold model where the liability is defined by $\phi = m + \epsilon$ where an individual is a case ($z = 1$) if and only if $\phi \geq 0$ and is a control otherwise ($z = 0$). In this parameterization $m$ determines the disease prevalence ($\Phi(-m) = P(x \geq -m)$ where $x \sim N(0,1)$). Again we note that if we are interested in testing one SNP, $g$, we assume that the effect size is small enough such that $\phi = m + \beta g + \epsilon$ where $\epsilon \sim N(0,1)$.

Consider testing one SNP of interest, $g$ in the trio setting where we have both parents' disease status, the child's disease status, as well as the child's genotype. We can write the prospective likelihood as a function of effect size $\beta$ as well as the genotypes of the trio (assuming we know the genotypes of the parents; genotypes standardized to have mean 0 and variance of 1). The underlying liabilities are,

$$\psi_{p1} \approx m + \beta g_{p1} + \underbrace{\epsilon_{p_1}}_{\sim N(0,1)};\psi_{p2} \approx m + \beta g_{p2} + \underbrace{\epsilon_{p_2}}_{\sim N(0,1)};\psi_{o} \approx m + \beta g_{o} + \underbrace{\epsilon_{o}}_{\sim N(0,1)},$$

where,

$$\underline{\epsilon} = \begin{bmatrix}\epsilon_{p_1}\\ \epsilon_{p_2}\\ \epsilon_o\end{bmatrix} \sim MVN_3\left(\begin{pmatrix}0\\0\\0\end{pmatrix}, \begin{pmatrix}1 & 0 & 0.5h^2\\ 0 & 1 & 0.5h^2\\ 0.5h^2 & 0.5h^2 & 1\end{pmatrix}\right).$$

Let

$$\begin{pmatrix}1 & 0 & 0.5h^2\\ 0 & 1 & 0.5h^2\\ 0.5h^2 & 0.5h^2 & 1\end{pmatrix} \equiv V; \qquad V^{-1} = Q.$$

Therefore,

$$\mathcal{L}(\beta, \boldsymbol{g}) \propto \int_{L_3}^{U_3}\int_{L_2}^{U_2}\int_{L_1}^{U_1} \exp\left\{-\frac{1}{2}\underline{\epsilon}^T Q \underline{\epsilon}\right\} d\epsilon_{p_1} d\epsilon_{p_2} d\epsilon_o = \int_{L_3}^{U_3}\int_{L_2}^{U_2}\int_{L_1}^{U_1} \exp\left\{-\frac{1}{2}\sum_{ij} Q_{ij}\epsilon_i\epsilon_j\right\} d\epsilon_{p_1} d\epsilon_{p_2} d\epsilon_o$$

where $L_i = -\infty, U_i = -m - \beta g_i$ for controls and $L_i = -m - \beta g_i, U_i = \infty$ for cases. Let $\epsilon_j = \epsilon_j^* - \beta g_j$. Thus,

$$\mathcal{L}(\beta, \boldsymbol{g}) \propto \int_{L_3^*}^{U_3^*}\int_{L_2^*}^{U_2^*}\int_{L_1^*}^{U_1^*} \exp\left\{-\frac{1}{2}\sum_{ij} Q_{ij}(\epsilon_i^* - \beta g_i)(\epsilon_j^* - \beta g_j)\right\} d\epsilon_{p_1}^* d\epsilon_{p_2}^* d\epsilon_o^*$$

where now $L_i^* = -\infty, U_i^* = -m$ for controls and $L_i^* = -m, U_i^* = \infty$ for cases. Let $\mathcal{P}1, \mathcal{P}2, \mathcal{O}$ denote the respective regions of integration for $\epsilon_{p_1}^*, \epsilon_{p_2}^*, \epsilon_o^*$. Therefore,

$$\frac{\delta}{\delta\beta}\log\mathcal{L}(\beta, \boldsymbol{g})|_{\beta=0} = \frac{\int\int\int_{\mathcal{P}1,\mathcal{P}2,\mathcal{O}} \left[\frac{1}{2}\sum_{ij} Q_{ij}(\epsilon_i^* g_j + \epsilon_j^* g_i)\right]\exp\left\{-\frac{1}{2}\sum_{ij} Q_{ij}\epsilon_i^*\epsilon_j^*\right\} d\epsilon_{p_1}^* d\epsilon_{p_2}^* d\epsilon_o^*}{\int\int\int_{\mathcal{P}1,\mathcal{P}2,\mathcal{O}} \exp\left\{-\frac{1}{2}\sum_{ij} Q_{ij}\epsilon_i^*\epsilon_j^*\right\} d\epsilon_{p_1}^* d\epsilon_{p_2}^* d\epsilon_o^*}$$

Note that $Q$ is symmetric so $Q_{ij} = Q_{ji}$ therefore,

$$\frac{1}{2}\sum_{ij} Q_{ij}(\epsilon_i^* g_j + \epsilon_j^* g_i) = \frac{1}{2}\left(\sum_{ij} Q_{ij}\epsilon_i^* g_j + \sum_{ij} Q_{ij}\epsilon_j^* g_i\right) = \sum_{ij} Q_{ij}\epsilon_i^* g_j$$

Therefore,

$$\frac{\delta}{\delta\beta}\log\mathcal{L}(\beta,\boldsymbol{g})|_{\beta=0}=\frac{\int\int\int_{\mathcal{P}1,\mathcal{P}2,\mathcal{O}}\left[\sum_i\epsilon_i^*\sum_j Q_{ij}g_j\right]\exp\left\{-\frac{1}{2}\sum_{ij}Q_{ij}\epsilon_i^*\epsilon_j^*\right\}d\epsilon_{p1}^*d\epsilon_{p2}^*d\epsilon_o^*}{\int\int\int_{\mathcal{P}1,\mathcal{P}2,\mathcal{O}}\exp\left\{-\frac{1}{2}\sum_{ij}Q_{ij}\epsilon_i^*\epsilon_j^*\right\}d\epsilon_{p1}^*d\epsilon_{p2}^*d\epsilon_o^*}.$$

In practice we only observe one genotype, assuming our known genotype is $g_o$:

$$\mathcal{L}(\beta)=\sum_{\boldsymbol{g}}P(\boldsymbol{g}|g_o)\mathcal{L}(\beta,\boldsymbol{g})$$

Therefore,

$$\frac{\delta}{\delta\beta}\mathcal{L}(\beta)=\sum_{\boldsymbol{g}}P(\boldsymbol{g}|g_o)\frac{\delta}{\delta\beta}\mathcal{L}(\beta,\boldsymbol{g})$$

or,

$$\frac{\delta}{\delta\beta}\mathcal{L}(\beta)|_{\beta=0}=\sum_{\boldsymbol{g}}P(\boldsymbol{g}|g_o)\left\{\frac{\delta}{\delta\beta}\mathcal{L}(\beta,\boldsymbol{g})|_{\beta=0}\right\}$$

By the linearity of the log likelihood in $\boldsymbol{g}$ we find,

$$\frac{\delta}{\delta\beta}\mathcal{L}(\beta)|_{\beta=0}=\frac{\delta}{\delta\beta}\mathcal{L}(\beta,E(\boldsymbol{g}|g_o))|_{\beta=0}$$

Note in the case of offspring-parents we obtain, $E(\boldsymbol{g}|g_o)=(g_o/2,g_o/2,g_o)$ therefore one can show,

$$\frac{\delta}{\delta\beta}\log\mathcal{L}(\beta)|_{\beta=0}=\frac{g_o}{1-(h^2)^2/2}E\left[(\frac{1}{2}-h^2/2)\epsilon_{p_1}^*+(\frac{1}{2}-h^2/2)\epsilon_{p_2}^*+(1-h^2/2)\epsilon_o^*|z_o,z_{p_1},z_{p_2}\right],$$

where $\underline{\epsilon^*}\sim\underline{\epsilon}$. Note that the distribution of $\epsilon_i^*$ is distributed the same as $\epsilon_i$ when $\beta\equiv 0$. It follows the Score statistic for a collection of trios is equal to the square of

$$\sum_i\frac{g_{o,i}}{1-(h^2)^2/2}E\left[(\frac{1}{2}-h^2/2)\epsilon_{p_1}^*+(\frac{1}{2}-h^2/2)\epsilon_{p_2}^*+(1-h^2/2)\epsilon_o^*|z_{o,i},z_{p_1,i},z_{p_2,i}\right],$$

divided by its empirical variance, which is equivalent to computing the number of samples times the squared correlation between $g_o$ and $\frac{1}{1-(h^2)^2/2}E\left[(\frac{1}{2}-h^2/2)\epsilon_{p_1}^*+(\frac{1}{2}-h^2/2)\epsilon_{p_2}^*+(1-h^2/2)\epsilon_o^*|z_o,z_{p_1},z_{p_2}\right]$ (generalizing the Armitage trend test[1]).

We posit that

$$\frac{1}{h^2}E\left[\epsilon_{o,g}|z_o,z_{p_1},z_{p_2}\right]=\frac{1}{1-(h^2)^2/2}E\left[(\frac{1}{2}-h^2/2)\epsilon_{p_1}^*+(\frac{1}{2}-h^2/2)\epsilon_{p_2}^*+(1-h^2/2)\epsilon_o^*|z_o,z_{p_1},z_{p_2}\right]$$

and therefore computing the number of samples times the squared correlation between $g_o$ and posterior mean genetic liability is equivalent to the score test.

Noting that $z_o,z_{p_1},z_{p_2}\implies\epsilon_o\in\mathcal{O},\epsilon_{p1}\in\mathcal{P}1,\epsilon_{p2}\in\mathcal{P}2$ we consider,

$$E\left[\epsilon_{o,g}|\epsilon_{o,g}+\epsilon_{o,e}\in\mathcal{O},\epsilon_{p1}\in\mathcal{P}1,\epsilon_{p2}\in\mathcal{P}2\right]$$

where,

$$\epsilon_o = \epsilon_{o,g} + \epsilon_{o,e}; \qquad \begin{pmatrix} \epsilon_{o,e} \\ \epsilon_{o,g} \\ \epsilon_{p1} \\ \epsilon_{p2} \end{pmatrix} \sim N\left(\begin{pmatrix} 0 \\ 0 \\ 0 \\ 0 \end{pmatrix}, \begin{pmatrix} 1-h^2 & 0 & 0 & 0 \\ 0 & h^2 & 0.5h^2 & 0.5h^2 \\ 0 & 0.5h^2 & 1 & 0 \\ 0 & 0.5h^2 & 0 & 1 \end{pmatrix}\right)$$

We can see,

$$\begin{pmatrix} \epsilon_{o,g} \\ \epsilon_o \\ \epsilon_{p1} \\ \epsilon_{p2} \end{pmatrix} = \begin{pmatrix} \epsilon_{o,g} \\ \epsilon_{o,g} + \epsilon_{o,e} \\ \epsilon_{p1} \\ \epsilon_{p2} \end{pmatrix} = \begin{bmatrix} 0 & 1 & 0 & 0 \\ 1 & 1 & 0 & 0 \\ 0 & 0 & 1 & 0 \\ 0 & 0 & 0 & 1 \end{bmatrix} \begin{pmatrix} \epsilon_{o,e} \\ \epsilon_{o,g} \\ \epsilon_{p1} \\ \epsilon_{p2} \end{pmatrix} \rightarrow \begin{pmatrix} \epsilon_{o,g} \\ \epsilon_o \\ \epsilon_{p1} \\ \epsilon_{p2} \end{pmatrix} \sim N\left(\mathbf{0}, \begin{pmatrix} h^2 & h^2 & 0.5h^2 & 0.5h^2 \\ h^2 & 1 & 0.5h^2 & 0.5h^2 \\ 0.5h^2 & 0.5h^2 & 1 & 0 \\ 0.5h^2 & 0.5h^2 & 0 & 1 \end{pmatrix}\right)$$

Thus $\epsilon_{o,g} | \epsilon_o, \epsilon_{p1}, \epsilon_{p2} \sim N(\mu^*, \Sigma^*)$ with,

$$\mu^* = \frac{1}{1 - 2(0.5h^2)^2} \left\{ (h^2 - 2(0.5h^2)^2)\epsilon_o + (0.5h^2 - 0.5(h^2)^2)\epsilon_{p1} + (0.5h^2 - 0.5(h^2)^2)\epsilon_{p2} \right\}$$

Therefore, denoting $(\epsilon_o, \epsilon_{p1}, \epsilon_{p2}) \in (\mathcal{O}, \mathcal{P}1, \mathcal{P}2)$ by $\boldsymbol{\epsilon} \in \boldsymbol{\mathcal{F}}$

$$\begin{aligned} \frac{1}{h^2} E\left[\epsilon_{o,g} | \boldsymbol{\epsilon} \in \boldsymbol{\mathcal{F}}\right] &= \frac{1}{h^2} E\left\{ E\left[\epsilon_{o,g} | \boldsymbol{\epsilon} \in \boldsymbol{\mathcal{F}}, \epsilon_o, \epsilon_{p1}, \epsilon_{p2}\right] | \boldsymbol{\epsilon} \in \boldsymbol{\mathcal{F}} \right\} \\ &= \frac{2 - h^2}{2 - (h^2)^2} E[\epsilon_o | \boldsymbol{\epsilon} \in \boldsymbol{\mathcal{F}}] + \frac{1 - h^2}{2 - (h^2)^2} E[\epsilon_{p1} | \boldsymbol{\epsilon} \in \boldsymbol{\mathcal{F}}] + \frac{1 - h^2}{2 - (h^2)^2} E[\epsilon_{p2} | \boldsymbol{\epsilon} \in \boldsymbol{\mathcal{F}}], \end{aligned}$$

thus we have shown,

$$\frac{1}{h^2} E\left[\epsilon_{o,g} | z_o, z_{p_1}, z_{p_2}\right] = \frac{1}{1 - (h^2)^2/2} E\left[ (\frac{1}{2} - h^2/2)\epsilon_{p_1} + (\frac{1}{2} - h^2/2)\epsilon_{p_2} + (1 - h^2/2)\epsilon_o | z_o, z_{p_1}, z_{p_2} \right]$$

as desired.

**2-df F-test for GWAX-2df**

Rather than conducting a Pearson's chi-square test we perform the following two regressions:

$$\begin{aligned} \boldsymbol{g} &= \boldsymbol{X\gamma} + \beta_1 \boldsymbol{y} + \beta_2 \boldsymbol{y}^*; \qquad &(F) \\ \boldsymbol{g} &= \boldsymbol{X\tilde{\gamma}}, &(R) \end{aligned}$$

where $\boldsymbol{g}$ is a vector of the genotypes of interest, $\boldsymbol{y}$ denotes case status, $\boldsymbol{y}^*$ denotes proxy-case status (unaffected individual with a family history of disease), and $\boldsymbol{X}$ represents other covariates (consider $p$ covariates; thus as we include an intercept both $\boldsymbol{\gamma}, \tilde{\boldsymbol{\gamma}}$ have $p+1$ terms). We can then test the null hypothesis that $\beta_1 = \beta_2 = 0$ with the following F statistic:

$$F^* = \frac{(SSE(R) - SSE(F)) * df_F}{(df_R - df_F) * SSE(F)} \sim F_{df_R - df_F, df_F}.$$

where $df_R, df_F$ are the degrees of freedom associated with the reduced and full model error sums of squares. From $(F), (R)$ we can see, $df_R = n - (p+1), df_F = n - (p+1+2)$. Also note that by properties of F distributions we know,

$$\lim_{df_F \to \infty} (df_R - df_F) F^* = \lim_{df_F \to \infty} \frac{(SSE(R) - SSE(F)) * df_F}{SSE(F)} \sim \chi^2_{df_R - df_F} = \chi^2_2.$$

We can find, letting $\boldsymbol{X}^* = \begin{bmatrix}\boldsymbol{X} & \boldsymbol{y} & \boldsymbol{y}^*\end{bmatrix}$,

$$SSE(F) = \boldsymbol{g}^T \left[\boldsymbol{I} - \boldsymbol{X}^*(\boldsymbol{X}^{*T}\boldsymbol{X}^*)^{-1}\boldsymbol{X}^{*T}\right]\boldsymbol{g};$$
$$SSE(R) = \boldsymbol{g}^T \left[\boldsymbol{I} - \boldsymbol{X}(\boldsymbol{X}^{T}\boldsymbol{X})^{-1}\boldsymbol{X}^{T}\right]\boldsymbol{g}.$$

In simulations without covariates this proves to be almost identical to the Pearson's Chi-Square test on the $3 \times 2$ table but the above formulation provides a way to control for various covariates in the data application.

**Variance Heterogeneity**

We compared an unweighted binary-scale method to a binary-scale method that incorporates weights equal to the inverse of the genetic predictor error variance. We follow the derivation in ref.[2]. In detail, we assumed $\boldsymbol{Y} = \beta\boldsymbol{X} + \boldsymbol{Z}\boldsymbol{u} + \boldsymbol{e}$ where $\boldsymbol{X} = \boldsymbol{1}^T$, $\boldsymbol{u} \sim (0, G) \perp \boldsymbol{e} \sim (0, R)$, $G = \sigma_A^2 A$ (A is the additive genetic relationship matrix) and $R = \sigma_e^2\boldsymbol{I}$. Thus, $\boldsymbol{Z}\boldsymbol{u}$ represents the genetic component of $\boldsymbol{Y}$ ($\epsilon_g$ on the binary scale). In our setting, in which we do not have repeated measures on individuals, $\boldsymbol{Z} = \boldsymbol{I}$. Let $V = \boldsymbol{Z}G\boldsymbol{Z}^T + R = G + R$ and $\beta = (\boldsymbol{X}^T V^{-1}\boldsymbol{X})^{-1}\boldsymbol{X}^T V^{-1}\boldsymbol{Y}$. Consider $N$ individuals, of which $C\%$ have case-control status only and $1 - C\%$ have case-control status and parental history. In this scenario, we let

$$\boldsymbol{Y} = \begin{pmatrix} Y_{1,p1} & Y_{1,p2} & Y_{1,o} & \cdots & Y_{(1-C\%)N,p1} & Y_{(1-C\%)N,p2} & Y_{(1-C\%)N,o} & Y_{(1-C\%)N+1,o} & \cdots & Y_{N,o}\end{pmatrix}^T$$

Note that

$$V = \begin{pmatrix} \boldsymbol{V_1} & \boldsymbol{0} \\ \boldsymbol{0} & \boldsymbol{V_2}\end{pmatrix}; \qquad \boldsymbol{V_1} = \begin{pmatrix} F & 0 & \cdots & 0 \\ 0 & F & \cdots & 0 \\ \vdots & \vdots & \ddots & \vdots \\ 0 & 0 & \cdots & F \end{pmatrix}, F = \begin{pmatrix} 1 & 0 & 0.5\sigma_A^2 \\ 0 & 1 & 0.5\sigma_A^2 \\ 0.5\sigma_A^2 & 0.5\sigma_A^2 & 1 \end{pmatrix}, \boldsymbol{V_2} = I,$$

where $\boldsymbol{V_1}$ is a $N * (1 - C\%) \times N * (1 - C\%)$ matrix and $\boldsymbol{V_2}$ is $N * C\% \times N * C\%$. It follows that

$$\hat{\beta} = \frac{\sum F_{\cdot 1}^{-1} \sum_1^{(1-C\%)N} Y_{i,p1} + \sum F_{\cdot 2}^{-1} \sum_1^{(1-C\%)N} Y_{i,p2} + \sum F_{\cdot 3}^{-1} \sum_1^{(1-C\%)N} Y_{i,o} + \sum_{(1-C\%)N+1}^{N} Y_{i,o}}{\sum F_{\cdot\cdot}^{-1} * ((1 - C\%)N) + (C\%N)}$$

Letting $\tilde{\boldsymbol{Y}} = \boldsymbol{Y} - \boldsymbol{X}\hat{\beta}$, we have

$$\hat{u}_i = \sigma_A^2 \begin{cases} (\frac{F_{11}^{-1}}{2} + \frac{F_{21}^{-1}}{2} + F_{31}^{-1})\tilde{Y}_{i,p1} + (\frac{F_{12}^{-1}}{2} + \frac{F_{22}^{-1}}{2} + F_{32}^{-1})\tilde{Y}_{i,p2} + (\frac{F_{13}^{-1}}{2} + \frac{F_{23}^{-1}}{2} + F_{33}^{-1})\tilde{Y}_{i,o} & \text{cc +FH} \\ \tilde{Y}_i & \text{cc only} \end{cases}$$

Now we can estimate the genetic predictor error variance ($var(\hat{u} - u)$) as the diagonal of $C_{22}$ in the following:

$$\begin{pmatrix} \boldsymbol{X}^T R^{-1}\boldsymbol{X} & \boldsymbol{X}^T R^{-1}\boldsymbol{Z} \\ \boldsymbol{Z}^T R^{-1}\boldsymbol{X} & \boldsymbol{Z}^T R^{-1}\boldsymbol{Z} + G^{-1} \end{pmatrix}^{-1} = \begin{pmatrix} C_{11} & C_{12} \\ C_{21} & C_{22} \end{pmatrix}.$$

It follows that the diagonals of $C_{22}$ are;

$$\begin{cases} FH_{33} + \frac{N(3-2C\%)/\sigma_e^2 - ((1-C\%)N\sum FH_{\cdot\cdot} + C\%N\sigma_A^2\sigma_e^2)/(\sigma_e^2)^2}{(\sigma_e^2)^2} \sum FH_{3\cdot} \sum FH_{\cdot 3} & \text{cc +FH} \\ \sigma_A^2\sigma_e^2 + \frac{N(3-2C\%)/\sigma_e^2 - ((1-C\%)N\sum FH_{\cdot\cdot} + C\%N\sigma_A^2\sigma_e^2)/(\sigma_e^2)^2}{(\sigma_e^2)^2} (\sigma_A^2\sigma_e^2)^2 & \text{cc only,} \end{cases}$$

where

$$FH = \left( \frac{1}{\sigma_e^2} \boldsymbol{I}_3 + \frac{1}{\sigma_A^2} \begin{pmatrix} 1 & 0 & 0.5 \\ 0 & 1 & 0.5 \\ 0.5 & 0.5 & 1 \end{pmatrix}^{-1} \right)^{-1} .$$

The unweighted binary-scale method uses $\hat{\boldsymbol{u}}$ as the response variable; the weighted binary-scale method uses $\hat{\boldsymbol{u}}$ as the response variable but has weights equal to the inverse of the predictor error variance ($Var(\hat{\boldsymbol{u}} - \boldsymbol{u})$).

There are complexities that arise when incorporating weights in the case of sibling history (as collected in UK Biobank). If sibling history is provided as a binary response as in UK Biobank data, there is no straightforward method to estimate $\hat{\boldsymbol{u}}$ and therefore $Var(\hat{\boldsymbol{u}} - \boldsymbol{u})$. Thus in this scenario, we would have to assume that exactly one sibling (rather than at least one sibling) is affected in the case of positive sibling history. Making this assumption when applying the (analytical) PA formula resulted in a less powerful method than our (Monte Carlo) LT-FH method, as a function of number of siblings and disease prevalence (Table S11 and Table S34). There are also complexities that arise when using BOLT-LMM. BOLT-LMM software currently does not accept user-specified weights, therefore in order to perform a weighted analysis the software would have to be modified. One could also modify the response (e.g. divide $\hat{u}$ by the predictor error variance) and then perform an unweighted analysis, however this changes interpretation of the regression parameters.

Within UK Biobank, there is a limited amount of variance heterogeneity actually present (Table S18). Most individuals with reported case-control status also have data on both parents' history (average of 87% for non-sex-specific diseases), confirming that our simulations in which 10% of the individuals have case-control status data only and 90% of the individuals have case-control status plus parental history information represent a realistic scenario.

Due to (1) the limited benefit of incorporating weights to account for variance heterogeneity in simulations using parental history and linear regression (Table S13), (2) the complexities of incorporating weights in the case of sibling history or BOLT-LMM (see above), and (3) the limited amount of variance heterogeneity actually present in the UK Biobank (Table S18), we elected to not to further investigate modeling variance heterogeneity in real UK Biobank traits.

Table S1: **Results of simulations with default parameter settings.** Number of individuals ($N$) and number of SNPs ($M$) is $100K$; $h^2 = 0.5$; disease prevalence is 5%; we assume perfect knowledge of $h^2/K$ when implementing LT-FH; no environmental correlation between parents-offspring; we consider 10 simulation replicates. * denotes 2df test statistics that are not directly comparable to 1df test statistics. The standard error of the mean $\chi^2$ for null and causal SNPs and the standard error for the power ($\sqrt{\hat{p}(1-\hat{p})/n}$) are reported in parentheses.

| | GWAS | GWAX | GWAX-2df | LT-FH |
|---|---|---|---|---|
| $\bar{\chi}^2_{null}$ (SEM) | 1.00(0.001) | 1.00(0.001) | 2.00(0.002)* | 1.00(0.001) |
| $\bar{\chi}^2_{causal}$ (SEM) | 24.72(0.15) | 27.34(0.15) | 33.75(0.17)* | 33.24(0.17) |
| Power | 0.282(0.006) | 0.371(0.007) | 0.460(0.007) | 0.576(0.007) |

Table S2: **Type 1 errors of simulations with default parameter settings.** We report the percentage of type 1 errors (false positive rate) for simulations with default parameter settings at various $\alpha$ levels. Results are based on 10 simulation replicates. The total number of null SNPs across replicates is 995,000; the standard error of the false positive rate ($\sqrt{\hat{p}(1-\hat{p})/n}$) are reported in parentheses.

| $\alpha$ | GWAS | GWAX | GWAX-2df | LT-FH |
|---|---|---|---|---|
| $5 * 10^{-2}$ | 0.05(0.00022) | 0.05(0.00022) | 0.05(0.00022) | 0.05(0.00022) |
| $5 * 10^{-3}$ | 0.005(7.3e-05) | 0.005(7.2e-05) | 0.0049(7.2e-05) | 0.0049(7.2e-05) |
| $5 * 10^{-4}$ | 0.00054(2.4e-05) | 0.00044(2.2e-05) | 0.00051(2.3e-05) | 0.00049(2.3e-05) |
| $5 * 10^{-5}$ | 5.9e-05(7.9e-06) | 4.4e-05(6.8e-06) | 4.7e-05(7.1e-06) | 4.7e-05(7.1e-06) |
| $5 * 10^{-6}$ | 5e-06(2.3e-06) | 6e-06(2.5e-06) | 7e-06(2.7e-06) | 3e-06(1.8e-06) |

Table S3: **Results of simulations at different values of prevalence.** Number of individuals ($N$) and number of SNPs ($M$) is $100K$; $h^2 = 0.5$; we assume perfect knowledge of $h^2/K$ when implementing LT-FH; no environmental correlation between parents-offspring; we consider 10 simulation replicates. For these simulation scenarios we modify the number of causal SNPs such that the GWAS power (and average GWAS causal) is approximately the across all scenarios for comparative reasons. * denotes 2df test statistics that are not directly comparable to 1df test statistics. The standard error of the mean $\chi^2$ for null and causal SNPs and the standard error for the power ($\sqrt{\hat{p}(1-\hat{p})/n}$) are reported in parentheses.

| | GWAS | GWAX | GWAX-2df | LT-FH |
|---|---|---|---|---|
| Disease prevalence 1% (180 causal SNPs) | | | | |
| $\bar{\chi}^2_{null}$ | 1.00(0.001) | 1.00(0.001) | 2.00(0.002)* | 1.00(0.001) |
| $\bar{\chi}^2_{causal}$ | 24.07(0.26) | 29.50(0.29) | 34.91(0.32)* | 34.00(0.32) |
| Power | 0.265(0.010) | 0.456(0.012) | 0.492(0.012) | 0.584(0.012) |
| Disease prevalence 5% (500 causal SNPs) | | | | |
| $\bar{\chi}^2_{null}$ | 1.00(0.001) | 1.00(0.001) | 2.00(0.002)* | 1.00(0.001) |
| $\bar{\chi}^2_{causal}$ | 24.72(0.15) | 27.34(0.15) | 33.75(0.17)* | 33.24(0.17) |
| Power | 0.282(0.006) | 0.371(0.007) | 0.460(0.007) | 0.576(0.007) |
| Disease prevalence 25% (1150 causal SNPs) | | | | |
| $\bar{\chi}^2_{null}$ | 1.00(0.001) | 1.00(0.001) | 2.00(0.002)* | 1.00(0.001) |
| $\bar{\chi}^2_{causal}$ | 24.71(0.09) | 20.09(0.08) | 30.11(0.10)* | 31.10(0.10) |
| Power | 0.282(0.004) | 0.140(0.003) | 0.347(0.004) | 0.513(0.005) |

Table S4: **Results of simulations at different values of number of causal SNPs.** Number of individuals ($N$) and number of SNPs ($M$) is $100K$; $h^2 = 0.5$; $K = 0.05$; we assume perfect knowledge of $h^2/K$ when implementing LT-FH; no environmental correlation between parents-offspring; we consider 10 simulation replicates. * denotes 2df test statistics that are not directly comparable to 1df test statistics. The standard error of the mean $\chi^2$ for null and causal SNPs and the standard error for the power ($\sqrt{\hat{p}(1-\hat{p})/n}$) are reported in parentheses.

| | GWAS | GWAX | GWAX-2df | LT-FH |
|---|---|---|---|---|
| 250 causal SNPs | | | | |
| $\bar{\chi}^2_{null}$ | 1.00(0.001) | 1.00(0.001) | 2.00(0.002)* | 1.00(0.001) |
| $\bar{\chi}^2_{causal}$ | 49.10(0.31) | 54.29(0.31) | 66.43(0.36)* | 66.41(0.37) |
| Power | 0.914(0.006) | 0.961(0.004) | 0.984(0.003) | 0.994(0.002) |
| 500 causal SNPs | | | | |
| $\bar{\chi}^2_{null}$ | 1.00(0.001) | 1.00(0.001) | 2.00(0.002)* | 1.00(0.001) |
| $\bar{\chi}^2_{causal}$ | 24.72(0.15) | 27.34(0.15) | 33.75(0.17)* | 33.24(0.17) |
| Power | 0.282(0.006) | 0.371(0.007) | 0.460(0.007) | 0.576(0.007) |
| 750 causal SNPs | | | | |
| $\bar{\chi}^2_{null}$ | 1.00(0.001) | 1.00(0.001) | 2.00(0.002)* | 1.00(0.001) |
| $\bar{\chi}^2_{causal}$ | 16.65(0.10) | 18.50(0.10) | 23.06(0.11)* | 22.41(0.11) |
| Power | 0.076(0.003) | 0.105(0.004) | 0.139(0.004) | 0.213(0.005) |

Table S5: **Results of simulations at different values of heritability.** Number of individuals ($N$) and number of SNPs ($M$) is $100K$; $K = 0.05$; we assume perfect knowledge of $h^2/K$ when implementing LT-FH; no environmental correlation between parents-offspring; we consider 10 simulation replicates. * denotes 2df test statistics that are not directly comparable to 1df test statistics. The standard error of the mean $\chi^2$ for null and causal SNPs and the standard error for the power ($\sqrt{\hat{p}(1-\hat{p})/n}$) are reported in parentheses. For these simulation scenarios we modify the underlying heritability of the disease; we modify $C$ such that the GWAS power (and average GWAS causal) is approximately the across all scenarios for comparative reasons.

| | GWAS | GWAX | GWAX-2df | LT-FH |
|---|---|---|---|---|
| $h_l^2 = 0.25$ (250 causal SNPs) | | | | |
| $\bar{\chi}^2_{null}$ | 1.00(0.001) | 1.00(0.001) | 2.00(0.002)* | 1.00(0.001) |
| $\bar{\chi}^2_{causal}$ | 24.85(0.21) | 29.19(0.22) | 35.36(0.25)* | 35.09(0.25) |
| Power | 0.287(0.009) | 0.442(0.010) | 0.514(0.010) | 0.643(0.010) |
| $h_l^2 = 0.50$ (500 causal SNPs) | | | | |
| $\bar{\chi}^2_{null}$ | 1.00(0.001) | 1.00(0.001) | 2.00(0.002)* | 1.00(0.001) |
| $\bar{\chi}^2_{causal}$ | 24.72(0.15) | 27.34(0.15) | 33.75(0.17)* | 33.24(0.17) |
| Power | 0.282(0.006) | 0.371(0.007) | 0.460(0.007) | 0.576(0.007) |
| $h_l^2 = 0.75$ (750 causal SNPs) | | | | |
| $\bar{\chi}^2_{null}$ | 1.00(0.001) | 1.00(0.001) | 2.00(0.002)* | 1.00(0.001) |
| $\bar{\chi}^2_{causal}$ | 24.36(0.12) | 25.01(0.12) | 31.81(0.13)* | 31.03(0.13) |
| Power | 0.271(0.005) | 0.293(0.005) | 0.394(0.006) | 0.506(0.006) |

Table S6: **Results of simulations with misspecified prevalence or heritability.** Number of individuals ($N$) and number of SNPs ($M$) is $100K$; $h_l^2 = 0.5$, $K = 0.05$; no environmental correlation between parents and offspring; we consider 10 simulation replicates. * denotes 2df test statistics that are not directly comparable to 1df test statistics. The standard error of the mean $\chi^2$ for null and causal SNPs and the standard error for the power ($\sqrt{\hat{p}(1-\hat{p})/n}$) are reported in parentheses. The same genotype-phenotype information is used across these scenarios with an LT-FH value calculated from a misspecified model (misspecified $h^2$ or $K$). LT-FH $\Delta$ the difference between the $\chi^2$ value from models in which we misspecify $h^2$ or $K$ and the $\chi^2$ value when $h^2$ and $K$ are correctly specified. Although we find that the power is negligibly affected, we see significant (small) decreases in the average causal $\chi^2$ when heritability and prevalence are misspecified in the LT-FH method.

| | GWAS | GWAX | GWAX-2df | LT-FH | LT-FH $\Delta$ |
|---|---|---|---|---|---|
| $\bar{\chi}^2_{null}$ | 1.00(0.001) | 1.00(0.001) | 2.00(0.002)* | 1.00(0.001) | |
| $\bar{\chi}^2_{causal}$ | 24.72(0.15) | 27.34(0.15) | 33.75(0.17)* | 33.24(0.17) | |
| Power | 0.282(0.006) | 0.371(0.007) | 0.460(0.007) | 0.576(0.007) | |
| Assume $K = 0.025$ | | | | | |
| $\bar{\chi}^2_{null}$ | - | - | - | 1.00(0.001) | |
| $\bar{\chi}^2_{causal}$ | - | - | - | 33.23(0.17) | -0.01(0.002) |
| Power | | | | 0.576(0.007) | |
| Assume $K = 0.075$ | | | | | |
| $\bar{\chi}^2_{null}$ | - | - | - | 1.00(0.001) | |
| $\bar{\chi}^2_{causal}$ | - | - | - | 33.23(0.17) | -0.002(0.002) |
| Power | | | | 0.576(0.007) | |
| Assume $h_l^2 = 0.25$ | | | | | |
| $\bar{\chi}^2_{null}$ | - | - | - | 1.00(0.001) | |
| $\bar{\chi}^2_{causal}$ | - | - | - | 33.19(0.17) | -0.044(0.007) |
| Power | | | | 0.577(0.007) | |
| Assume $h_l^2 = 0.75$ | | | | | |
| $\bar{\chi}^2_{null}$ | - | - | - | 1.00(0.001) | |
| $\bar{\chi}^2_{causal}$ | - | - | - | 33.17(0.17) | -0.062(0.007) |
| Power | | | | 0.573(0.007) | |

Table S7: **Results of simulations with shared environment.** Number of individuals ($N$) and number of SNPs ($M$) is $100K$; $h_l^2 = 0.5$, $K = 0.05$; we assume perfect knowledge of $h^2/K$ when implementing LT-FH; we consider 10 simulation replicates. * denotes 2df test statistics that are not directly comparable to 1df test statistics. The standard error of the mean $\chi^2$ for null and causal SNPs and the standard error for the power ($\sqrt{\hat{p}(1-\hat{p})/n}$) are reported in parentheses.

| | GWAS | GWAX | GWAX-2df | LT-FH |
|---|---|---|---|---|
| No environmental correlation | | | | |
| $\bar{\chi}^2_{null}$ | 1.00(0.001) | 1.00(0.001) | 2.00(0.002)* | 1.00(0.001) |
| $\bar{\chi}^2_{causal}$ | 24.72(0.15) | 27.34(0.15) | 33.75(0.17)* | 33.24(0.17) |
| Power | 0.282(0.006) | 0.371(0.007) | 0.460(0.007) | 0.576(0.007) |
| Environmental correlation = $0.5(0.5h^2)$ | | | | |
| $\bar{\chi}^2_{null}$ | 1.00(0.001) | 1.00(0.001) | 2.00(0.002) | 1.00(0.001) |
| $\bar{\chi}^2_{causal}$ | 24.39(0.15) | 25.16(0.15) | 31.93(0.17) | 31.16(0.17) |
| Power | 0.279(0.006) | 0.295(0.006) | 0.403(0.007) | 0.508(0.007) |

Table S8: **Results of simulations with two forms of family history reporting bias.** Number of individuals (N) and number of SNPs (M) is 100K; $h^2 = 0.5$; disease prevalence is 5%; we assume perfect knowledge of $h^2/K$ when implementing LT-FH; no environmental correlation between parents-offspring; we consider 10 simulation replicates. The standard error of the mean $\chi^2$ for null and causal SNPs and the standard error for the power are reported in parentheses. We report results of LT-FH when all individuals report correct family history information for comparison reasons. We run two separate scenarios investigating family history reporting bias: (1) every control has missing family history (Control FH missing) and (2) every control reports both parents as unaffected (Control report 0FH; investigate impact of recall bias). We note that for GWAX, when every control has missing family history (1) the analysis has no controls and no results can be obtained (as individuals with missing family history are removed) and when every control reports both parents as unaffected (2) the results are equivalent to GWAS.

| | GWAS | LT-FH | Control FH missing | Control report 0FH |
|---|---|---|---|---|
| $\chi^2_{causal}$ (SEM) | 24.72 (0.1) | 33.24 (0.2) | 25.1 (0.1) | 25.1 (0.1) |
| Power | 0.28 (0.006) | 0.58 (0.007) | 0.30 (0.006) | 0.29 (0.006) |
| $\chi^2_{null}$ (SEM) | 1 (0.001) | 1 (0.001) | 1 (0.001) | 1 (0.001) |

Table S9: **Results of PA formula in simulations.** Number of individuals ($N$) and number of SNPs ($M$) is 100K; $h^2 = 0.5$; we assume perfect knowledge of $h^2/K$ when implementing LT-FH; no environmental correlation between parents-offspring; we consider 10 simulation replicates. For these simulation scenarios we modify the number of causal SNPs such that the GWAS power (and average GWAS causal) is approximately the across all scenarios for comparative reasons (as in Table S3).We compute LT-PA (the LT-FH phenotype using selection theory (not Monte-Carlo integration)) and LT-PA$_{binary}$ (LT-PA phenotype on the binary scale (uses observed scale heritability and normalized phenotypes)). The mean correlation between LT-FH and LT-PA across the 10 simulations is 0.9999963 and between LT-PA and LT-PA$_{binary}$ is 0.9971677 when prevalence is 5% (similar phenotypic correlations are seen for K=1% and 25%).

| | LT-FH | LT-PA | LT-PA$_{binary}$ |
|---|---|---|---|
| | | K=0.01 | |
| $\chi^2_{causal}$ (SEM) | 34 (0.316) | 34 (0.316) | 33.9 (0.316) |
| Power | 0.584 (0.01) | 0.584 (0.01) | 0.587 (0.01) |
| $\chi^2_{null}$ (SEM) | 1 (0.001) | 1 (0.001) | 1 (0.001) |
| | | K=0.05 | |
| $\chi^2_{causal}$ (SEM) | 33.2 (0.169) | 33.2 (0.169) | 33.1 (0.169) |
| Power | 0.576 (0.007) | 0.576 (0.007) | 0.574 (0.007) |
| $\chi^2_{null}$ (SEM) | 1.001 (0.001) | 1.001 (0.001) | 1.001 (0.001) |
| | | K=0.25 | |
| $\chi^2_{causal}$ (SEM) | 31.1 (0.104) | 31.1 (0.104) | 31.1 (0.104) |
| Power | 0.513 (0.005) | 0.514 (0.005) | 0.512 (0.005) |
| $\chi^2_{null}$ (SEM) | 0.998 (0.001) | 0.998 (0.001) | 0.998 (0.001) |

Table S10: **Results of simulations with sibling history.** Number of individuals (N) is 100K and number of SNPs (M) is 1000 (1650 for $K = 0.25$ scenario); we assume perfect knowledge of $h^2$ and $K$ when implementing LT-FH; no environmental correlation between parents, offspring, or siblings; we consider 10 simulation replicates. The standard error of the mean $\chi^2$ for null and causal SNPs and the standard error for the power are reported in parentheses. For these simulation scenarios we modify the underlying prevalence of the disease ($K = 0.01$, 0.05, and 0.25) as well as the number of siblings each individual has ($n_s = 1, 2, 5, 10$); we modify C such that the GWAS power (and average GWAS causal $\chi^2$) is approximately the across all scenarios for comparative reasons. For both GWAX and LT-FH we assume that as in UK Biobank the number of affected siblings is not available; we assume all that is known is whether none or at least one sibling is affected.

| | | $n_s = 1$ | | $n_s = 2$ | | $n_s = 5$ | | $n_s = 10$ | |
|---|---|---|---|---|---|---|---|---|---|
| | GWAS | GWAX | LT-FH | GWAX | LT-FH | GWAX | LT-FH | GWAX | LT-FH |
| | K=0.01 | | | | | | | | |
| $\chi^2_{causal}$ (SEM) | 24 (0.3) | 32.86 (0.3) | 38.13 (0.3) | 36.19 (0.3) | 42.08 (0.4) | 45.35 (0.4) | 52.6 (0.4) | 57.49 (0.4) | 66.14 (0.5) |
| Power | 0.26 (0.01) | 0.53 (0.01) | 0.72 (0.01) | 0.66 (0.01) | 0.8 (0.009) | 0.85 (0.008) | 0.93 (0.006) | 0.97 (0.004) | 0.99 (0.002) |
| $\chi^2_{null}$ (SEM) | 1.01 (0.02) | 1.03 (0.02) | 1.02 (0.02) | 1.02 (0.02) | 1.01 (0.02) | 1.01 (0.02) | 1.01 (0.02) | 0.99 (0.02) | 1 (0.02) |
| | K=0.05 | | | | | | | | |
| $\chi^2_{causal}$ (SEM) | 24.63 (0.1) | 28.95 (0.2) | 36.35 (0.2) | 30.51 (0.2) | 39.01 (0.2) | 34.27 (0.2) | 45.06 (0.2) | 38.14 (0.2) | 51.28 (0.2) |
| Power | 0.28 (0.006) | 0.43 (0.007) | 0.68 (0.007) | 0.48 (0.007) | 0.75 (0.006) | 0.61 (0.007) | 0.87 (0.005) | 0.73 (0.006) | 0.94 (0.003) |
| $\chi^2_{null}$ (SEM) | 0.98 (0.02) | 0.95 (0.02) | 0.97 (0.02) | 0.94 (0.02) | 0.96 (0.02) | 0.97 (0.02) | 0.97 (0.02) | 0.99 (0.02) | 0.96 (0.02) |
| | K=0.25 | | | | | | | | |
| $\chi^2_{causal}$ (SEM) | 24.75 (0.09) | 18.88 (0.08) | 32.85 (0.1) | 17.9 (0.08) | 33.94 (0.1) | 15.44 (0.07) | 35.26 (0.1) | 12.5 (0.06) | 35.38 (0.1) |
| Power | 0.28 (0.004) | 0.11 (0.003) | 0.57 (0.005) | 0.089 (0.003) | 0.61 (0.005) | 0.045 (0.002) | 0.66 (0.004) | 0.018 (0.001) | 0.66 (0.004) |
| $\chi^2_{null}$ (SEM) | 1.03 (0.02) | 1.04 (0.02) | 1.04 (0.02) | 1.04 (0.02) | 1.04 (0.02) | 1.01 (0.02) | 1.02 (0.02) | 1.01 (0.02) | 1.02 (0.02) |

Table S11: **Results of PA formula in simulations with sibling history.** Number of individuals (N) is 100K and number of SNPs (M) is 1000 (1650 for $K = 0.25$ scenario); we assume perfect knowledge of $h^2$ and $K$ when implementing LT-FH; no environmental correlation between parents, offspring, or siblings; we consider 10 simulation replicates. The standard error of the mean $\chi^2$ for null and causal SNPs and the standard error for the power are reported in parentheses. For these simulation scenarios we modify the underlying prevalence of the disease ($K = 0.01$, 0.05, and 0.25) as well as the number of siblings each individual has ($n_s = 1, 2, 5, 10$); we modify C such that the GWAS power (and average GWAS causal $\chi^2$) is approximately the across all scenarios for comparative reasons. For LT-FH we assume that as in UK Biobank the number of affected siblings is not available; we assume all that is known is whether none or at least one sibling is affected. The PA formula can be used to approximate the mean posterior genetic liability assuming at least one affected sibling implies exactly one is affected. The effect of assuming exactly one sibling is affected, rather than at least one sibling, depends on the prevalence of the disease and the number of siblings an individual has.

| | $n_s = 1$ | | $n_s = 2$ | | $n_s = 5$ | | $n_s = 10$ | |
|---|---|---|---|---|---|---|---|---|
| | LT-FH | LT-PA | LT-FH | LT-PA | LT-FH | LT-PA | LT-FH | LT-PA |
| | K=0.01 | | | | | | | |
| $\chi^2_{causal}$ (SEM) | 38.13 (0.3) | 38.13 (0.3) | 42.08 (0.4) | 42 (0.4) | 52.6 (0.4) | 52.44 (0.4) | 66.14 (0.5) | 65.93 (0.5) |
| Power | 0.72 (0.01) | 0.71 (0.01) | 0.8 (0.009) | 0.8 (0.01) | 0.93 (0.006) | 0.93 (0.006) | 0.99 (0.002) | 0.99 (0.002) |
| $\chi^2_{null}$ (SEM) | 1.02 (0.02) | 1.02 (0.02) | 1.01 (0.02) | 1.01 (0.02) | 1.01 (0.02) | 1.01 (0.02) | 1 (0.02) | 1 (0.02) |
| | K=0.05 | | | | | | | |
| $\chi^2_{causal}$ (SEM) | 36.35 (0.2) | 36.35 (0.2) | 39.01 (0.2) | 38.82 (0.2) | 45.06 (0.2) | 44.85 (0.2) | 51.28 (0.2) | 50.82 (0.2) |
| Power | 0.68 (0.007) | 0.68 (0.007) | 0.75 (0.006) | 0.74 (0.006) | 0.87 (0.005) | 0.86 (0.005) | 0.94 (0.003) | 0.94 (0.003) |
| $\chi^2_{null}$ (SEM) | 0.97 (0.02) | 0.97 (0.02) | 0.96 (0.02) | 0.96 (0.02) | 0.97 (0.02) | 0.97 (0.02) | 0.96 (0.02) | 0.96 (0.02) |
| | K=0.25 | | | | | | | |
| $\chi^2_{causal}$ (SEM) | 32.85 (0.1) | 32.85 (0.1) | 33.94 (0.1) | 33.62 (0.1) | 35.26 (0.1) | 34.74 (0.1) | 35.38 (0.1) | 33.29 (0.1) |
| Power | 0.57 (0.005) | 0.57 (0.005) | 0.61 (0.005) | 0.6 (0.005) | 0.66 (0.004) | 0.64 (0.004) | 0.66 (0.004) | 0.59 (0.005) |
| $\chi^2_{null}$ (SEM) | 1.04 (0.02) | 1.04 (0.02) | 1.04 (0.02) | 1.04 (0.02) | 1.02 (0.02) | 1.03 (0.02) | 1.02 (0.02) | 1.02 (0.02) |

Table S12: **Results of GWAX+ method in simulations.** Number of individuals (N) and number of SNPs (M) is 100K; $h^2 = 0.5$; disease prevalence is 5%; we assume perfect knowledge of $h^2/K$ when implementing LT-FH; no environmental correlation between parents-offspring; we consider 10 simulation replicates. The standard error of the mean $\chi^2$ for null and causal SNPs and the standard error for the power are reported in parentheses. A GWAX alternative (GWAX+) defines controls to have value 0, proxy-cases value 0.5, and cases 1. A simple linear regression is then run, and the resulting $\chi^2$ is a 1df test statistic.

| | GWAS | GWAX-1df | GWAX+ | LT-FH |
|---|---|---|---|---|
| | K=0.01 | | | |
| $\chi^2_{causal}$ (SEM) | 24.1 (0.259) | 29.5 (0.288) | 33.8 (0.315) | 34 (0.316) |
| Power | 0.26 (0.01) | 0.46 (0.01) | 0.59 (0.01) | 0.58 (0.01) |
| $\chi^2_{null}$ (SEM) | 1 (0.001) | 1.001 (0.001) | 1 (0.001) | 1 (0.001) |
| | K=0.05 | | | |
| $\chi^2_{causal}$ (SEM) | 24.7 (0.146) | 27.3 (0.151) | 32.6 (0.167) | 33.2 (0.169) |
| Power | 0.28 (0.006) | 0.37 (0.007) | 0.56 (0.007) | 0.58 (0.007) |
| $\chi^2_{null}$ (SEM) | 1 (0.001) | 1.001 (0.001) | 1.001 (0.001) | 1.001 (0.001) |
| | K=0.25 | | | |
| $\chi^2_{causal}$ (SEM) | 24.7 (0.0928) | 20.1 (0.0824) | 28.6 (0.0993) | 31.1 (0.104) |
| Power | 0.28 (0.004) | 0.14 (0.003) | 0.42 (0.005) | 0.51 (0.005) |
| $\chi^2_{null}$ (SEM) | 0.997 (0.001) | 0.999 (0.001) | 0.998 (0.001) | 0.998 (0.001) |

Table S13: **Results of modeling variance heterogeneity.** Number of individuals (N) is 100K and number of SNPs (M) is 9680, 10000, and 10650 for the $K = 0.01, 0.05$, and 0.25 scenarios, respectively; we assume perfect knowledge of $h^2$ and $K$ when implementing LT-FH; no environmental correlation between parents and offspring; we consider 10 simulation replicates. For these simulation scenarios we modify the underlying prevalence of the disease (K = 0.01, 0.05 and 0.25) and modify the amount of variability in available family history. We explore when 5%, 10%, and 20% of the individuals have case-control information only and the rest have case-control and parent's history. The genetic value of an individual can be estimated using BLUP ($\hat{u}$), and the predictor error variance (variance of $\hat{u} - u$; PEV) can be computed (see Supplementary Note). In this simulation scenario it is straightforward to derive an analytical formula for $\hat{u}$ for individuals as well as the correponding PEV. We use linear regression to run LT-FH, LT-PA, LT-PA$_{binary}$, or use BLUP $\hat{u}$ without any weighting, or do a weighted linear regression with the inverse of the PEV as the weights.

| | | LT-FH | LT-PA | LT-PA$_{binary}$ | $\hat{u}$ | Weighted $\hat{u}$ |
|---|---|---|---|---|---|---|
| 5% case-control only; 95% case-control and parental history | | | | | | |
| K=0.01 | $\chi^2_{causal}$ (SEM) | 33.5 (0.313) | 33.5 (0.313) | 33.3 (0.313) | 33.3 (0.313) | 33.4 (0.313) |
| | Power | 0.571 (0.01) | 0.571 (0.01) | 0.566 (0.01) | 0.567 (0.01) | 0.568 (0.01) |
| | $\chi^2_{null}$ (SEM) | 1 (0.005) | 1 (0.005) | 1 (0.005) | 1 (0.005) | 1 (0.005) |
| K=0.05 | $\chi^2_{causal}$ (SEM) | 32.8 (0.168) | 32.8 (0.168) | 32.7 (0.168) | 32.7 (0.168) | 32.7 (0.168) |
| | Power | 0.564 (0.007) | 0.564 (0.007) | 0.558 (0.007) | 0.558 (0.007) | 0.559 (0.007) |
| | $\chi^2_{null}$ (SEM) | 0.992 (0.005) | 0.992 (0.005) | 0.992 (0.005) | 0.992 (0.005) | 0.993 (0.005) |
| K=0.25 | $\chi^2_{causal}$ (SEM) | 30.8 (0.104) | 30.8 (0.104) | 30.8 (0.104) | 30.8 (0.104) | 30.8 (0.104) |
| | Power | 0.502 (0.005) | 0.502 (0.005) | 0.501 (0.005) | 0.501 (0.005) | 0.502 (0.005) |
| | $\chi^2_{null}$ (SEM) | 1.009 (0.005) | 1.009 (0.005) | 1.008 (0.005) | 1.008 (0.005) | 1.01 (0.005) |
| 10% case-control only; 90% case-control and parental history | | | | | | |
| K=0.01 | $\chi^2_{causal}$ (SEM) | 32.9 (0.31) | 32.9 (0.31) | 32.8 (0.31) | 32.8 (0.31) | 32.8 (0.31) |
| | Power | 0.553 (0.01) | 0.551 (0.01) | 0.552 (0.01) | 0.552 (0.01) | 0.551 (0.01) |
| | $\chi^2_{null}$ (SEM) | 1.001 (0.005) | 1.001 (0.005) | 1.001 (0.005) | 1.001 (0.005) | 1.001 (0.005) |
| K =0.05 | $\chi^2_{causal}$ (SEM) | 32.4 (0.167) | 32.4 (0.167) | 32.3 (0.167) | 32.3 (0.167) | 32.3 (0.167) |
| | Power | 0.551 (0.007) | 0.55 (0.007) | 0.547 (0.007) | 0.547 (0.007) | 0.55 (0.007) |
| | $\chi^2_{null}$ (SEM) | 0.991 (0.005) | 0.991 (0.005) | 0.992 (0.005) | 0.992 (0.005) | 0.993 (0.005) |
| K=0.25 | $\chi^2_{causal}$ (SEM) | 30.5 (0.103) | 30.5 (0.103) | 30.4 (0.103) | 30.4 (0.103) | 30.5 (0.104) |
| | Power | 0.494 (0.005) | 0.494 (0.005) | 0.49 (0.005) | 0.49 (0.005) | 0.492 (0.005) |
| | $\chi^2_{null}$ (SEM) | 1.009 (0.005) | 1.009 (0.005) | 1.009 (0.005) | 1.009 (0.005) | 1.012 (0.005) |
| 20% case-control only; 80% case-control and parental history | | | | | | |
| K=0.01 | $\chi^2_{causal}$ (SEM) | 31.9 (0.306) | 31.9 (0.306) | 31.8 (0.305) | 31.8 (0.305) | 31.9 (0.305) |
| | Power | 0.519 (0.01) | 0.519 (0.01) | 0.513 (0.01) | 0.513 (0.01) | 0.514 (0.01) |
| | $\chi^2_{null}$ (SEM) | 1.003 (0.005) | 1.003 (0.005) | 1.003 (0.005) | 1.003 (0.005) | 1.004 (0.005) |
| K=0.05 | $\chi^2_{causal}$ (SEM) | 31.6 (0.165) | 31.6 (0.165) | 31.5 (0.164) | 31.5 (0.164) | 31.5 (0.165) |
| | Power | 0.52 (0.007) | 0.52 (0.007) | 0.519 (0.007) | 0.519 (0.007) | 0.522 (0.007) |
| | $\chi^2_{null}$ (SEM) | 0.991 (0.005) | 0.991 (0.005) | 0.991 (0.005) | 0.991 (0.005) | 0.994 (0.005) |
| K=0.25 | $\chi^2_{causal}$ (SEM) | 29.8 (0.102) | 29.8 (0.102) | 29.8 (0.102) | 29.8 (0.102) | 29.9 (0.103) |
| | Power | 0.469 (0.005) | 0.469 (0.005) | 0.468 (0.005) | 0.468 (0.005) | 0.472 (0.005) |
| | $\chi^2_{null}$ (SEM) | 1.007 (0.005) | 1.007 (0.005) | 1.007 (0.005) | 1.007 (0.005) | 1.013 (0.005) |

Table S14: **ICD9 and ICD10 codes for phenotype definition.** We report the ICD9 and ICD10 codes to define phenotypes for 12 UK Biobank diseases. We use instance 0 for self-reported and other non-accrruing phenotypes (i.e. collected during an in-person visit to an assessment center); for accruing data obtained from registries (e.g. death or cancer register) we use data from all instances. We included both information from hospitalization records and information from self-reported phenotypes.

| Trait | ICD9 codes | ICD10 codes | Other codes/definition | Notes |
|---|---|---|---|---|
| AD | Data-Fields 41203,41205; 3310 | Data-Fields 40001, 40002, 41202, 41204; F00, F000, F001, F002, F009, G30, G300, G301, G308, G309 | Non-cancer illness code, self-reported (Data-Field 20002; 1263) | Included self-report of dementia/alzheimers/cognitive impairment |
| Bowel Ca. | Data-Fields 41203,41205,40013; 153,1530:1539, 154,1540:1543, 1548 | Data-Fields 40001, 40002, 41202, 41204, 40006; C18,C180,C181,C182,C183, C184,C185,C186,C187, C188,C189,C19,C20, C21,C210,C211,C212,C218 | Cancer code, self-reported; (Data-Field 20001; 1020,1021,1022,1023) | Included colorectal cancer (any of colorectal, anal, colon, rectal) |
| Breast Ca. | Data-Fields 41203,41205,40013; 174,1740:1749 | Data-Fields 40001, 40002, 41202, 41204, 40006; C50, C500, C501, C502, C503, C504, C505, C506, C508, C509 | Cancer code, self-reported; (Data-Field 20001; 1002) | |
| CAD | | Data-Fields 40001, 40002, 41202, 41204; I20, I200, I201, I208, I209, I21, I210, I211, I212, I213, I214, I219, I21X, I22, I220, I221, I228, I229, I23, I230, I231, I232, I233, I234, I235, I236, I238, I24, I240, I241, I248, I249, I25, I250, I251, I252, I253, I254, I255, I256, I258, I259 | Vascular/heart problems diagnosed by doctor (Data-Field 6150; 1,2) and Non-cancer illness code, self-reported (Data-Field 20002; 1074,1075) | Defined heart disease as heart attack or angina and ischaemic heart diseases |
| COPD | Data-Fields 41203, 41205; 490, 4909, 491, 4910:4912, 4918, 4919, 492, 4929, 496, 4969 | Data-Fields 40001, 40002, 41202, 41204; J40, J41,J410, J411, J418, J42, J43, J430 ,J431, J432, J438, J439, J44, J440, J441, J448, J449 | Emphysema/chronic bronchitis diagnosed by doctor (Data-Field 6152; 6) & Non-cancer illness code, self-reported (Data-Field 20002; 1112,1113,1472) | Also include chronic obstructive pulmonary disease or chronic airways obstruction |
| HTN | | Data-Fields 40001, 40002, 41202, 41204; I10, I11, I110, I119, I12, I120, I129, I13, I130, I131, I132, I139, I15, I150, I151, I152, I158, I159 | Vascular/heart problems diagnosed by doctor (Data-Field 6150; 4) and Non-cancer illness code, self-reported (Data-Field 20002; 1065,1072) | |
| Lung Ca. | Data-Fields 41203,41205,40013; 162,1620, 1622:1625, 1628, 1629 | Data-Fields 40001, 40002, 41202, 41204, 40006; C33, C34, C340, C341, C342, C343, C348, C349 | Cancer code, self-reported; (Data-Field 20001; 1001, 1027, 1028,1080) | Lung cancer was cancer in any of trachea, bronchus, or lung |
| PD | Data-Fields 41203,41205; 332, 3320 | Data-Fields 40001, 40002, 41202, 41204; F023,G20 | Non-cancer illness code, self-reported (Data-Field 20002; 1262) | Excluded secondary parkinsonism |
| Prostate Ca. | Data-Fields 41203, 41205, 40013; 185, 1859 | Data-Fields 40001, 40002, 41202, 41204, 40006; C61 | Cancer code, self-reported (Data-Field 20001; 1044) | |
| Depression | | Data-Fields 40001, 40002, 41202, 41204; F32, F320, F321, F322, F323, F328, F329, F33, F330, F331, F332, F333, F334, F338, F339 | Non-cancer illness code, self-reported (Data-Field 20002; 1286) | Consider any depressive episodes or depressive disorders |
| Stroke | | Data-Fields 40001, 40002, 41202, 41204; I61, I610, I611, I612, I613, I614, I615, I616, I618, I619, I62, I620, I621, I629, I63, I630, I631, I632, I633, I634, I635, I636, I638, I639, I64 | Vascular/heart problems diagnosed by doctor (Data-Field 6150; 3) and Non-cancer illness code, self-reported (Data-Field 20002; 1081,1583) | |
| T2D | | | Diabetes diagnosed by doctor and age of diagnosis $> 30$ | |

Table S15: **Estimates of (liability-scale) narrow-sense heritability for 12 UK Biobank diseases.** We report estimates of (liability-scale) narrow-sense heritability ($h^2$) from the literature for the 12 diseases analyzed. These estimates are used as input to LT-FH.

| Trait | $h^2$ | Notes | References |
|---|---|---|---|
| AD | 0.79 | | Estimated using Swedish twins[3] |
| PD | 0.34 | Excluded secondary Parkinson's Disease | Estimated using Swedish twins[4] |
| Lung cancer | 0.18 | | Estimated using twins from the Nordic Twin Study of Cancer[5] |
| Bowel cancer | 0.40 | Heritability for colorectal cancer | Estimated using twins from the Nordic Twin Study of Cancer[6] |
| Stroke | 0.17 | Heritability for stroke hospitalization or stroke death | Estimated using Danish twins[7] |
| COPD | 0.6 | | Estimated using Danish (63%;95% CI: 46-77%) and Swedish twins (61%;95% CI: 48-72%)[8]; conclude estimate around 60%[8,9] |
| Prostate cancer | 0.57 | | Estimated using twins from the Nordic Twin Study of Cancer[5] |
| T2D | 0.72 | Assume family history of diabetes refers to type 2 diabetes | Estimated using twins from The Discordant Twin Consortium[10] |
| Breast cancer | 0.31 | | Estimated using twins from the Nordic Twin Study of Cancer[5] |
| Depression | 0.37 | Heritability for major depression | Meta-analysis of five twin studies[11] |
| CAD | 0.49 | Heritability for left main coronary artery disease | Estimated using German siblings[12] |
| HTN | 0.45 | Simple average of estimated heritability for SBP and DBP | Estimated using Dutch twin families[13] |

Table S16: **MAF thresholds applied to 12 UK Biobank diseases to avoid type I error in unbalanced case-control settings.** We report the MAF thresholds applied for GWAS, following recommendations of ref.[14] and ref.[15] (see Table S8 of ref.[15]) (in primary analyses, we used these MAF thresholds for GWAX and LT-FH as well); the MAF thresholds applied for GWAX in secondary analyses; and the MAF thresholds applied for LT-FH in secondary analyses. Because LT-FH phenotypes are not binary, MAF thresholds for LT-FH were computed based on relative kurtosis (where kurtosis is defined as $\kappa = E[(x-\mu)^4]/(E[(x-\mu)^2])^2$ and relative kurtosis is defined as $\kappa/3$); kurtosis was computed using the `R` package `moments`. MAF thresholds were chosen accordingly, based on the correspondence between disease prevalence and kurtosis reported in Table S17.

| | GWAS | | | GWAX | | | LT-FH | |
|---|---|---|---|---|---|---|---|---|
| Trait | K | $\hat{\kappa}/3$ | MAF | K | $\hat{\kappa}/3$ | MAF | $\hat{\kappa}/3$ | MAF |
| AD | 0.001 | 286.000 | 0.100 | 0.141 | 1.750 | 0.001 | 4.630 | 0.001 |
| PD | 0.003 | 106.000 | 0.100 | 0.052 | 5.780 | 0.001 | 15.300 | 0.010 |
| Lung cancer | 0.006 | 56.100 | 0.100 | 0.154 | 1.560 | 0.001 | 5.650 | 0.001 |
| Bowel cancer | 0.013 | 25.200 | 0.010 | 0.144 | 1.710 | 0.001 | 5.590 | 0.001 |
| Stroke | 0.023 | 13.600 | 0.010 | 0.318 | 0.536 | 0.001 | 3.070 | 0.001 |
| COPD | 0.035 | 8.890 | 0.010 | 0.209 | 1.010 | 0.001 | 3.670 | 0.001 |
| Prostate cancer | 0.037 | 8.240 | 0.010 | 0.107 | 2.500 | 0.001 | 7.010 | 0.010 |
| T2D | 0.042 | 7.360 | 0.010 | 0.262 | 0.722 | 0.001 | 2.960 | 0.001 |
| Breast cancer | 0.061 | 4.800 | 0.001 | 0.146 | 1.670 | 0.001 | 4.230 | 0.001 |
| Depression | 0.073 | 3.900 | 0.001 | 0.215 | 0.978 | 0.001 | 2.640 | 0.001 |
| CAD | 0.083 | 3.360 | 0.001 | 0.522 | 0.336 | 0.001 | 1.560 | 0.001 |
| HTN | 0.318 | 0.537 | 0.001 | 0.657 | 0.479 | 0.001 | 0.721 | 0.001 |

Table S17: **Correspondence between disease prevalence and kurtosis determines MAF thresholds for LT-FH in secondary analyses.** We report the correspondence between disease prevalence and kurtosis: prevalence $< 1\%$ or relative kurtosis $> 32.670$ implies MAF $\geq 0.1$; prevalence $1-5\%$ or relative kurtosis of 6.018 to 32.670 implies MAF $\geq 0.01$; and prevalence$> 5\%$ or relative kurtosis of $< 6.018$ implies MAF $\geq 0.001$.

| Prev. | $\kappa/3$ |
|---|---|
| 0.001 | 332.667 |
| 0.005 | 66.002 |
| 0.010 | 32.670 |
| 0.050 | 6.018 |
| 0.100 | 2.704 |
| 0.250 | 0.778 |
| 0.500 | 0.333 |

Table S18: **Completeness of parental history and sibling history information for 12 UK Biobank diseases.** For the 381,493 unrelated European individuals, we report the percentage that report complete parental history (presence or absence of disease in both mother and father) among all individuals and those with known case-control status and the percentage that report complete sibling history (either 0 siblings, or $>0$ siblings and presence or absence of disease in the set of all siblings). For sex-specific traits (breast and prostate cancer) we report the percentage reporting disease information in the parent with relevant sex (e.g. for breast cancer the proportion reporting maternal history of breast cancer) and when reporting the percentage with complete sibling history we restrict to the number of siblings with the relevant sex (e.g. for breast cancer the proportion reporting either 0 sisters, or $>0$ sisters and presence or absence of disease in the set of all sisters).

| | Complete Parental History | | |
|---|---|---|---|
| Traits | All | Known case-control | Complete Sibling History |
| AD | 0.879 | 0.879 | 0.924 |
| PD | 0.865 | 0.865 | 0.924 |
| LungCancer | 0.865 | 0.865 | 0.924 |
| BowelCancer | 0.865 | 0.865 | 0.924 |
| Stroke | 0.879 | 0.879 | 0.924 |
| COPD | 0.879 | 0.879 | 0.924 |
| ProstateCancer | 0.893 | 0.882 | 0.938 |
| T2D | 0.879 | 0.880 | 0.924 |
| BreastCancer | 0.934 | 0.950 | 0.944 |
| Depression | 0.865 | 0.865 | 0.924 |
| CAD | 0.879 | 0.879 | 0.924 |
| HTN | 0.879 | 0.879 | 0.924 |
| Average | 0.880 | 0.881 | 0.927 |

Table S19: **Definition of GWAS and GWAX phenotypes for UK Biobank diseases.** We report GWAS, GWAX, and GWAX-2df phenotypes for all combinations of case-control status and parental, or parental and sibling, history of disease. Sibling history can be 1 ($\geq 1$ affected), 0 (none affected and $\geq 1$ sibling) and $NA$. When an individual reports having no siblings, we define GWAX phenotypes based on parental history only, otherwise we define GWAX phenotypes based on the parent & sibling history table. Sex-specific diseases are breast cancer and prostate cancer.

<table>
<tr><th colspan="5">Parent History</th></tr>
<tr><td colspan="5">Non-sex specific diseases</td></tr>
<tr><td>Child</td><td>Family history</td><td>GWAS</td><td>GWAX</td><td>GWAX-2df</td></tr>
<tr><td>1</td><td>Anything</td><td>1</td><td>1</td><td>2</td></tr>
<tr><td>0</td><td>p1=1|p2=1</td><td>0</td><td>1</td><td>1</td></tr>
<tr><td>0</td><td>p1,p2 $\in \{0, NA\}$ & p1!=p2!=0</td><td>0</td><td>NA</td><td>NA</td></tr>
<tr><td>0</td><td>p1=p2=0</td><td>0</td><td>0</td><td>0</td></tr>
<tr><td>NA</td><td>p1=1|p2=1</td><td>NA</td><td>1</td><td>NA</td></tr>
<tr><td>NA</td><td>p1 != 1 & p2!=1</td><td>NA</td><td>NA</td><td>NA</td></tr>
<tr><td colspan="5">Sex-specific diseases</td></tr>
<tr><td>Child</td><td>Family history</td><td>GWAS</td><td>GWAX</td><td>GWAX-2df</td></tr>
<tr><td>1</td><td>Anything</td><td>1</td><td>1</td><td>2</td></tr>
<tr><td>0</td><td>p1=1</td><td>0</td><td>1</td><td>1</td></tr>
<tr><td>0</td><td>p1=NA</td><td>0</td><td>NA</td><td>NA</td></tr>
<tr><td>0</td><td>p1=0</td><td>0</td><td>0</td><td>0</td></tr>
<tr><td>NA (relevant sex)</td><td>p1=1</td><td>NA</td><td>1</td><td>NA</td></tr>
<tr><td>NA (non-relevant sex)</td><td>p1=1</td><td>NA</td><td>1</td><td>1</td></tr>
<tr><td>NA (relevant sex)</td><td>p1=0</td><td>NA</td><td>NA</td><td>NA</td></tr>
<tr><td>NA (non-relevant sex)</td><td>p1=0</td><td>NA</td><td>0</td><td>0</td></tr>
<tr><td>NA</td><td>p1=NA</td><td>NA</td><td>NA</td><td>NA</td></tr>
<tr><th colspan="5">Parent & Sibling History</th></tr>
<tr><td colspan="5">Non-sex specific diseases</td></tr>
<tr><td>Child</td><td>Family history</td><td>GWAS</td><td>GWAX</td><td>GWAX-2df</td></tr>
<tr><td>1</td><td>Anything</td><td>1</td><td>1</td><td>2</td></tr>
<tr><td>0</td><td>p1=1|p2=1|s=1</td><td>0</td><td>1</td><td>1</td></tr>
<tr><td>0</td><td>p1,p2,s $\in \{0, NA\}$ & p1!=p2!=s!=0</td><td>0</td><td>NA</td><td>NA</td></tr>
<tr><td>0</td><td>s=p1=p2=0 | ∅&p1=p2=0</td><td>0</td><td>0</td><td>0</td></tr>
<tr><td>NA</td><td>p1=1|p2=1|s=1</td><td>NA</td><td>1</td><td>NA</td></tr>
<tr><td>NA</td><td>p1 != 1 & p2!=1 & s!=1</td><td>NA</td><td>NA</td><td>NA</td></tr>
<tr><td colspan="5">Sex-specific diseases</td></tr>
<tr><td>Child</td><td>Family history</td><td>GWAS</td><td>GWAX</td><td>GWAX-2df</td></tr>
<tr><td>1</td><td>Anything</td><td>1</td><td>1</td><td>2</td></tr>
<tr><td>0</td><td>p1=1|s=1</td><td>0</td><td>1</td><td>1</td></tr>
<tr><td>0</td><td>p1,s $\in \{0, NA\}$ & p1!=s!=0</td><td>0</td><td>NA</td><td>NA</td></tr>
<tr><td>0</td><td>s=p1=0 | ∅&p1=0</td><td>0</td><td>0</td><td>0</td></tr>
<tr><td>NA (relevant sex)</td><td>p1=1|s=1</td><td>NA</td><td>1</td><td>NA</td></tr>
<tr><td>NA (non-relevant sex)</td><td>p1=1|s=1</td><td>NA</td><td>1</td><td>1</td></tr>
<tr><td>NA (relevant sex)</td><td>p1=s=0| ∅&p1=0</td><td>NA</td><td>NA</td><td>NA</td></tr>
<tr><td>NA (non-relevant sex)</td><td>p1=s=0| ∅&p1=0</td><td>NA</td><td>0</td><td>0</td></tr>
<tr><td>NA</td><td>p1,s $\in \{0, NA\}$ & p1 != s != 0</td><td>NA</td><td>NA</td><td>NA</td></tr>
</table>

Table S20: **Computational cost of computing LT-FH phenotypes and computing association statistics.** We report the number of hours required to compute LT-FH phenotypes (constructed using the LT-FH software v2; we note that this includes computation of both LT-FH$_{no-sib}$ and LT-FH), the number of hours required to compute association statistics using linear regression, and the number of hours required to compute association statistics using BOLT-LMM.

| Trait | Constructing LT-FH | Linear regression (BOLT-LMM software) | BOLT-LMM (BOLT-LMM software) |
|---|---:|---:|---:|
| AD | 0.7 | 26.4 | 55.2 |
| PD | 0.7 | 25.5 | 29.8 |
| LungCancer | 0.7 | 21.7 | 35.7 |
| BowelCancer | 0.7 | 21.3 | 33.5 |
| Stroke | 0.9 | 25.1 | 32.3 |
| COPD | 0.7 | 24.7 | 36.6 |
| ProstateCancer | 0.5 | 22.1 | 34.1 |
| T2D | 0.9 | 23.8 | 40.3 |
| BreastCancer | 0.6 | 22.1 | 35.7 |
| Depression | 1.0 | 19.3 | 31.5 |
| CAD | 0.7 | 22.0 | 49.6 |
| HTN | 0.7 | 22.7 | 63.0 |
| Mean | 0.74 | 23.06 | 39.79 |
| Median | 0.72 | 22.41 | 35.67 |

Table S21: **Results of GWAS, GWAX and LT-FH in analyses of 12 UK Biobank diseases (restricted to unrelated individuals) using linear regression.** We report **(a)** attenuation ratios and difference in ratios between LT-FH and GWAS (standard errors in parentheses); **(b)** number of independent loci; and average $\chi^2$ for **(c)** genome wide-significant SNPs ($p \leq 5 * 10^{-8}$ for at least one method) and **(d)** all SNPs. In (c), we compute weighted averages in which the weight is determined by the number of genome-wide significant SNPs (shown for each disease in parentheses). In (c) and (d), we restrict to SNPs above the MAF threshold for each disease (reported in Table S16).

| **(a)** | Attenuation Ratio | | | |
|---|---|---|---|---|
| Traits | GWAS | GWAX | LT-FH | LT-FH−GWAS |
| AD | 0.420 (0.884) | 0.408 (0.188) | 0.381 (0.190) | -0.038 (0.812) |
| PD | 0.810 (0.454) | 0.254 (0.202) | 0.316 (0.172) | -0.493 (0.408) |
| Lung cancer | 0.000 (0.298) | 0.104 (0.060) | 0.069 (0.068) | 0.069 (0.280) |
| Bowel cancer | 0.160 (0.146) | 0.149 (0.081) | 0.106 (0.074) | -0.054 (0.127) |
| Stroke | 0.179 (0.180) | 0.149 (0.077) | 0.139 (0.069) | -0.040 (0.173) |
| COPD | 0.077 (0.059) | 0.158 (0.036) | 0.106 (0.034) | 0.029 (0.048) |
| Prostate cancer | 0.140 (0.093) | 0.085 (0.086) | 0.116 (0.075) | -0.024 (0.057) |
| T2D | 0.123 (0.043) | 0.164 (0.036) | 0.136 (0.038) | 0.012 (0.021) |
| Breast cancer | 0.117 (0.081) | 0.191 (0.064) | 0.176 (0.053) | 0.058 (0.059) |
| Depression | 0.034 (0.054) | 0.115 (0.035) | 0.108 (0.037) | 0.074 (0.038) |
| CAD | 0.047 (0.035) | 0.084 (0.033) | 0.086 (0.021) | 0.039 (0.027) |
| HTN | 0.095 (0.017) | 0.090 (0.021) | 0.075 (0.016) | -0.020 (0.007) |
| Flat mean | 0.183 (0.090) | 0.162 (0.028) | 0.151 (0.025) | -0.032 (0.082) |
| Inv-var. weighted mean | 0.089 (0.013) | 0.105 (0.014) | 0.090 (0.011) | 0.001 (0.007) |

| **(b)** | Number of independent loci | | |
|---|---|---|---|
| Traits | GWAS | GWAX | LT-FH |
| AD | 1 | 8 | 11 |
| PD | 1 | 4 | 4 |
| Lung cancer | 0 | 5 | 5 |
| Bowel cancer | 4 | 9 | 17 |
| Stroke | 0 | 3 | 7 |
| COPD | 5 | 14 | 12 |
| Prostate cancer | 28 | 29 | 38 |
| T2D | 57 | 82 | 120 |
| Breast cancer | 28 | 40 | 49 |
| Depression | 1 | 4 | 6 |
| CAD | 35 | 46 | 92 |
| HTN | 263 | 114 | 329 |
| Total | 423 | 358 | 690 |

| **(c)** | Mean $\chi^2$ over all genome-significant SNPs* | | |
|---|---|---|---|
| Traits | GWAS | GWAX | LT-FH |
| AD (551) | 23.07 | 207.23 | 224.71 |
| PD (2518) | 9.28 | 42.61 | 46.06 |
| Lung cancer (627) | 9.31 | 56.92 | 60.65 |
| Bowel cancer (526) | 21.18 | 36.77 | 45.15 |
| Stroke (372) | 11.21 | 35.72 | 41.48 |
| COPD (1034) | 19.43 | 45.07 | 46.72 |
| Prostate cancer (2862) | 40.27 | 40.39 | 56.31 |
| T2D (7918) | 31.21 | 44.90 | 58.91 |
| Breast cancer (3844) | 32.38 | 45.00 | 54.39 |
| Depression (400) | 23.66 | 36.67 | 37.02 |

| **(d)** | Mean $\chi^2$ over all tested SNPs† | | |
|---|---|---|---|
| Traits | GWAS | GWAX | LT-FH |
| AD | 1.01 | 1.11 | 1.11 |
| PD | 1.02 | 1.07 | 1.08 |
| Lung cancer | 1.03 | 1.21 | 1.19 |
| Bowel cancer | 1.04 | 1.09 | 1.10 |
| Stroke | 1.04 | 1.10 | 1.10 |
| COPD | 1.14 | 1.25 | 1.25 |
| Prostate cancer | 1.09 | 1.10 | 1.14 |
| T2D | 1.29 | 1.40 | 1.50 |
| Breast cancer | 1.06 | 1.10 | 1.12 |
| Depression | 1.09 | 1.15 | 1.15 |

| | | | |
|---|---|---|---|
| CAD (5620) | 29.26 | 36.12 | 56.93 |
| HTN (28353) | 37.72 | 25.20 | 46.25 |
| Average | 33.09 | 35.01 | 52.14 |

| | | | |
|---|---|---|---|
| CAD | 1.17 | 1.19 | 1.29 |
| HTN | 1.56 | 1.36 | 1.63 |
| Average | 1.13 | 1.18 | 1.22 |

Table S22: **Relative effective sample sizes of LT-FH vs. GWAS and LT-FH vs. GWAX in analyses of 12 UK Biobank diseases (restricted to unrelated individuals) using linear regression.** We estimate the relative effective sample size achieved by LT-FH by measuring the boosts in $\chi^2$ linear regression association statistics of LT-FH versus GWAS (or GWAX) on unrelated European samples (as outlined in Loh et al. 2018 *Nature Genetics*[15]). In more detail, the relative effective sample size of LT-FH versus GWAS (and LT-FH versus GWAX) is computed as the median ratio of LT-FH $\chi^2$ statistics to GWAS (GWAX) $\chi^2$ statistics across genotyped SNPs with $\chi^2 \geq 30$ in GWAS applied via BOLT-LMM to all related Europeans. Ratio values are shown only when the number of SNPs used to calculate the ratio is $\geq 10$, otherwise NA is displayed; the number of SNPs used to calculate the ratio is shown (next to the trait in parentheses). The average effective sample size relative to GWAS, weighted by the number of significant loci found by GWAS, is 1.31 (this removes PD, Lung Cancer, Stroke, and Depression). The relative improvement of LT-FH vs GWAS in terms of number of loci found via linear regression is 1.63 across all traits; the relative improvement restricting to all traits for which $N_{eff}$ is non-NA is 1.59. The average effective sample size relative to GWAX, weighted by the number of significant loci found by GWAX, is 1.65. Finally, if for each trait we select the best method between GWAS and GWAX as the method which discovers the most loci, the average effective sample size for LT-FH relative to this trait-specific "best" method (weighted by the number of loci found by the trait-specific "best" method) is 1.27.

| | Relative effective sample size of LT-FH vs. | |
|---|---|---|
| Trait | GWAS | GWAX |
| AD (11) | 10.020 | 1.092 |
| PD (1) | NA | NA |
| LungCancer (0) | NA | NA |
| BowelCancer (11) | 1.568 | 1.482 |
| Stroke (0) | NA | NA |
| COPD (39) | 2.513 | 1.109 |
| ProstateCancer (129) | 1.259 | 1.512 |
| T2D (269) | 1.705 | 1.432 |
| BreastCancer (118) | 1.417 | 1.362 |
| Depression (5) | NA | NA |
| CAD (159) | 1.745 | 1.722 |
| HTN (1458) | 1.098 | 2.019 |

Table S23: **Correlations between $-log_{10}(p)$ values for GWAS, GWAX and LT-FH for 12 UK Biobank diseases.** We report the correlation between $-log_{10}$ association $p$-values for each pair of GWAS, GWAX and LT-FH. We restrict to SNPs above the MAF threshold for each disease (reported in Table S16). $p$-values reported as 0 are replaced with $2.23 * 10^{-308}$, the smallest positive number meeting requirements of the IEEE Standard for Floating-Point Arithmetic.

| Trait | Correlation between $-log_{10}(p)$ | | |
|---|---|---|---|
| | GWAS/ GWAX | GWAS/ LT-FH | GWAX/ LT-FH |
| AD | 0.30 | 0.33 | 0.98 |
| PD | 0.13 | 0.34 | 0.90 |
| Lung cancer | 0.08 | 0.26 | 0.86 |
| Bowel cancer | 0.14 | 0.44 | 0.82 |
| Stroke | 0.08 | 0.39 | 0.74 |
| COPD | 0.25 | 0.63 | 0.79 |
| Prostate cancer | 0.35 | 0.69 | 0.84 |
| T2D | 0.56 | 0.80 | 0.89 |
| Breast cancer | 0.38 | 0.70 | 0.85 |
| Depression | 0.39 | 0.71 | 0.80 |
| CAD | 0.31 | 0.69 | 0.72 |
| HTN | 0.61 | 0.89 | 0.76 |
| Average | 0.30 | 0.57 | 0.83 |

Table S24: **Sample size times observed-scale SNP-heritability for GWAS, GWAX and LT-FH phenotypes for 12 UK Biobank diseases.** We report estimates of $N * h^2_{g,o}$, which provides a measure of total genetic signal[16]. Observed-scale SNP-heritability is estimated using BOLT-REML[17]. Estimates of $N * h^2_{g,o}$ are 59% higher for LT-FH vs. GWAS, and higher for all 12 diseases. Standard errors are reported in parentheses.

| | GWAS | | | GWAX | | | LT-FH | | |
|---|---|---|---|---|---|---|---|---|---|
| Trait | N | $\hat{h}^2_{g,o}$ (s.e.) | $N * \hat{h}^2_{g,o}$ | N | $\hat{h}^2_{g,o}$ (s.e.) | $N * \hat{h}^2_{g,o}$ | N | $\hat{h}^2_{g,o}$ (s.e.) | $N * \hat{h}^2_{o}$ |
| AD | 381493 | 0.001 (0.001) | 381.49 | 324512 | 0.035 (0.002) | 11357.92 | 381493 | 0.031 (0.002) | 11826.28 |
| PD | 381493 | 0.004 (0.001) | 1525.97 | 318792 | 0.012 (0.002) | 3825.50 | 381493 | 0.013 (0.002) | 4959.41 |
| Lung cancer | 381493 | 0.005 (0.001) | 1907.47 | 323838 | 0.039 (0.002) | 12629.68 | 381493 | 0.030 (0.002) | 11444.79 |
| Bowel cancer | 381493 | 0.012 (0.002) | 4577.92 | 322887 | 0.028 (0.002) | 9040.84 | 381493 | 0.027 (0.002) | 10300.31 |
| Stroke | 381493 | 0.010 (0.002) | 3814.93 | 332468 | 0.027 (0.002) | 8976.64 | 381493 | 0.026 (0.002) | 9918.82 |
| COPD | 381493 | 0.031 (0.002) | 11826.28 | 329495 | 0.064 (0.002) | 21087.68 | 381493 | 0.057 (0.002) | 21745.10 |
| Prostate cancer | 175450 | 0.056 (0.004) | 9825.20 | 331458 | 0.032 (0.002) | 10606.66 | 368940 | 0.040 (0.002) | 14757.60 |
| T2D | 380180 | 0.072 (0.002) | 27372.96 | 330267 | 0.109 (0.002) | 35999.10 | 381390 | 0.119 (0.002) | 45385.41 |
| Breast cancer | 206043 | 0.050 (0.003) | 10302.15 | 348170 | 0.042 (0.002) | 14623.14 | 371064 | 0.048 (0.002) | 17811.07 |
| Depression | 381493 | 0.033 (0.002) | 12589.27 | 328186 | 0.057 (0.002) | 18706.60 | 381493 | 0.050 (0.002) | 19074.65 |
| CAD | 381493 | 0.062 (0.002) | 23652.57 | 341810 | 0.075 (0.002) | 25635.75 | 381493 | 0.101 (0.002) | 38530.79 |
| HTN | 381493 | 0.180 (0.002) | 68668.74 | 354208 | 0.121 (0.002) | 42859.17 | 381493 | 0.197 (0.002) | 75154.12 |
| Average | 349592.5 | 0.043 | 14703.75 | 332174.2 | 0.053 | 17945.72 | 379569.2 | 0.062 | 23409.03 |

Table S25: **Results of replication analysis of 4 diseases in non-UK Biobank data sets.** We conducted a replication analysis of loci identified by GWAS and/or LT-FH in independent non-UK Biobank data sets for 4 diseases (coronary artery disease, type 2 diabetes, breast cancer, and prostate cancer) with publicly available summary statistics[18–21]. For type 2 diabetes, the summary statistics used were computed using only stage 1 data consisting of 12,171 cases and 56,862 controls[19], for prostate cancer the summary statistics used were computed using the OncoArray European sample consisting of 27,904 controls and 44,825 cases[21]. The replication summary statistics are from studies consisting of predominantly non-UK Europeans and were always computed using GWAS (not LT-FH). The replication slope (the slope of a regression of standardized effect sizes of lead SNPs in case-control replication data vs. GWAS or LT-FH UK Biobank discovery data) is shown on a trait-specific basis and over all traits.

| | GWAS | | | | LT-FH | | | |
|---|---|---|---|---|---|---|---|---|
| | GWAS Only | | All GWAS | | LT-FH Only | | All LT-FH | |
| Trait | n | Slope(se) | n | Slope(se) | n | Slope(se) | n | Slope(se) |
| BreastCancer | 1 | NA (NA) | 24 | 0.85 (0.03) | 17 | 0.76 (0.04) | 40 | 0.84 (0.02) |
| CAD | 2 | 417.91 (NaN) | 24 | 0.72 (0.07) | 41 | 0.92 (0.06) | 63 | 0.84 (0.05) |
| ProstateCancer | 3 | 0.61 (0.05) | 26 | 0.85 (0.03) | 10 | 0.64 (0.07) | 33 | 0.82 (0.03) |
| T2D | 1 | NA (NA) | 50 | 0.74 (0.03) | 58 | 0.61 (0.05) | 107 | 0.71 (0.03) |
| All | 7 | 0.67 (0.06) | 124 | 0.81 (0.02) | 126 | 0.69 (0.03) | 243 | 0.79 (0.01) |

Table S26: **Power of GWAS, GWAX, GWAX-2df and LT-FH in analyses of 12 UK Biobank diseases (restricted to unrelated individuals and genotyped SNPs).** We report the number of independent loci (restricted to genotyped SNPs) for GWAS, GWAX, GWAX-2df, and LT-FH.

| Trait | GWAS | GWAX | GWAX-2df | LT-FH |
|---|---|---|---|---|
| AD | 1 | 7 | 8 | 9 |
| PD | 1 | 3 | 3 | 4 |
| Lung cancer | 0 | 5 | 3 | 3 |
| Bowel cancer | 3 | 8 | 10 | 12 |
| Stroke | 0 | 2 | 3 | 3 |
| COPD | 2 | 10 | 9 | 10 |
| Prostate cancer | 26 | 25 | 33 | 34 |
| T2D | 46 | 65 | 77 | 96 |
| Breast cancer | 22 | 32 | 36 | 39 |
| Depression | 1 | 1 | 1 | 3 |
| CAD | 27 | 39 | 54 | 75 |
| HTN | 195 | 74 | 175 | 244 |
| Total | 324 | 271 | 412 | 532 |

Table S27: **Results of GWAX$_{no-sib}$ and LT-FH$_{no-sib}$ in analyses of 12 UK Biobank diseases (restricted to unrelated individuals) using linear regression.** We report **(a)** attenuation ratios and **(b)** number of independent loci for GWAS, GWAX$_{no-sib}$, GWAX, LT-FH$_{no-sib}$ and LT-FH across 12 diseases. Standard errors are reported in parentheses.

| Trait | GWAS | GWAX$_{no-sib}$ | GWAX | LT-FH $_{no-sib}$ | LT-FH |
|---|---|---|---|---|---|
| **(a)** | | | Attenuation Ratio | | |
| AD | 0.420 (0.884) | 0.424 (0.184) | 0.408 (0.188) | 0.395 (0.179) | 0.381 (0.190) |
| PD | 0.810 (0.454) | 0.333 (0.222) | 0.254 (0.202) | 0.334 (0.189) | 0.316 (0.172) |
| Lung cancer | 0.000 (0.298) | 0.084 (0.071) | 0.104 (0.060) | 0.057 (0.075) | 0.069 (0.068) |
| Bowel cancer | 0.160 (0.146) | 0.187 (0.082) | 0.149 (0.081) | 0.185 (0.075) | 0.106 (0.074) |
| Stroke | 0.179 (0.180) | 0.119 (0.087) | 0.149 (0.077) | 0.116 (0.072) | 0.139 (0.069) |
| COPD | 0.077 (0.059) | 0.150 (0.039) | 0.158 (0.036) | 0.112 (0.035) | 0.106 (0.034) |
| Prostate cancer | 0.140 (0.093) | 0.134 (0.088) | 0.085 (0.086) | 0.136 (0.078) | 0.116 (0.075) |
| T2D | 0.123 (0.043) | 0.184 (0.039) | 0.164 (0.036) | 0.155 (0.040) | 0.136 (0.038) |
| Breast cancer | 0.117 (0.081) | 0.230 (0.063) | 0.191 (0.064) | 0.190 (0.056) | 0.176 (0.053) |
| Depression | 0.034 (0.054) | 0.143 (0.041) | 0.115 (0.035) | 0.091 (0.041) | 0.108 (0.037) |
| CAD | 0.047 (0.035) | 0.072 (0.034) | 0.084 (0.033) | 0.083 (0.022) | 0.086 (0.021) |
| HTN | 0.095 (0.017) | 0.096 (0.020) | 0.090 (0.021) | 0.082 (0.016) | 0.075 (0.016) |
| Flat mean | 0.183 (0.090) | 0.180 (0.029) | 0.162 (0.028) | 0.161 (0.026) | 0.151 (0.025) |
| Inv-var. weighted mean | 0.089 (0.013) | 0.112 (0.014) | 0.105 (0.014) | 0.097 (0.011) | 0.090 (0.011) |
| **(b)** | | | Number of independent loci | | |
| AD | 1 | 9 | 8 | 13 | 11 |
| PD | 1 | 4 | 4 | 5 | 4 |
| Lung cancer | 0 | 5 | 5 | 5 | 5 |
| Bowel cancer | 4 | 9 | 9 | 14 | 17 |
| Stroke | 0 | 3 | 3 | 7 | 7 |
| COPD | 5 | 11 | 14 | 11 | 12 |
| Prostate cancer | 28 | 26 | 29 | 41 | 38 |
| T2D | 57 | 78 | 82 | 112 | 120 |
| Breast cancer | 28 | 32 | 40 | 39 | 49 |
| Depression | 1 | 5 | 4 | 4 | 6 |
| CAD | 35 | 44 | 46 | 83 | 92 |
| HTN | 263 | 125 | 114 | 317 | 329 |
| Total | 423 | 351 | 358 | 651 | 690 |

Table S28: **Correlation of self-reported family history between sibling pairs for 12 UK Biobank diseases.** We report the correlation of self-reported parental history and self-reported sibling history between sibling pairs. The correlation of self-reported sibling history is restricted to concordant sibling pairs (e.g. both cases or both controls). For sex-specific diseases (breast cancer and prostate cancer), we restrict to concordant sibling pairs of the relevant sex, sibling pairs of the non-relevant sex, and sibling pairs of discordant sex for which the sibling of the relevant sex is a control. The correlation of self-reported number of siblings between sibling pairs is 0.956. The sibling pair correlation of self-reported family history incorporates the inaccuracy of both siblings; the correlation between true and self-reported family history is equal to the square root of the correlation of self-reported family history, if errors are uncorrelated between siblings. The square root of the average correlation is 0.827 for number of affected parents and 0.764 for sibling history.

| Traits | $r_{sib}$(# affected parents) | $r_{sib}$(sibling history) |
|---|---|---|
| AD | 0.692 | 0.591 |
| PD | 0.799 | 0.759 |
| Lung cancer | 0.792 | 0.676 |
| Bowel cancer | 0.722 | 0.633 |
| Stroke | 0.656 | 0.517 |
| COPD | 0.584 | 0.275 |
| Prostate cancer | 0.722 | 0.660 |
| T2D | 0.820 | 0.685 |
| Breast cancer | 0.891 | 0.805 |
| Depression | 0.443 | 0.322 |
| CAD | 0.608 | 0.589 |
| HTN | 0.485 | 0.488 |
| Average | 0.685 | 0.583 |

Table S29: **Impact of modifying the LT-FH method to downweight family history information based on its accuracy for 12 UK Biobank diseases.** We compared LT-FH$_{no-sib}$ and LT-FH$_{no-sib}^{FHweighted}$. We report phenotypic correlations, attenuation ratios, and values of the number of independent loci across 12 diseases. Standard errors are reported in parentheses.

| | | Attenuation Ratio | | Independent loci | |
|---|---|---|---|---|---|
| Traits | $\rho$ | LT-FH$_{no-sib}$ | LT-FH$_{no-sib}^{FHweighted}$ | LT-FH$_{no-sib}$ | LT-FH$_{no-sib}^{FHweighted}$ |
| AD | 0.9983 | 0.395 (0.179) | 0.397 (0.181) | 13 | 12 |
| PD | 0.9985 | 0.334 (0.189) | 0.341 (0.189) | 5 | 5 |
| Lung cancer | 0.9986 | 0.057 (0.075) | 0.055 (0.076) | 5 | 4 |
| Bowel cancer | 0.9966 | 0.185 (0.075) | 0.183 (0.076) | 14 | 14 |
| Stroke | 0.9945 | 0.116 (0.072) | 0.117 (0.073) | 7 | 8 |
| COPD | 0.9918 | 0.112 (0.035) | 0.106 (0.036) | 11 | 11 |
| Prostate cancer | 0.9970 | 0.136 (0.078) | 0.138 (0.078) | 41 | 41 |
| T2D | 0.9989 | 0.155 (0.040) | 0.153 (0.040) | 112 | 111 |
| Breast cancer | 0.9997 | 0.190 (0.056) | 0.188 (0.056) | 39 | 39 |
| Depression | 0.9894 | 0.091 (0.041) | 0.073 (0.043) | 4 | 3 |
| CAD | 0.9928 | 0.083 (0.022) | 0.080 (0.023) | 83 | 83 |
| HTN | 0.9907 | 0.082 (0.016) | 0.085 (0.016) | 317 | 320 |
| Flat mean/Total | 0.996 | 0.161 (0.026) | 0.160 (0.026) | 651 | 651 |
| Inv-var. weighted mean | | 0.097 (0.011) | 0.096 (0.011) | | |

Table S30: **Liability threshold model parameters for incorporating age into LT-FH for 12 UK Biobank diseases.** We incorporate age into the liability model through modeling $\phi = m + c_{age}(age - a\bar{g}e) + \epsilon$[22]. $m$ is an affine parameter determining prevalence at the mean age, thus a more negative $m$ value represents a less prevalence trait. A positive $c_{age}$ value implies increasing prevalence with age. For breast cancer these values were estimated in females only (for prostate cancer we restricted to males). In short, we found the disease prevalence for each 5 year age bin (<45,45-50,. . .,85-90,90+) by combining genotyped individuals' and their parents' case-status; we clumped bins until every bin contained at least 100 cases (Table S31). We then used this age (mean age of individuals in each 5-year age bin) and prevalence data to compute the effect of age on the liability scale using LTSOFT[22] (see URLs). We assigned relevant parental age as either age of death or age at first assessment; the mean parental age (74.2) was given to parents who were alive and less than 16 years older than the genotyped individual, who had a relevant age less than 16, or who did not have a reported age. In this additional analysis we model the effect of age on a linear scale however there seem to exist non-linear trends (Table S31). Future methods could examine whether modeling these non-linear trends increases power.

| Traits | m | $\overline{\text{age}}$ | $c_{age}$ |
|---|---|---|---|
| AD | -2.16 | 68.32 | 0.0542 |
| PD | -2.39 | 68.27 | 0.0266 |
| Lung cancer | -1.82 | 68.27 | -0.0028 |
| Bowel cancer | -1.81 | 68.27 | 0.0105 |
| Stroke | -1.39 | 68.32 | 0.0262 |
| COPD | -1.61 | 68.32 | 0.0085 |
| Prostate cancer | -1.7 | 67.44 | 0.0305 |
| T2D | -1.56 | 68.33 | 0.0178 |
| Breast cancer | -1.47 | 69.03 | -1e-04 |
| Depression | -1.57 | 68.27 | -0.006 |
| CAD | -0.94 | 68.32 | 0.0134 |
| HTN | -0.65 | 68.32 | 0.0093 |

Table S31: **Prevalence of 12 UK Biobank diseases by age.** We report the disease prevalence for each 5 year age bin (<45,45-50,...,85-90,90+), computed by combining genotyped individuals' and their parents' case-status; we merged consecutive bins until each bin contained at least 100 cases. Age in bin refers to the average age within each age bin.

| Age in bin | Prev. in bin | Age in bin | Prev. in bin | Age in bin | Prev. in bin | Age in bin | Prev. in bin |
|---|---|---|---|---|---|---|---|
| AD | | PD | | Lung Cancer | | Bowel Cancer | |
| 48.124 | 0.001 | 44.967 | 0.001 | 41.779 | 0.021 | 41.779 | 0.015 |
| 58.171 | 0.002 | 53.072 | 0.002 | 48.107 | 0.024 | 48.107 | 0.016 |
| 63.051 | 0.006 | 58.171 | 0.003 | 53.072 | 0.036 | 53.072 | 0.024 |
| 67.945 | 0.013 | 63.050 | 0.006 | 58.171 | 0.045 | 58.171 | 0.031 |
| 73.190 | 0.038 | 67.943 | 0.011 | 63.050 | 0.049 | 63.050 | 0.035 |
| 78.063 | 0.066 | 73.186 | 0.020 | 67.943 | 0.058 | 67.943 | 0.044 |
| 82.990 | 0.109 | 78.061 | 0.028 | 73.186 | 0.071 | 73.186 | 0.053 |
| 87.701 | 0.145 | 82.990 | 0.031 | 78.061 | 0.052 | 78.061 | 0.053 |
| 93.379 | 0.153 | 87.701 | 0.027 | 82.990 | 0.034 | 82.990 | 0.050 |
| | | 93.381 | 0.018 | 87.701 | 0.020 | 87.701 | 0.048 |
| | | | | 93.381 | 0.010 | 93.381 | 0.041 |
| Stroke | | COPD | | Prostate Cancer | | T2D | |
| 41.766 | 0.016 | 41.766 | 0.019 | 44.760 | 0.004 | 41.764 | 0.013 |
| 48.107 | 0.023 | 48.107 | 0.026 | 53.082 | 0.013 | 48.107 | 0.020 |
| 53.073 | 0.035 | 53.073 | 0.038 | 58.150 | 0.027 | 53.073 | 0.032 |
| 58.171 | 0.047 | 58.171 | 0.054 | 63.092 | 0.049 | 58.170 | 0.045 |
| 63.051 | 0.063 | 63.051 | 0.068 | 67.933 | 0.065 | 63.051 | 0.063 |
| 67.945 | 0.091 | 67.945 | 0.084 | 73.174 | 0.072 | 67.946 | 0.083 |
| 73.190 | 0.136 | 73.190 | 0.094 | 78.024 | 0.096 | 73.190 | 0.107 |
| 78.063 | 0.156 | 78.063 | 0.088 | 82.957 | 0.114 | 78.063 | 0.111 |
| 82.990 | 0.175 | 82.990 | 0.075 | 87.649 | 0.131 | 82.990 | 0.107 |
| 87.701 | 0.182 | 87.701 | 0.059 | 93.170 | 0.123 | 87.701 | 0.093 |
| 93.379 | 0.167 | 93.379 | 0.037 | | | 93.379 | 0.073 |
| Breast Cancer | | Depression | | CAD | | HTN | |
| 42.072 | 0.055 | 41.779 | 0.072 | 41.766 | 0.066 | 41.766 | 0.110 |
| 48.102 | 0.063 | 48.107 | 0.078 | 48.107 | 0.088 | 48.107 | 0.167 |
| 53.064 | 0.074 | 53.072 | 0.075 | 53.073 | 0.124 | 53.073 | 0.225 |
| 58.192 | 0.084 | 58.171 | 0.070 | 58.171 | 0.159 | 58.171 | 0.281 |
| 63.007 | 0.087 | 63.050 | 0.061 | 63.051 | 0.188 | 63.051 | 0.337 |
| 67.954 | 0.087 | 67.943 | 0.054 | 67.945 | 0.227 | 67.945 | 0.345 |
| 73.198 | 0.077 | 73.186 | 0.054 | 73.190 | 0.250 | 73.190 | 0.269 |
| 78.095 | 0.069 | 78.061 | 0.054 | 78.063 | 0.249 | 78.063 | 0.297 |
| 83.015 | 0.067 | 82.990 | 0.054 | 82.990 | 0.238 | 82.990 | 0.307 |
| 87.733 | 0.065 | 87.701 | 0.049 | 87.701 | 0.223 | 87.701 | 0.301 |
| 93.480 | 0.062 | 93.381 | 0.038 | 93.379 | 0.175 | 93.379 | 0.245 |

Table S32: **Impact of modifying the LT-FH method to incorporate age information for 12 UK Biobank diseases.** We report **(a)** attenuation ratios and **(b)** number of independent loci across the 12 diseases analyzed for each of GWAS, GWAX, LT-FH$_{no-sib}$, LT-PA$_{no-sib}$, LT-PA$_{no-sib,age}$ and LT-FH. PA denotes the use of the Pearson-Aitken formula[23–25] to approximate $E[\epsilon_{o,g}|\cdot]$, implemented for computational reasons in LT-PA$no-sib, age$. To verify that our results were not affected by this approximation, we additionally considered LT-PA$_{no-sib}$. The average phenotypic correlation between LT-FH$_{no-sib}$ and LT-PA$_{no-sib}$ was 0.9995, and association results were very similar. Standard errors are reported in parentheses.

| Trait | GWAS | GWAX | LT-FH$_{no-sib}$ | LT-PA$_{no-sib}$ | LT-PA$_{no-sib,age}$ | LT-FH |
|---|---|---|---|---|---|---|
| **(a)** | | | Attenuation Ratio | | | |
| AD | 0.420 (0.884) | 0.408 (0.188) | 0.395 (0.179) | 0.396 (0.179) | 0.417 (0.211) | 0.381 (0.190) |
| PD | 0.810 (0.454) | 0.254 (0.202) | 0.334 (0.189) | 0.333 (0.189) | 0.322 (0.195) | 0.316 (0.172) |
| Lung cancer | 0.000 (0.298) | 0.104 (0.060) | 0.057 (0.075) | 0.057 (0.075) | 0.047 (0.080) | 0.069 (0.068) |
| Bowel cancer | 0.160 (0.146) | 0.149 (0.081) | 0.185 (0.075) | 0.185 (0.075) | 0.188 (0.075) | 0.106 (0.074) |
| Stroke | 0.179 (0.180) | 0.149 (0.077) | 0.116 (0.072) | 0.117 (0.072) | 0.122 (0.060) | 0.139 (0.069) |
| COPD | 0.077 (0.059) | 0.158 (0.036) | 0.112 (0.035) | 0.112 (0.035) | 0.113 (0.035) | 0.106 (0.034) |
| Prostate cancer | 0.140 (0.093) | 0.085 (0.086) | 0.136 (0.078) | 0.136 (0.078) | 0.129 (0.080) | 0.116 (0.075) |
| T2D | 0.123 (0.043) | 0.164 (0.036) | 0.155 (0.040) | 0.156 (0.040) | 0.153 (0.039) | 0.136 (0.038) |
| Breast cancer | 0.117 (0.081) | 0.191 (0.064) | 0.190 (0.056) | 0.192 (0.056) | 0.194 (0.055) | 0.176 (0.053) |
| Depression | 0.034 (0.054) | 0.115 (0.035) | 0.091 (0.041) | 0.091 (0.041) | 0.091 (0.041) | 0.108 (0.037) |
| CAD | 0.047 (0.035) | 0.084 (0.033) | 0.083 (0.022) | 0.083 (0.022) | 0.088 (0.022) | 0.086 (0.021) |
| HTN | 0.095 (0.017) | 0.090 (0.021) | 0.082 (0.016) | 0.083 (0.016) | 0.084 (0.016) | 0.075 (0.016) |
| Flat mean | 0.183 (0.090) | 0.162 (0.028) | 0.161 (0.026) | 0.162 (0.026) | 0.162 (0.028) | 0.151 (0.025) |
| Inv-var. weighted mean | 0.089 (0.013) | 0.105 (0.014) | 0.097 (0.011) | 0.097 (0.011) | 0.098 (0.011) | 0.090 (0.011) |
| **(b)** | | | Number of independent loci | | | |
| AD | 1 | 8 | 13 | 13 | 13 | 11 |
| PD | 1 | 4 | 5 | 5 | 5 | 4 |
| Lung cancer | 0 | 5 | 5 | 5 | 5 | 5 |
| Bowel cancer | 4 | 9 | 14 | 14 | 14 | 17 |
| Stroke | 0 | 3 | 7 | 7 | 9 | 7 |
| COPD | 5 | 14 | 11 | 11 | 11 | 12 |
| Prostate cancer | 28 | 29 | 41 | 40 | 43 | 38 |
| T2D | 57 | 82 | 112 | 112 | 114 | 120 |
| Breast cancer | 28 | 40 | 39 | 39 | 39 | 49 |
| Depression | 1 | 4 | 4 | 4 | 4 | 6 |
| CAD | 35 | 46 | 83 | 83 | 84 | 92 |
| HTN | 263 | 114 | 317 | 317 | 321 | 329 |
| Total | 423 | 358 | 651 | 650 | 662 | 690 |

Table S33: **Impact of allowing different MAF thresholds for each method.** We report the number of independent loci across 12 diseases for GWAS, GWAX and LT-FH in secondary analyses using different MAF thresholds for each method (Table S16). Results were very similar to primary analyses using the same MAF threshold for each method (Table S21).

| Trait | GWAS | GWAX | LT-FH |
|---|---|---|---|
| AD | 1 | 11 | 15 |
| PD | 1 | 3 | 5 |
| Lung cancer | 0 | 6 | 6 |
| Bowel cancer | 4 | 9 | 18 |
| Stroke | 0 | 3 | 7 |
| COPD | 5 | 14 | 12 |
| Prostate cancer | 28 | 30 | 38 |
| T2D | 57 | 82 | 121 |
| Breast cancer | 28 | 40 | 49 |
| Depression | 1 | 4 | 6 |
| CAD | 35 | 46 | 92 |
| HTN | 263 | 114 | 329 |
| Total | 423 | 362 | 698 |

Table S34: **Results of PA formula in analyses of 12 UK Biobank diseases.** Correlation between LT-FH (used Monte-Carlo integration and assumed at least one sibling affected) and LT-PA (used selection theory and assumed exactly one sibling affected) and the number of independent loci discovered using LT-FH and LT-PA . The RIA (all traits) for LT-FH v. LT-PA is 0.018 (0.007) while for HTN alone is 0.031 (0.014).

| Disease | $\rho$ | LT-PA | LT-FH |
|---|---|---|---|
| AD | 0.9997 | 11 | 11 |
| PD | 1.0000 | 4 | 4 |
| LungCancer | 1.0000 | 5 | 5 |
| BowelCancer | 0.9999 | 17 | 17 |
| Stroke | 0.9999 | 7 | 7 |
| COPD | 0.9998 | 12 | 12 |
| ProstateCancer | 0.9999 | 38 | 38 |
| T2D | 0.9996 | 119 | 120 |
| BreastCancer | 0.9999 | 49 | 49 |
| Depression | 0.9994 | 6 | 6 |
| CAD | 0.9992 | 91 | 92 |
| HTN | 0.9946 | 319 | 329 |

Table S35: **Results of GWAS, GWAX and LT-FH in analyses of 12 UK Biobank diseases (restricted to unrelated individuals) using BOLT-LMM.** We report **(a)** the attenuation ratios and difference in ratios between LT-FH and GWAS (standard errors in parentheses); **(b)** number of independent loci; and average $\chi^2$ for **(c)** genome wide-significant SNPs ($p \leq 5 * 10^{-8}$ for at least one method) and **(d)** all SNPs. In (c), we compute weighted averages in which the weight is determined by the number of genome-wide significant SNPs (shown for each disease in parentheses). In (c) and (d), we restrict to SNPs above the MAF threshold for each disease (reported in Table S16).

| **(a)** | Attenuation Ratio | | | | **(b)** | Number of independent loci | | |
|---|---|---|---|---|---|---|---|---|
| Traits | GWAS | GWAX | LT-FH | LT-FH−GWAS | Traits | GWAS | GWAX | LT-FH |
| AD | 0.419 (0.878) | 0.412 (0.188) | 0.376 (0.189) | -0.043 (0.808) | AD | 1 | 7 | 12 |
| PD | 0.798 (0.452) | 0.258 (0.202) | 0.321 (0.172) | -0.477 (0.406) | PD | 1 | 4 | 4 |
| Lung cancer | 0.000 (0.300) | 0.077 (0.062) | 0.054 (0.071) | 0.054 (0.282) | Lung cancer | 0 | 5 | 5 |
| Bowel cancer | 0.162 (0.146) | 0.147 (0.082) | 0.106 (0.074) | -0.056 (0.127) | Bowel cancer | 4 | 9 | 17 |
| Stroke | 0.165 (0.184) | 0.113 (0.080) | 0.124 (0.071) | -0.041 (0.176) | Stroke | 0 | 3 | 8 |
| COPD | 0.053 (0.061) | 0.128 (0.039) | 0.086 (0.035) | 0.033 (0.049) | COPD | 5 | 14 | 11 |
| Prostate cancer | 0.154 (0.092) | 0.075 (0.086) | 0.119 (0.074) | -0.035 (0.056) | Prostate cancer | 28 | 28 | 39 |
| T2D | 0.118 (0.043) | 0.139 (0.036) | 0.123 (0.038) | 0.005 (0.021) | T2D | 57 | 83 | 121 |
| Breast cancer | 0.120 (0.081) | 0.195 (0.064) | 0.172 (0.054) | 0.051 (0.058) | Breast cancer | 29 | 39 | 50 |
| Depression | 0.032 (0.054) | 0.106 (0.035) | 0.093 (0.037) | 0.060 (0.039) | Depression | 1 | 5 | 7 |
| CAD | 0.027 (0.036) | 0.063 (0.034) | 0.073 (0.022) | 0.046 (0.028) | CAD | 35 | 50 | 92 |
| HTN | 0.088 (0.017) | 0.068 (0.021) | 0.072 (0.016) | -0.017 (0.007) | HTN | 281 | 110 | 370 |
| Flat mean | 0.178 (0.089) | 0.148 (0.028) | 0.143 (0.026) | -0.035 (0.082) | Total | 442 | 357 | 736 |
| Inv-var. weighted mean | 0.082 (0.013) | 0.084 (0.014) | 0.083 (0.011) | 0.001 (0.007) | | | | |

| **(c)** | Mean $\chi^2$ over all genome-significant SNPs* | | | **(d)** | Mean $\chi^2$ over all tested SNPs† | | |
|---|---|---|---|---|---|---|---|
| Traits | GWAS | GWAX | LT-FH | Traits | GWAS | GWAX | LT-FH |
| AD (557) | 22.89 | 204.99 | 222.64 | AD | 1.01 | 1.11 | 1.12 |
| PD (2519) | 9.30 | 42.33 | 45.95 | PD | 1.02 | 1.07 | 1.08 |
| Lung cancer (608) | 9.68 | 58.88 | 63.18 | Lung cancer | 1.03 | 1.20 | 1.18 |
| Bowel cancer (530) | 21.26 | 36.55 | 45.13 | Bowel cancer | 1.04 | 1.09 | 1.10 |
| Stroke (376) | 11.27 | 35.12 | 41.19 | Stroke | 1.04 | 1.09 | 1.10 |
| COPD (910) | 19.74 | 47.14 | 49.94 | COPD | 1.13 | 1.22 | 1.24 |
| Prostate cancer (2863) | 40.84 | 40.06 | 56.27 | Prostate cancer | 1.09 | 1.10 | 1.14 |
| T2D (8071) | 30.99 | 45.37 | 59.70 | T2D | 1.29 | 1.38 | 1.51 |
| Breast cancer (3812) | 32.75 | 44.99 | 54.65 | Breast cancer | 1.06 | 1.10 | 1.12 |
| Depression (299) | 24.06 | 40.41 | 39.61 | Depression | 1.09 | 1.14 | 1.15 |

| | | | | | | | | |
|---|---|---|---|---|---|---|---|---|
| CAD (5726) | 29.17 | 35.73 | 57.36 | | CAD | 1.16 | 1.18 | 1.28 |
| HTN (29663) | 40.04 | 24.81 | 48.02 | | HTN | 1.58 | 1.34 | 1.66 |
| Average | 34.49 | 34.63 | 53.26 | | Average | 1.13 | 1.17 | 1.22 |

Table S36: **Results of GWAS, GWAX and LT-FH in analyses of 12 UK Biobank diseases (including related individuals) using BOLT-LMM.** We report **(a)** attenuation ratios and difference in ratios between LT-FH and GWAS (standard errors in parentheses) and **(b)** number of independent loci. *The inverse-variance weighted mean difference was significantly greater than 0 ($p < 0.001$; two-tailed z-test).

| **(a)** | Attenuation Ratio | | | | **(b)** | Number of independent loci | | |
|---|---|---|---|---|---|---|---|---|
| Traits | GWAS | GWAX | LT-FH | LT-FH−GWAS | Traits | GWAS | GWAX | LT-FH |
| AD | 0.090 (1.070) | 0.520 (0.117) | 0.450 (0.124) | 0.360 (1.010) | AD | 1 | 12 | 14 |
| PD | 0.724 (0.379) | 0.487 (0.094) | 0.483 (0.095) | -0.240 (0.345) | PD | 1 | 5 | 6 |
| Lung cancer | 0.000 (0.275) | 0.248 (0.047) | 0.210 (0.053) | 0.210 (0.264) | Lung cancer | 2 | 7 | 9 |
| Bowel cancer | 0.121 (0.125) | 0.324 (0.064) | 0.252 (0.061) | 0.131 (0.109) | Bowel cancer | 5 | 14 | 25 |
| Stroke | 0.057 (0.142) | 0.214 (0.058) | 0.185 (0.053) | 0.127 (0.130) | Stroke | 1 | 5 | 11 |
| COPD | 0.055 (0.050) | 0.197 (0.030) | 0.125 (0.028) | 0.070 (0.040) | COPD | 5 | 16 | 21 |
| Prostate cancer | 0.163 (0.078) | 0.282 (0.063) | 0.218 (0.060) | 0.055 (0.048) | Prostate cancer | 37 | 33 | 51 |
| T2D | 0.122 (0.041) | 0.192 (0.032) | 0.160 (0.035) | 0.037 (0.018) | T2D | 76 | 112 | 158 |
| Breast cancer | 0.119 (0.065) | 0.335 (0.046) | 0.273 (0.042) | 0.154 (0.047) | Breast cancer | 36 | 46 | 64 |
| Depression | 0.078 (0.045) | 0.160 (0.029) | 0.152 (0.029) | 0.074 (0.034) | Depression | 4 | 12 | 13 |
| CAD | 0.053 (0.031) | 0.111 (0.027) | 0.093 (0.020) | 0.040 (0.024) | CAD | 46 | 63 | 121 |
| HTN | 0.091 (0.015) | 0.095 (0.018) | 0.084 (0.015) | -0.007 (0.007) | HTN | 376 | 155 | 472 |
| Flat mean | 0.139 (0.099) | 0.264 (0.017) | 0.224 (0.017) | 0.084 (0.093) | Total | 590 | 480 | 965 |
| Inv-var. weighted mean | 0.088 (0.012) | 0.131 (0.012) | 0.110 (0.010) | 0.022 (0.007)* | | | | |

Table S37: **Results of GWAS, GWAX and LT-FH with modification for related individuals in analyses of 12 UK Biobank diseases (including related individuals) using BOLT-LMM.** We report results for GWAS, GWAX restricted to unrelated individuals, and LT-FH modified to use only case-control status for all sibling pairs and parent-offspring pairs within the set of target samples. We report **(a)** the attenuation ratios and difference in ratios between LT-FH and GWAS (standard errors in parentheses); **(b)** number of independent loci; and average $\chi^2$ for **(c)** genome wide-significant SNPs ($p \leq 5 * 10^{-8}$ for at least one method) and **(d)** all SNPs. In (c), we compute weighted averages in which the weight is determined by the number of genome-wide significant SNPs (shown for each disease in parentheses). In (c) and (d), we restrict to SNPs above the MAF threshold for each disease (reported in Table S16). We note that the +43% (s.e. 4%) increase in power for LT-FH vs. the trait-specific maximum of GWAS and GWAX is larger than the corresponding +38% increase in power in BOLT-LMM analyses of unrelated individuals (Table S35), because the relative power of GWAX is reduced by restricting to unrelated individuals.

| **(a)** | Attenuation Ratio | | | |
|---|---|---|---|---|
| Traits | GWAS | GWAX | LT-FH | LT-FH−GWAS |
| AD | 0.090 (1.070) | 0.412 (0.188) | 0.356 (0.202) | 0.266 (0.980) |
| PD | 0.724 (0.379) | 0.258 (0.202) | 0.261 (0.153) | -0.462 (0.335) |
| Lung cancer | 0.000 (0.275) | 0.077 (0.062) | 0.093 (0.063) | 0.093 (0.259) |
| Bowel cancer | 0.121 (0.125) | 0.147 (0.082) | 0.113 (0.074) | -0.008 (0.108) |
| Stroke | 0.057 (0.142) | 0.113 (0.080) | 0.094 (0.061) | 0.036 (0.132) |
| COPD | 0.055 (0.050) | 0.128 (0.039) | 0.076 (0.030) | 0.021 (0.039) |
| Prostate cancer | 0.163 (0.078) | 0.075 (0.086) | 0.134 (0.068) | -0.029 (0.045) |
| T2D | 0.122 (0.041) | 0.139 (0.036) | 0.118 (0.038) | -0.004 (0.017) |
| Breast cancer | 0.119 (0.065) | 0.195 (0.064) | 0.176 (0.051) | 0.056 (0.045) |
| Depression | 0.078 (0.045) | 0.106 (0.035) | 0.120 (0.031) | 0.042 (0.032) |
| CAD | 0.053 (0.031) | 0.063 (0.034) | 0.075 (0.021) | 0.022 (0.024) |
| HTN | 0.091 (0.015) | 0.068 (0.021) | 0.075 (0.015) | -0.016 (0.006) |
| Flat mean | 0.139 (0.099) | 0.148 (0.028) | 0.141 (0.025) | 0.002 (0.090) |
| Inv-var. weighted mean | 0.088 (0.012) | 0.085 (0.014) | 0.087 (0.010) | -0.001 (0.006) |

| **(b)** | Number of independent loci | | |
|---|---|---|---|
| Traits | GWAS | GWAX | LT-FH |
| AD | 1 | 7 | 12 |
| PD | 1 | 4 | 5 |
| Lung cancer | 2 | 5 | 9 |
| Bowel cancer | 5 | 9 | 24 |
| Stroke | 1 | 3 | 8 |
| COPD | 5 | 14 | 21 |
| Prostate cancer | 37 | 28 | 50 |
| T2D | 76 | 83 | 148 |
| Breast cancer | 36 | 39 | 57 |
| Depression | 4 | 5 | 11 |
| CAD | 46 | 50 | 110 |
| HTN | 376 | 110 | 453 |
| Total | 590 | 357 | 908 |

| **(c)** | Mean $\chi^2$ over all genome-significant SNPs* | | |
|---|---|---|---|
| Traits | GWAS | GWAX | LT-FH |
| AD (600) | 22.58 | 192.00 | 220.20 |
| PD (2567) | 11.71 | 41.96 | 53.15 |
| Lung cancer (653) | 12.07 | 56.18 | 69.05 |
| Bowel cancer (691) | 22.95 | 31.74 | 44.74 |
| Stroke (409) | 12.82 | 33.44 | 41.01 |
| COPD (1300) | 22.41 | 38.53 | 48.04 |

| **(d)** | Mean $\chi^2$ over all tested SNPs† | | |
|---|---|---|---|
| Traits | GWAS | GWAX | LT-FH |
| AD | 1.01 | 1.11 | 1.13 |
| PD | 1.03 | 1.07 | 1.09 |
| Lung cancer | 1.04 | 1.20 | 1.20 |
| Bowel cancer | 1.06 | 1.09 | 1.11 |
| Stroke | 1.05 | 1.09 | 1.12 |
| COPD | 1.16 | 1.22 | 1.28 |

| | | | | | | | |
|---|---|---|---|---|---|---|---|
| Prostate cancer (3480) | 43.37 | 36.13 | 58.55 | Prostate cancer | 1.12 | 1.10 | 1.16 |
| T2D (11268) | 31.50 | 37.18 | 56.27 | T2D | 1.34 | 1.38 | 1.58 |
| Breast cancer (4803) | 35.62 | 39.84 | 56.15 | Breast cancer | 1.08 | 1.10 | 1.14 |
| Depression (916) | 25.60 | 28.54 | 36.50 | Depression | 1.11 | 1.14 | 1.17 |
| CAD (6961) | 32.11 | 31.73 | 59.11 | CAD | 1.20 | 1.18 | 1.32 |
| HTN (40669) | 41.74 | 20.99 | 48.35 | HTN | 1.70 | 1.34 | 1.77 |
| Average | 36.64 | 29.36 | 53.05 | Average | 1.16 | 1.17 | 1.26 |

Table S38: **Results of GWAS and LT-FH with alternative modification for related individuals in analyses of 12 UK Biobank diseases (including related individuals) using BOLT-LMM.** We report results for GWAS and LT-FH modified to incorporate family history information for exactly one sibling for each set of siblings within the set of target samples (with no filter on family history information for parent-offspring pairs) (LT-FH$_{alt}$). We report **(a)** attenuation ratios and difference in ratios between LT-FH$_{alt}$ and either GWAS or the recommended LT-FH (Table S37); and **(b)** number of independent loci. *The inverse-variance weighted mean difference was significantly greater than 0 ($p < 10^{-6}$; two-tailed z-test).

| | Attenuation Ratio | | | | Number of independent loci | | |
|---|---|---|---|---|---|---|---|
| Traits | GWAS | LT-FH$_{alt}$ | LT-FH$_{alt}$−GWAS | LT-FH$_{alt}$−LT-FH | Traits | GWAS | LT-FH$_{alt}$ |
| AD | 0.090 (1.070) | 0.290 (0.177) | 0.200 (0.991) | -0.066 (0.039) | AD | 1 | 12 |
| PD | 0.724 (0.379) | 0.318 (0.141) | -0.406 (0.335) | 0.057 (0.039) | PD | 1 | 5 |
| Lung cancer | 0.000 (0.275) | 0.086 (0.062) | 0.086 (0.261) | -0.007 (0.014) | Lung cancer | 2 | 9 |
| Bowel cancer | 0.121 (0.125) | 0.130 (0.069) | 0.009 (0.110) | 0.017 (0.017) | Bowel cancer | 5 | 22 |
| Stroke | 0.057 (0.142) | 0.092 (0.060) | 0.035 (0.131) | -0.002 (0.015) | Stroke | 1 | 8 |
| COPD | 0.055 (0.050) | 0.086 (0.028) | 0.031 (0.040) | 0.009 (0.006) | COPD | 5 | 22 |
| Prostate cancer | 0.163 (0.078) | 0.159 (0.065) | -0.004 (0.046) | 0.025 (0.010) | Prostate cancer | 37 | 52 |
| T2D | 0.122 (0.041) | 0.136 (0.036) | 0.014 (0.017) | 0.018 (0.004) | T2D | 76 | 154 |
| Breast cancer | 0.119 (0.065) | 0.212 (0.047) | 0.093 (0.046) | 0.037 (0.009) | Breast cancer | 36 | 65 |
| Depression | 0.078 (0.045) | 0.124 (0.031) | 0.046 (0.033) | 0.004 (0.006) | Depression | 4 | 11 |
| CAD | 0.053 (0.031) | 0.080 (0.020) | 0.027 (0.024) | 0.004 (0.004) | CAD | 46 | 117 |
| HTN | 0.091 (0.015) | 0.078 (0.015) | -0.013 (0.006) | 0.003 (0.002) | HTN | 376 | 458 |
| Flat mean | 0.139 (0.099) | 0.149 (0.023) | 0.010 (0.091) | 0.008 (0.005) | Total | 590 | 935 |
| Inv-var. weighted mean | 0.088 (0.012) | 0.094 (0.010) | 0.005 (0.006) | 0.007 (0.001)* | | | |

Table S39: **Results of simulations with different convergence criteria.** We report the results of simulations in which we vary the convergence criteria of the posterior mean genetic liability. Number of individuals (N) is 100K, number of SNPs (M) is 100K when considering parents and offspring only and 1000 when considering parents, offspring, and 2 siblings; $h^2 = 0.5$; disease prevalence is 5%; we assume perfect knowledge of $h^2/K$ when implementing LT-FH; no environmental correlation between parents and offspring; we consider 10 simulation replicates. The standard error of the mean $\chi^2$ for null and causal SNPs and the standard error for the power are reported in parentheses. In simulations with only parents we estimate posterior mean with 1,000,000 samples from a truncated normal for both parents, this results in an estimate of posterior mean genetic liability with a SEM $< 0.01$. We investigate how using the sampling implemented in the released software differs enforcing a SEM less than 0.1, 0.01 (default), or 0.001. The mean correlation in LT-FH phenotype vector when we estimate the posterior mean with SEM<0.01 and SEM<0.001 across 10 simulation replicates is 0.9999994. In simulations with parents and siblings we estimate posterior mean genetic liability ensuring a SEM $< 0.01$ (as in released software). We investigate the impact of enforcing a SEM less than 0.1 or 0.001. We find mean correlation in LT-FH phenotype vector when we estimate the posterior mean with SEM<0.01 and SEM<0.001 across 10 simulation replicates is 0.9999992.

| | Case-control and parental history | | | |
|---|---|---|---|---|
| | LT-FH | SEM $<$ 0.1 | SEM $<$ 0.01 | SEM $<$ 0.001 |
| $\chi^2_{causal}$ (SEM) | 33.2 (0.169) | 33.2 (0.169) | 33.2 (0.169) | 33.2 (0.169) |
| Power | 0.58 (0.007) | 0.58 (0.007) | 0.58 (0.007) | 0.58 (0.007) |
| $\chi^2_{null}$ (SEM) | 1.001 (0.001) | 1.001 (0.001) | 1.001 (0.001) | 1.001 (0.001) |
| | Case-control, parental, and sibling ($n_s = 2$) history | | | |
| | LT-FH (SEM $<$ 0.01) | SEM $<$ 0.10 | SEM $<$ 0.001 | |
| $\chi^2_{causal}$ (SEM) | 39 (0.184) | 39 (0.184) | 39 (0.184) | |
| Power | 0.75 (0.006) | 0.75 (0.006) | 0.75 (0.006) | |
| $\chi^2_{null}$ (SEM) | 0.964 (0.019) | 0.964 (0.019) | 0.964 (0.019) | |

Table S40: **Concordance between GWAS and LT-FH BOLT-LMM-inf effect sizes.** We report the correlation ($\rho$) between genome-wide significant (GWS) effect sizes for GWAS and LT-FH BOLT-LMM-inf applied to all unrelated Europeans (SNPs above given MAF threshold). The standard error of $\rho$ is estimated as $\sqrt{\frac{1-\rho^2}{n-2}}$. GWS is defined as P$\leq 5*10^{-8}$ for both GWAS and LT-FH BOLT-LMM-inf. For traits for which 0 SNPs are genome-wide significant for both GWAS and LT-FH BOLT-LMM-inf, NA is reported for both $\rho$ and se($\rho$). We note that BOLT-LMM only outputs effect size estimates for BOLT-LMM-inf, the BOLT-LMM approximation to the infinitesimal mixed model. We report a weighted average of $\rho$ across traits weighting by the number of significant SNPs.

| disease | # SNPs | $\rho$ | se($\rho$) |
|---|---|---|---|
| AD | 57 | 0.999 | 0.007 |
| PD | 16 | 0.816 | 0.155 |
| LungCancer | 0 | NA | NA |
| BowelCancer | 85 | 0.994 | 0.012 |
| Stroke | 0 | NA | NA |
| COPD | 207 | 0.998 | 0.004 |
| ProstateCancer | 1521 | 0.995 | 0.002 |
| T2D | 2265 | 0.996 | 0.002 |
| BreastCancer | 1538 | 0.998 | 0.002 |
| Depression | 90 | 0.854 | 0.056 |
| CAD | 1631 | 0.995 | 0.003 |
| HTN | 18080 | 0.997 | 0.001 |
| Weighted Average | | 0.996 | |

Table S41: **Relative effective sample sizes of LT-FH vs. GWAS using linear regression, and LT-FH using BOLT-LMM-inf vs. GWAS using linear regression, in analyses of 12 UK Biobank diseases (restricted to unrelated individuals).** We estimate the relative effective sample size achieved by LT-FH by measuring the boosts in $\chi^2$ (linear regression or BOLT-LMM-inf) association statistics of LT-FH versus $\chi^2$ linear regression GWAS on unrelated European samples (as outlined in Loh et al. 2018 *Nature Genetics* [15]). We compute the relative effective sample size of LT-FH vs. GWAS using linear regression (see Table S22) and LT-FH using BOLT-LMM-inf vs. GWAS using linear regression. In more detail, the relative effective sample size of LT-FH versus GWAS is computed as the median ratio of LT-FH $\chi^2$ statistics (computed using either linear regression or BOLT-LMM-inf) to GWAS (computed using linear regression) $\chi^2$ statistics across genotyped SNPs with $\chi^2 \geq 30$ in GWAS applied via BOLT-LMM to all related Europeans. Ratio values are shown only when the number of SNPs used to calculate the ratio is $\geq$ 10, otherwise NA is displayed; the number of SNPs used to calculate the ratio is shown (next to the trait in parentheses).

| | Relative effective sample size of | |
|---|---|---|
| Trait | LT-FH (Linear Regression) vs. GWAS (Linear Regression) | LT-FH (BOLT-LMM-inf) vs. GWAS (Linear Regression) |
| AD (11) | 10.020 | 10.023 |
| PD (1) | NA | NA |
| LungCancer (0) | NA | NA |
| BowelCancer (11) | 1.568 | 1.563 |
| Stroke (0) | NA | NA |
| COPD (39) | 2.513 | 2.669 |
| ProstateCancer (129) | 1.259 | 1.254 |
| T2D (269) | 1.705 | 1.770 |
| BreastCancer (118) | 1.417 | 1.442 |
| Depression (5) | NA | NA |
| CAD (159) | 1.745 | 1.792 |
| HTN (1458) | 1.098 | 1.172 |

Table S42: **Assigning missing phenotypes to individuals with no ICD9/10 codes reduces sample size while increasing disease prevalence.** We report the sample size (N) and disease prevalence (K) for assigning individuals with no ICD9/10 codes as controls (GWAS) and assigning missing phenotypes to individuals with no ICD9/10 codes ($GWAS_{NA}$). Values are based on 381,493 unrelated individuals of European ancestry.

| | GWAS | | $GWAS_{NA}$ | |
|---|---|---|---|---|
| Trait | N | K | N | K |
| AD | 381493 | 0.001 | 349995 | 0.001 |
| PD | 381493 | 0.003 | 349995 | 0.003 |
| Lung cancer | 381493 | 0.006 | 304266 | 0.007 |
| Bowel cancer | 381493 | 0.013 | 304266 | 0.016 |
| Stroke | 381493 | 0.023 | 381459 | 0.023 |
| COPD | 381493 | 0.035 | 381469 | 0.035 |
| Prostate cancer | 175450 | 0.037 | 136639 | 0.048 |
| Breast cancer | 206043 | 0.061 | 167627 | 0.075 |
| Depression | 381493 | 0.073 | 349995 | 0.080 |
| CAD | 381493 | 0.083 | 381459 | 0.083 |
| HTN | 381493 | 0.318 | 381459 | 0.318 |

Table S43: **Assigning missing phenotypes to individuals with no ICD9/10 codes slightly but consistently reduces sample size times observed-scale SNP-heritability and has very little impact on GWAS power.** We report **(a)** $N$, $h_g^2$ and $Nh_g^2$ and (b) number of independent loci for assigning individuals with no ICD9/10 codes as controls (GWAS) and assigning missing phenotypes to individuals with no ICD9/10 codes (GWAS$_{NA}$). Results are based on 381,493 unrelated individuals of European ancestry. Association results are based on linear regression. $h_g^2$ is estimated using S-LDSC[16,26] with the baselineLD model (v1.1) and the standard error of the difference is computed via block jackknife.

**(a)**

| Trait | N | $h_g^2$ | $Nh_g^2$ | $N_{NA}$ | $h_{g,NA}^2$ | $N_{NA}h_{g,NA}^2$ | $N_{NA}h_{g,NA}^2 - N_{Ctrl}h_{g,Ctrl}^2$ |
|---|---|---|---|---|---|---|---|
| AD | 381493 | 5e-04 (0.0021) | 190.75 (785.38) | 349995 | 5e-04 (0.0022) | 175 (781.9) | -15.75 (29.94) |
| PD | 381493 | -6e-04 (0.0018) | -228.9 (704.06) | 349995 | -8e-04 (0.002) | -280 (706.98) | -51.1 (24.04) |
| Lung cancer | 381493 | 0.0053 (0.0018) | 2021.91 (697.05) | 304266 | 0.0065 (0.0023) | 1977.73 (696.12) | -44.18 (48.4) |
| Bowel cancer | 381493 | 0.0083 (0.0022) | 3166.39 (835.07) | 304266 | 0.01 (0.0027) | 3164.37 (830.44) | -2.03 (67.18) |
| Stroke | 381493 | 0.0056 (0.002) | 2136.36 (751.47) | 381459 | 0.0056 (0.002) | 2136.17 (751.47) | -0.19 (4.14) |
| COPD | 381493 | 0.021 (0.0023) | 7973.2 (890.98) | 381469 | 0.021 (0.0023) | 7972.7 (891.3) | -0.5 (5.23) |
| Prostate cancer | 175450 | 0.036 (0.0061) | 6333.74 (1065.88) | 136639 | 0.045 (0.0077) | 6176.08 (1057.88) | -157.66 (112.78) |
| Breast cancer | 206043 | 0.031 (0.0051) | 6304.92 (1054.16) | 167627 | 0.038 (0.0062) | 6286.01 (1043.11) | -18.9 (134.57) |
| Depression | 381493 | 0.023 (0.0023) | 8736.19 (885.56) | 349995 | 0.023 (0.0025) | 8049.89 (866.69) | -686.3 (115.43) |
| CAD | 381493 | 0.043 (0.0034) | 16251.6 (1292.38) | 381459 | 0.043 (0.0034) | 16250.15 (1292.59) | -1.45 (5.68) |
| HTN | 381493 | 0.13 (0.0055) | 49250.75 (2084.89) | 381459 | 0.13 (0.0055) | 49208.21 (2084.9) | -42.54 (12.11) |

**(b)**

| Traits | GWAS | GWAS$_{NA}$ |
|---|---|---|
| AD | 1 | 1 |
| PD | 1 | 1 |
| Lung cancer | 0 | 0 |
| Bowel cancer | 4 | 4 |
| Stroke | 0 | 0 |
| COPD | 5 | 5 |
| Prostate cancer | 28 | 30 |
| Breast cancer | 28 | 28 |
| Depression | 1 | 1 |
| CAD | 35 | 35 |
| HTN | 263 | 262 |
| Total | 366 | 367 |

Table S44: **GWAX prevalence (i.e. prevalence of proxy cases) is generally more than double the parental prevalence and many times larger than the disease prevalence.** We report sample size (N) and disease prevalence (K) for genotyped individuals (GWAS), parents of genotyped individuals, and proxy cases (GWAX). Values are based on 381,493 unrelated individuals of European ancestry.

| | GWAS | | Parents | | GWAX | |
|---|---|---|---|---|---|---|
| Trait | N | K | N | K | N | K |
| AD | 381493 | 0.001 | 705856 | 0.065 | 324512 | 0.141 |
| PD | 381493 | 0.003 | 696882 | 0.020 | 318792 | 0.052 |
| Lung cancer | 381493 | 0.006 | 696882 | 0.064 | 323838 | 0.154 |
| Bowel cancer | 381493 | 0.013 | 696882 | 0.054 | 322887 | 0.144 |
| Stroke | 381493 | 0.023 | 705856 | 0.144 | 332468 | 0.318 |
| COPD | 381493 | 0.035 | 705856 | 0.082 | 329495 | 0.209 |
| Prostate cancer | 175450 | 0.037 | 340547 | 0.075 | 331458 | 0.107 |
| T2D | 380180 | 0.042 | 705856 | 0.092 | 330267 | 0.262 |
| Breast cancer | 206043 | 0.061 | 356335 | 0.082 | 348170 | 0.146 |
| Depression | 381493 | 0.073 | 696882 | 0.052 | 328186 | 0.215 |
| CAD | 381493 | 0.083 | 705856 | 0.255 | 341810 | 0.522 |
| HTN | 381493 | 0.318 | 705856 | 0.260 | 354208 | 0.657 |

Table S45: **Configurations of family history in UK Biobank family history.** We list the 377 possible configurations of case-control status and family history of disease. We note that sibling history is a binary variable (i.e. at least one sibling has the disease).

See Excel file.

Figure S1: **QQ plots from simulations with default parameter settings.** We report quantile-quantile (QQ) plots for null SNPs in simulations with default parameter settings. Results are based on 10 simulation replicates. These QQ plots compare the observed distribution of p-values with the standard uniform distribution. We plot the observed $-\log_{10}(p)$ as a function of $-\log_{10}(\frac{\text{rank}}{n+1})$ and the 95% confidence bands are constructed pointwise using the beta distribution.

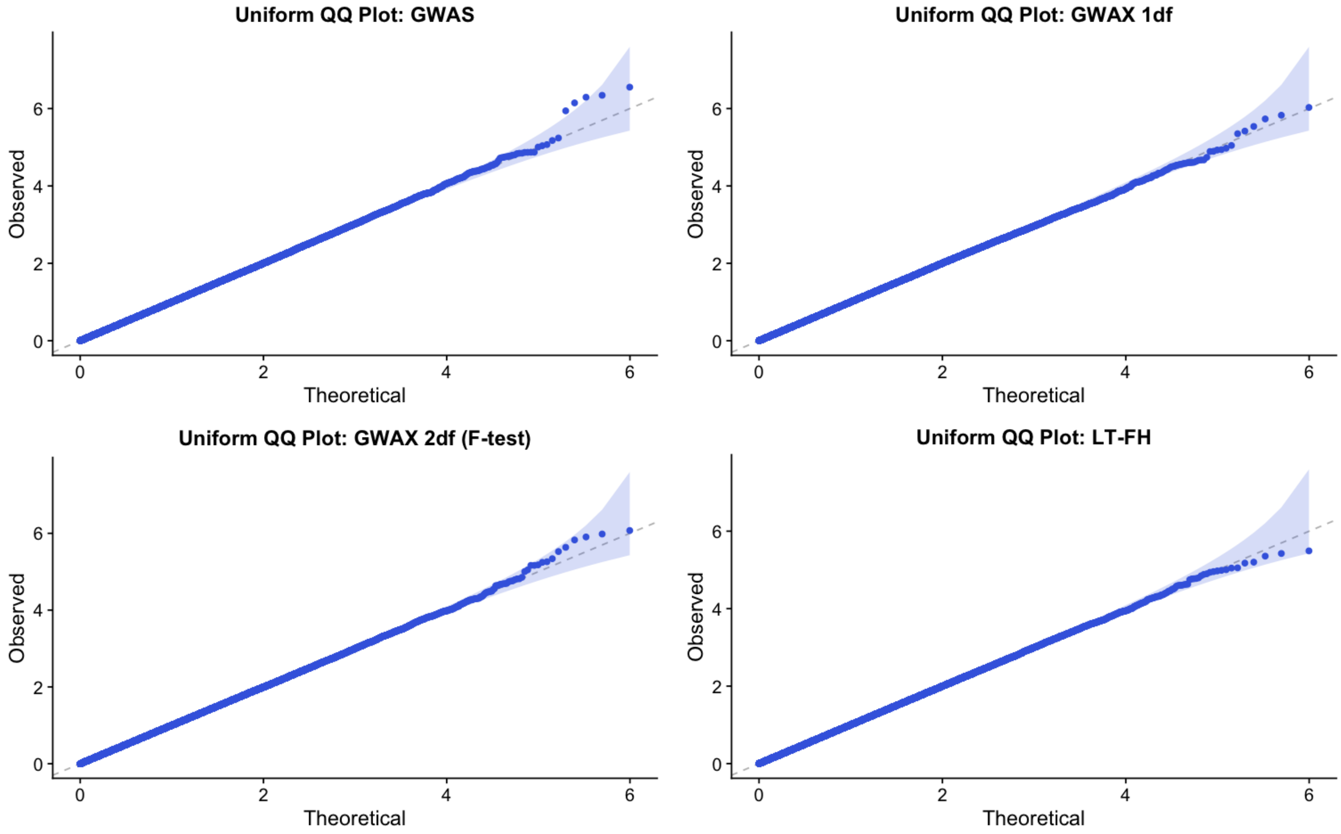

Figure S2: **Distribution of LT-FH phenotypes for 12 UK Biobank diseases.** We plot the distribution of the LT-FH phenotype for each disease. We also report the kurtosis for both GWAS and LT-FH; Pearson's measure of kurtosis, $\kappa = \frac{E[(X-\mu)^4]}{(E[(X-\mu)^2])^2}$, is calculated using the R package moments.

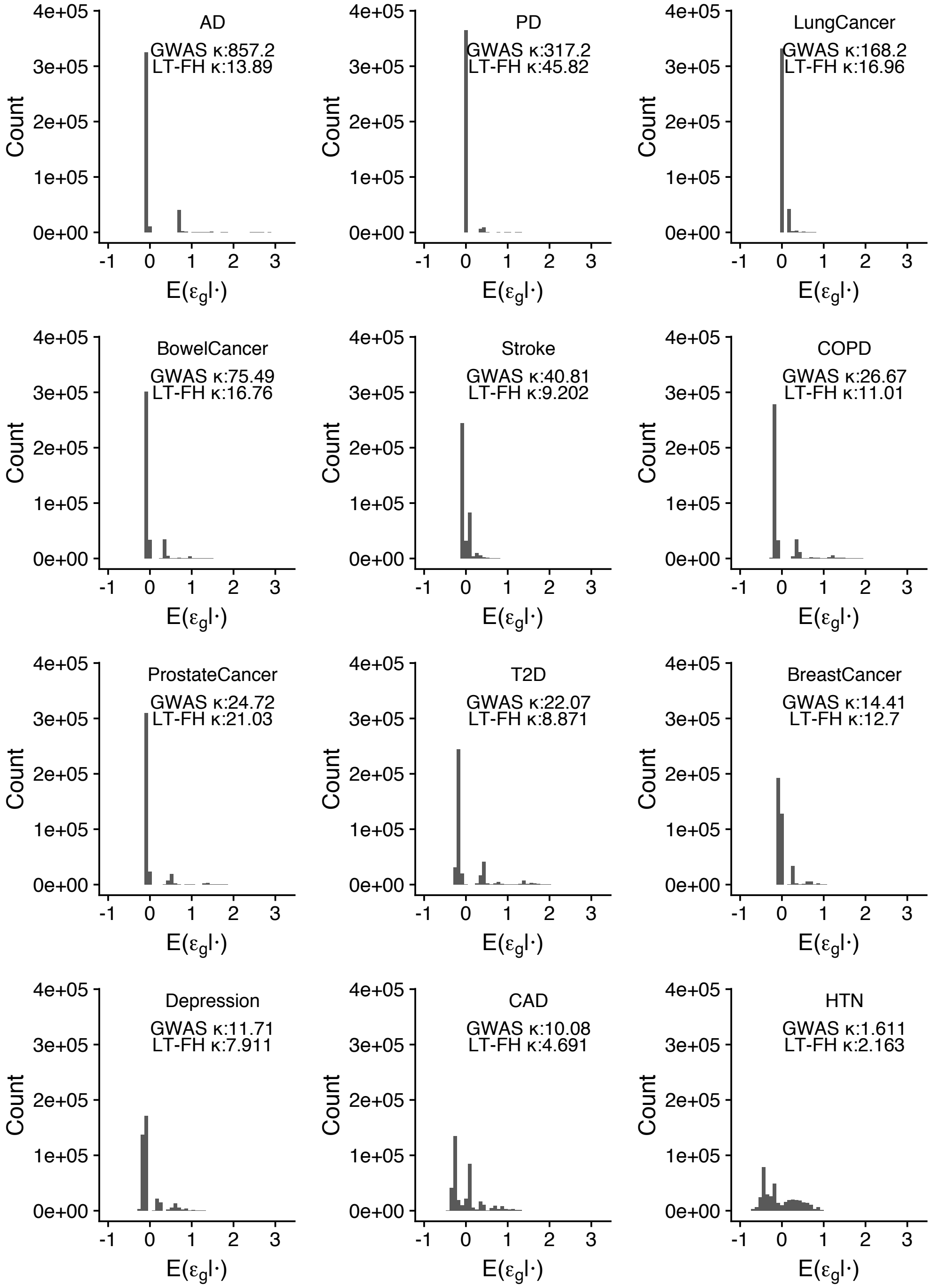

Figure S3: **Impact of modifying the LT-FH method to incorporate age information as a function of the liability threshold model parameter for age for 12 UK Biobank diseases.** We plot the increase in number of independent loci for LT-FH$^{PA}_{no-sib,age}$ relative to LT-FH$^{PA}_{no-sib}$ (Table S32) against the liability threshold model parameter $|c_{age}|$ (Table S30).

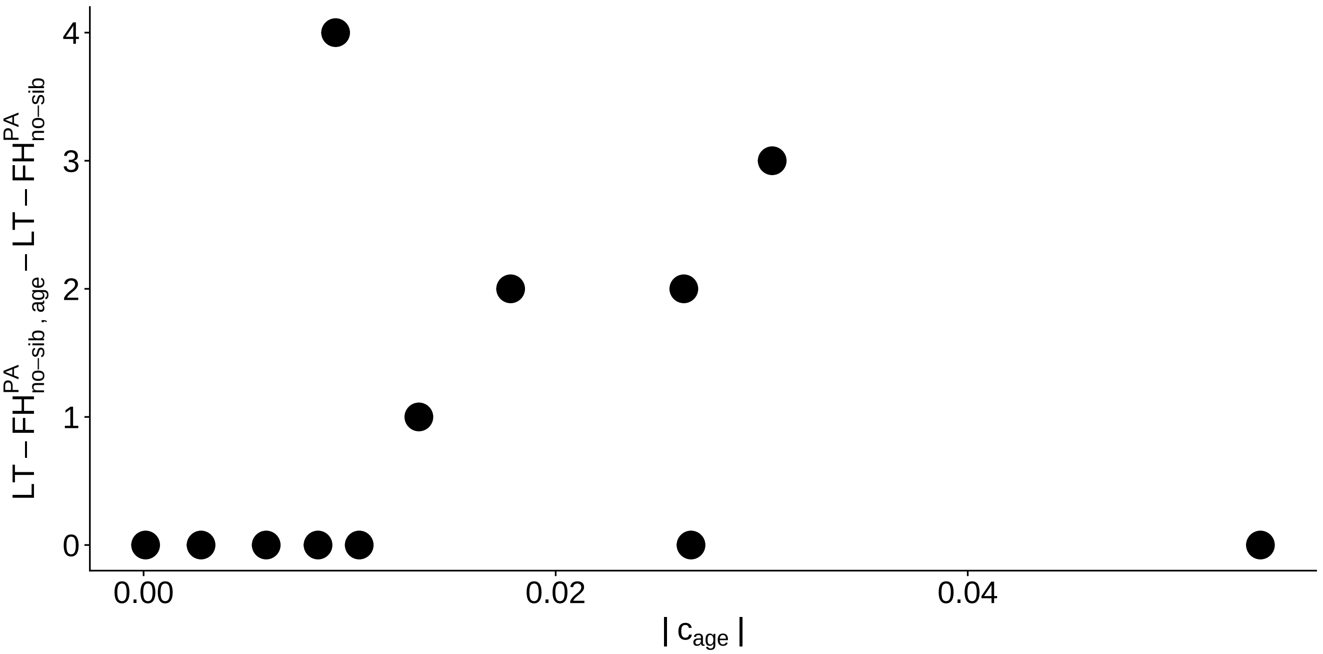

Figure S4: **LT-FH increases association power across 12 diseases from the UK Biobank in analyses incorporating related individuals.** We report results of GWAS using BOLT-LMM on related Europeans, GWAX using BOLT-LMM on unrelated Europeans, and LT-FH using BOLT-LMM on related Europeans using only case-control status for all sibling pairs and parent-offspring pairs within the set of target samples. Numerical results are reported in Table S37.

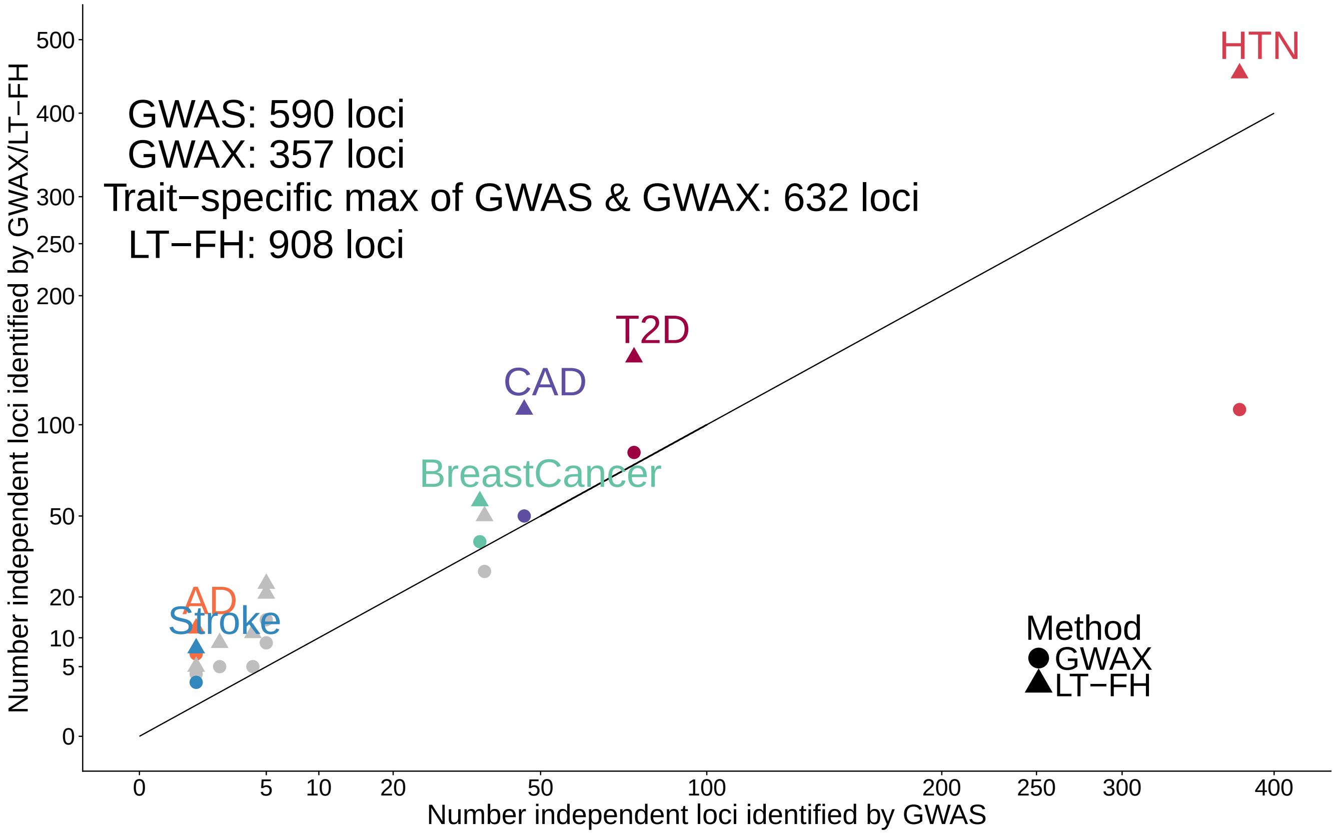

Figure S5: **Strong concordance between GWAS BOLT-LMM-inf effect sizes and transformed LT-FH BOLT-LMM-inf effect sizes.** We plot GWAS BOLT-LMM-inf effect sizes and transformed LT-FH BOLT-LMM-inf effect sizes for genome-wide significant effect sizes ($P \leq 5*10^{-8}$ for both GWAS and LT-FH BOLT-LMM-inf). We note that BOLT-LMM only outputs effect size estimates for BOLT-LMM-inf, the BOLT-LMM approximation to the infinitesimal mixed model. Our effect size for GWAS is the outputted $\beta_{GWAS,BOLT-LMM-inf}$ (per-allele observed scale) and for LT-FH we estimate a (per-allele observed scale) effect size as $\beta = \frac{\beta_{LTFH,BOLT-LMM-inf}}{se(\beta_{LTFH,BOLT-LMM-inf})\sqrt{N_{GWAS}*c}} * \frac{\sqrt{K(1-K)}}{\sqrt{2(MAF)(1-MAF)}}$, where $c$ is the boost in $N_{eff}$ for LT-FH relative to GWAS, $K$ is disease prevalence in GWAS and $MAF$ is the minor allele frequency of the SNP.

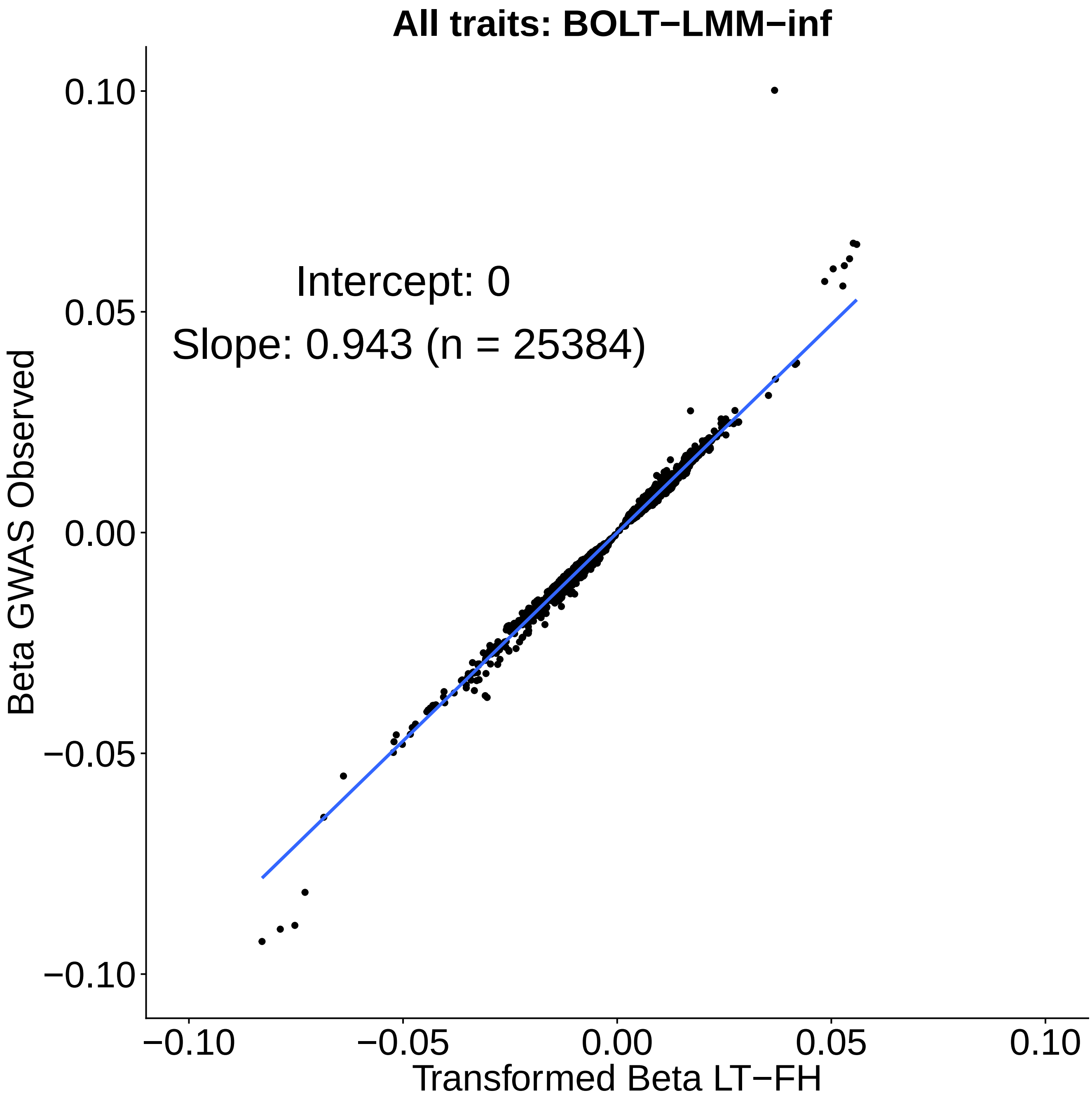
